# Atypical meiosis can be adaptive in outcrossed *S. pombe* due to *wtf* meiotic drivers

**DOI:** 10.1101/2020.04.28.066035

**Authors:** María Angélica Bravo Núñez, Ibrahim M. Sabbarini, Lauren E. Eide, Robert L. Unckless, Sarah E. Zanders

## Abstract

Killer meiotic drivers are genetic parasites that destroy ‘sibling’ gametes lacking the driver allele. The fitness costs of drive can lead to selection of unlinked suppressors. This suppression could involve evolutionary tradeoffs that compromise gametogenesis and contribute to infertility. *Schizosaccharomyces pombe*, an organism containing numerous gamete-killing *wtf* drivers, offers a tractable system to test this hypothesis. Here, we demonstrate that in scenarios analogous to outcrossing, *wtf* drivers generate a fitness landscape in which atypical gametes, such as aneuploids and diploids, are advantageous. In this context, *wtf* drivers can decrease the fitness cost of mutations that disrupt meiotic fidelity and, in some circumstances, can even make such mutations beneficial. Moreover, we find that *S. pombe* isolates vary greatly in their ability to make haploid gametes, with some isolates generating more than 25% aneuploid or diploid gametes. This work empirically demonstrates the potential for meiotic drivers to shape the evolution of gametogenesis.

## Introduction

Parasites are pervasive in biology and can impose extreme fitness costs on their hosts (McLaughlin and Malik 2017; Sorci and Garnier 2008). Due to these fitness effects, there can be strong selection for variants of host genes that can subvert such parasites (Kutzer and Armitage 2016; McLaughlin and Malik 2017; Sorci and Garnier 2008). However, gene variants that promote host defense may be maladapted for other facets of host physiology, leading to evolutionary tradeoffs. For example, the sickle cell trait has been selected in malaria-endemic human populations as it provides heterozygous individuals some protection against the malaria-causing parasite, *Plasmodium falciparum*. However, the advantages of this allele come with a high cost as homozygotes develop sickle cell disease (Elguero et al., 2015; Serjeant 2010).

In addition to external parasites like *P. falciparum*, organisms are also challenged by a variety of parasitic, or ‘selfish’, DNA sequences within their genomes (Burt and Trivers 2006). Meiotic drivers are one type of selfish DNA elements found throughout eukaryotes. Meiotic drive loci exploit meiosis to increase their chances of being passed on to the next generation. Rather than being transmitted to 50% of the progeny of a heterozygote, these selfish loci use a variety of tactics to promote their own transmission into up to 100% of the gametes (Sandler and Novitski 1957; Zimmering et al., 1970). This cheating can impose a variety of fitness costs on the host (Zanders and Unckless 2019). Due to these costs, variants that suppress meiotic drive can be favored by selection (Burt and Trivers 2006; Crow 1991; Hartl 1975). Analogous to the sickle cell trait, this could lead to evolutionary tradeoffs where variants that are suboptimal for some aspect of gametogenesis may be selected due to their ability to mitigate the costs of meiotic drivers.

In this work, we explore the potential selective pressures meiotic drivers can impose on the evolution of gametogenesis. We use the fission yeast *S. pombe* as it is infested with multiple meiotic drive genes belonging to the *wtf* (***<u>w</u>****ith **<u>Tf</u>***) gene family (Bravo Núñez et al., 2020; Eickbush et al., 2019; Hu et al., 2017; Nuckolls et al., 2017). Each *wtf* meiotic driver encodes two proteins from two largely overlapping transcripts with distinct start sites: a poison (Wtf^poison^) and an antidote (Wtf^antidote^) (Hu et al., 2017; Nuckolls et al., 2017). In *wtf*+/*wtf*- heterozygotes, all the developing meiotic products (spores) are exposed to the Wtf^poison^, but only those that inherit the *wtf+* allele express the corresponding Wtf^antidote^ and neutralize the poison. This allows *wtf* drivers to gain a transmission advantage into the next generation by killing the spores that do not inherit the *wtf*+ allele from heterozygous (*wtf+/wtf-*) diploids (Hu et al., 2017; Nuckolls et al., 2017).

The poison and antidote proteins of a given *wtf* meiotic driver share a considerable length of amino acid sequence (>200 residues) (Hu et al., 2017; Nuckolls et al., 2017). This shared amino acid sequence may be important for a Wtf^antidote^ to neutralize a given Wtf^poison^ protein. Strikingly, even a mistmatch of two amino acids within the C-terminus can disrupt the ability of a Wtf^antidote^ to neutralize a Wtf^poison^ (Bravo Núñez et al., 2018). The antidote of a given *wtf* gene generally does not neutralize poisons produced by other *wtf* drivers with distinct sequences (Bravo Núñez et al., 2018; Bravo Núñez et al., 2020; Hu et al., 2017). In diploids heterozygous for *wtf* loci, these drivers can be described as ‘competing’ as they exist on separate haplotypes. When *S. pombe* isolates outcross, multiple *wtf* drivers may be in competition during gametogenesis (Eickbush et al., 2019).

Here, we find that heterozygous, competing *wtf* drivers provide a selective advantage to atypical spores that inherit more than a haploid complement of *wtf* drivers. These spores are selected due to the preferential destruction of haploid spores by the *wtf* driver(s) they do not inherit. The selected atypical spores include aneuploids, diploids, and spores inheriting *wtf* gene duplications resulting from unequal crossovers between homologous chromosomes. We use a combination of empirical analyses and modeling to demonstrate that competing *wtf* drivers generate an environment where variants that disrupt meiotic chromosome segregation can increase fitness. Finally, we show that variants that generate high numbers of atypical meiotic products may be common in *S. pombe* populations. Overall, this work demonstrates the capacity of meiotic drivers to impact the evolution of gametogenesis and suggests meiotic drive could have indirectly contributed to the high frequency of atypical gametes generated by *S. pombe* diploids.

## Results

### The viable spores produced by outcrossed *S. pombe* diploids are frequently aneuploid or diploid

Most *S. pombe* research is conducted on isogenic strains derived from a single isolate called L972 (referred to in this work as *Sp*). Most of our knowledge about *S. pombe* meiosis thus stems from studying homozygous *Sp* diploids. Relatively little is known about the meiotic phenotypes of other isolates or of heterozygotes generated by crossing different haploid isolates (referred to as ‘outcrossing’ here) (Avelar et al., 2013; Hu et al., 2017; Jeffares et al., 2015; Zanders et al., 2014).

There are over 100 additional *S. pombe* isolates that have been sequenced and phenotypically characterized to some extent (Brown et al., 2011; Jeffares et al., 2015; Tusso et al., 2019). All known *S. pombe* isolates share an average DNA sequence identity of >99% for nonrepetitive regions. Despite minimal sequence divergence between the strains, outcrossing often yields diploids that exhibit low fertility (i.e. they produce few viable spores) (Avelar et al., 2013; Hu et al., 2017; Jeffares et al., 2015; Singh and Klar 2002; Zanders et al., 2014). Differences in karyotype (such as chromosomal rearrangements) between isolates and pervasive meiotic drive are the two demonstrated causes of infertility in heterozygous *S. pombe* diploids (Avelar et al., 2013; Hu et al., 2017; Nuckolls et al., 2017; Zanders et al., 2014).

We previously characterized diploids generated by outcrossing *Sp* to another isolate called *S. kambucha* (*Sk*) (Nuckolls et al., 2017; Zanders et al., 2014). Although these *Sp*/*Sk* heterozygotes make few viable spores (5% of *Sp* wild type), 79% of the spores that survive appear to be heterozygous diploids or aneuploids as they inherit centromere 3-linked marker genes from both *Sk* and *Sp* (Figure 1B, diploid 1) (Zanders et al., 2014). We refer to this phenotype as ‘disomy’ and refer to spores with this trait as ‘disomic.’

**Figure 1.**
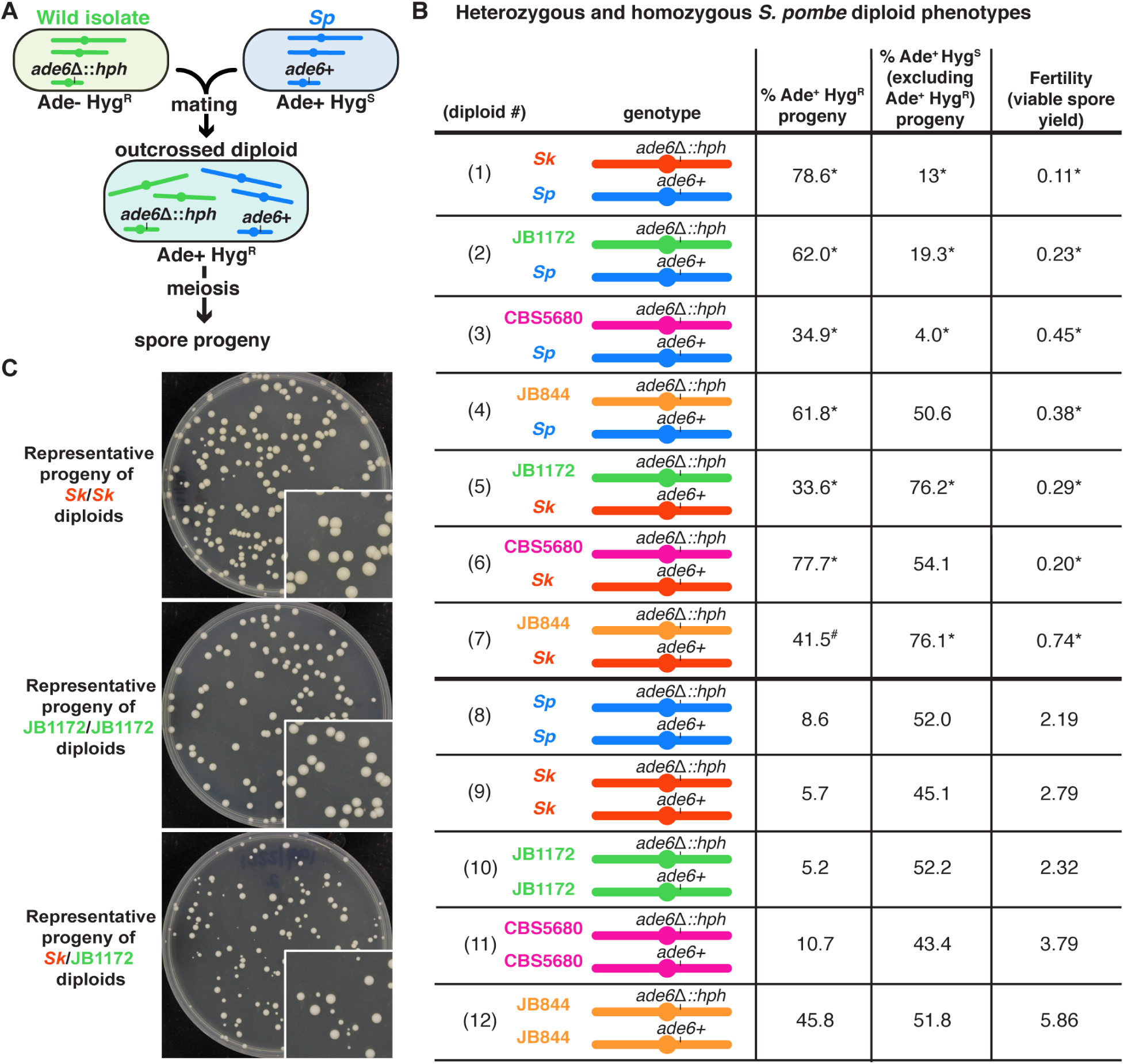
Outcrossed *S. pombe* diploids frequently produce disomic gametes. (A) Schematic of the experimental approach. The *ade6* gene is linked to centromere 3. The karyotypes of JB1172, CBS5680, and JB844 are unknown, but outside of an inversion on *Sp* chromosome 1, *Sp* likely represents the ancestral karyotype (Avelar et al., 2013; Brown et al., 2011). (B) Phenotypes of heterozygous or homozygous *S. pombe* diploids. Allele transmission of chromosome 3 was assayed using co-dominant markers at *ade6* (*ade6+* and *ade6*Δ::*hphMX6*). The *ade6+* allele confers an Ade+ phenotype, while the *ade6*Δ::*hphMX6* provides resistance to Hygromycin B (Hyg^R^). Heterozygous aneuploid or diploid spores are Ade+ Hyg^R^. The phenotypes of each heterozygote were compared to those of homozygous diploids from both parental strain backgrounds. * indicates p-value <0.025 (G-test [Ade+ Hyg^R^ progeny] and Wilcoxon test [fertility]) for the heterozygotes relative to homozygous diploid from both parental backgrounds. Diploid 7 was only significantly different (p-value < 0.025) in the frequency of Ade+ Hyg^R^ progeny when compared to diploid 9, but not when compared to diploid 12. This is indicated with #. To detect biased allele transmission, diploids 2-4 were compared to diploid 8 and diploids 5-7 were compared to diploid 9. * indicates p-value <0.05 (G-test [allele transmission]). More than 200 viable spores were scored for each diploid. Raw data can be found in Figure 1—figure supplement 2 and Figure 1—figure supplement 3. (C) Representative image of the viable spore colonies generated by homozygous *Sk* and JB1172 diploids and heterozygous *Sk/*JB1172 diploids. Images of colonies generated by other diploids are shown in Figure 1—figure supplement 1.

In this study, we tested if the phenotypes observed in *Sp/Sk* heterozygotes are common amongst the viable spores produced by other *S. pombe* diploids generated by outcrossing. To do this, we first inserted a centromere 3-linked genetic marker (*ade6*Δ::*hphMX6*) in three additional natural isolates: JB1172, CBS5680, and JB844. We chose these isolates because out of the seven strains attempted, we were able to transform the desired markers into only those three (see methods). Next, we mated each of these genetically marked natural isolates to *Sp* and *Sk* to generate a series of heterozygous diploids (Figure 1A). Similar to *Sp*/*Sk* heterozygotes, each of these heterozygous diploids had a low viable spore yield (viable spores recovered per diploid cell placed on sporulation media) compared to homozygous diploids (Figure 1B, compare diploids 2-7 to diploids 8-12). These reduced yields are likely due to increased spore death in the heterozygotes (Hu et al., 2017; Jeffares et al., 2015; Singh and Klar 2002; Zanders et al., 2014). The differences in viable spore yield amongst the homozygous strains could be due to numerous factors, including sporulation efficiency, the number of mitotic divisions a diploid completes before undergoing meiosis, and spore viability.

In addition to having low viable spore yields, the heterozygous diploids frequently produced spores that grew into small colonies with irregular shapes, a hallmark of aneuploidy (Figure 1C, Figure 1—figure supplement 1) (Niwa et al., 2006). Consistent with this, 34-78% of the viable spores produced by heterozygotes inherited both parental alleles of *ade6* (*ade6*+ and *ade6*Δ::*hphMX6*) and were thus likely disomic for at least chromosome 3 (Figure 1B, diploids 2-7). The frequency of disomic gametes was generally higher in heterozygotes than homozygotes. There was, however, considerable variation amongst the homozygotes with disomy frequencies ranging from 5-46% disomic gametes (Figure 1B, diploids 8-12).

### Multiple sets of competing meiotic drivers can select for disomic spores by killing haploids

We next wanted to determine why so many of the surviving spores produced by outcrossed *S. pombe* diploids were heterozygous disomes for chromosome 3. We previously proposed a model in which distinct *wtf* meiotic drivers found on competing chromosome 3 haplotypes were killing haploid spores (López Hernández and Zanders 2018). We hypothesized that in the presence of diverged meiotic drivers on opposite haplotypes, haploid spores will inherit one driver and be killed by the driver on the opposite haplotype. However, heterozygous disomic spores will inherit both sets of competing *wtf* drive alleles, which are almost all found on chromosome 3. These disomic spores should thus survive, as they will contain every Wtf^antidote^ necessary to counteract the Wtf poisons (Figure 2A). Consistent with this model, *Sp/Sk* heterozygotes do not make more disomic spores per meiosis than *Sp* or *Sk* homozygotes (Zanders et al., 2014).

**Figure 2.**
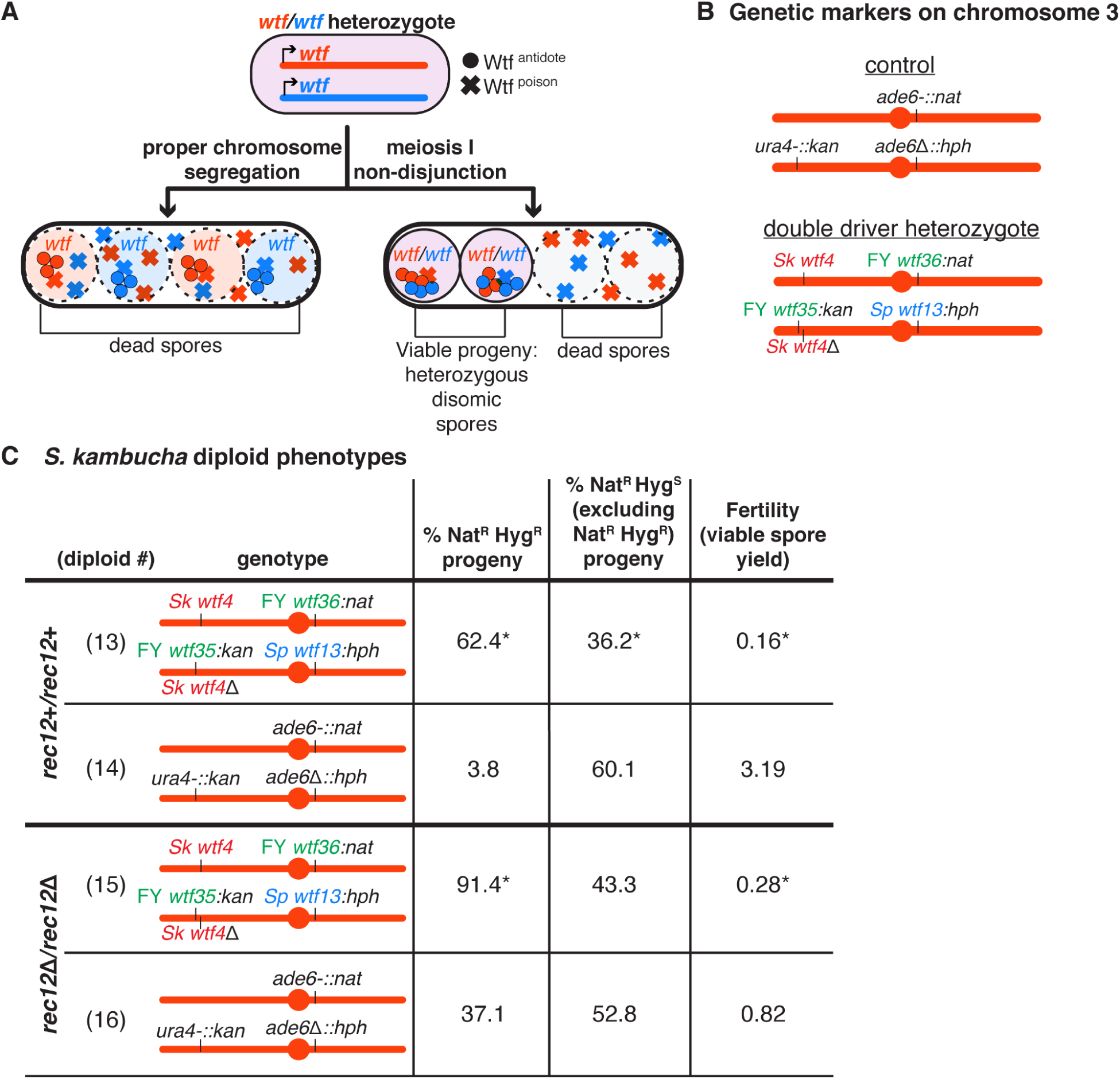
A high fraction of viable gametes are disomic in *Sk* strains with *wtf* competition at two loci. (A) Model for a diploid heterozygous for distinct *wtf* meiotic drivers. Spores are destroyed by any *wtf* driver that they do not inherit from the diploid progenitor cell. Meiosis I chromosome missegregation is one mechanism by which spores can inherit *wtf* alleles on competing haplotypes and survive. (B) Schematic of the genetic markers at *ura4* and *ade6* in the control diploid, and the *wtf* transgenes inserted at *ura4* and *ade6* in *Sp* chromosome 3 in the double driver heterozygote. *wtf* genes from the *Sp, Sk*, and FY29033 strains are depicted in blue, red, and green, respectively. The *wtf* drivers shown here drive when heterozygous and do not counteract the effect of the other drivers (see Figure 2—figure supplement 1). (C) Phenotypes of the double driver heterozygote or control diploid in *rec12*+ (top) and *rec12*Δ (bottom) strain backgrounds. For statistical analyses, the frequency of disomic progeny, allele transmission, and fertility in the double driver heterozygotes were compared to the control diploids. Diploid 13 was compared to control diploid 14, and diploid 15 was compared to control diploid 16. * indicates p-value <0.05 (G-test [allele transmission and Nat^R^ Hyg^R^ progeny] and Wilcoxon test [fertility]). The data for diploid 14 were previously published in Bravo Núñez et al., 2020. Raw data can be found in Figure 2—figure supplement 4 and Figure 2—figure supplement 5.

To test our model, we engineered an *Sk* diploid that is heterozygous for two unlinked sets of competing *wtf* drivers on chromosome 3. We refer to this diploid as the “double driver heterozygote” (Figure 2B). Importantly, all four of the drivers in this strain are functional and cannot fully suppress any of the other three drivers (Figure 2—figure supplement 1) (Bravo Núñez et al., 2018; Bravo Núñez et al., 2020; Nuckolls et al., 2017). Consistent with our hypothesis, we found that 62% of the viable spores generated by the double driver heterozygote inherited both of the parental alleles at *ade6*, compared to 4% of the viable spores generated by the control diploid (Figure 2C, compare diploid 13 to 14). Additionally, many of the viable spores produced by the double driver heterozygote generated small, misshapen colonies characteristic of aneuploids.

Finally, the fertility of the double driver heterozygote was 20-fold lower than the control diploid (Figure 2C, compare diploid 13 to diploid 14). This reduction in fertility is consistent with the destruction of haploid spores that do not inherit every driver. However, our results are also consistent with the alternative hypothesis that meiosis in the double driver heterozygote more frequently produces disomic spores. To distinguish these possibilities, we used the observed frequency of disomes and the viable spore yield to calculate the number of disomes produced per diploid cell placed on sporulation media. We found that the number of disomic progeny produced per cell did not increase between the double driver heterozygote and the control (Figure 2—figure supplement 2, compare diploid 13 to diploid 14), weakening support for the alternative hypothesis.

To test if our results were dependent on strain background, we made an analogous double driver heterozygote in the *Sp* strain background and observed a similar decrease in fertility and increase in disomy specifically amongst the surviving spores (Figure 2—figure supplement 3). These results are consistent with our hypothesis that diploids carrying multiple sets of heterozygous *wtf* meiotic drivers generate heterozygous disomic spores due to the destruction of haploid progeny.

### Heterozygosity at *wtf* loci contributes to the high frequency of disomic spores generated by outcrossed diploids

To further determine the contribution of competing *wtf* drivers to the high level of disomic spores, we decided to test our model in a strain background with more extensive heterozygosity, like those generated by outcrossing. For these experiments, we started with an *Sp/Sk* mosaic diploid strain that is heterozygous for eight known or predicted *wtf* meiotic drivers (Bravo Núñez et al., 2020; Eickbush et al., 2019). This mosaic diploid is homozygous for *Sk* chromosomes 1 and 2 but is heterozygous for most of chromosome 3 for *Sp* and *Sk*-derived sequences (Figure 3A). These diploids also lack *rec12*, a gene which encodes the endonuclease that initiates meiotic recombination by generating double-strand DNA breaks (DSBs) (Bergerat et al., 1997; Keeney et al., 1997). The lack of induced recombination in these diploids results in competition between the *Sp* and *Sk wtf* drivers on chromosome 3, as haploid spores will generally inherit either every *Sp* driver or every *Sk* driver. To determine the transmission of *Sp-* and *Sk-*derived sequences on chromosome 3, we genotyped the centromere-linked *ade6* locus.

**Figure 3.**
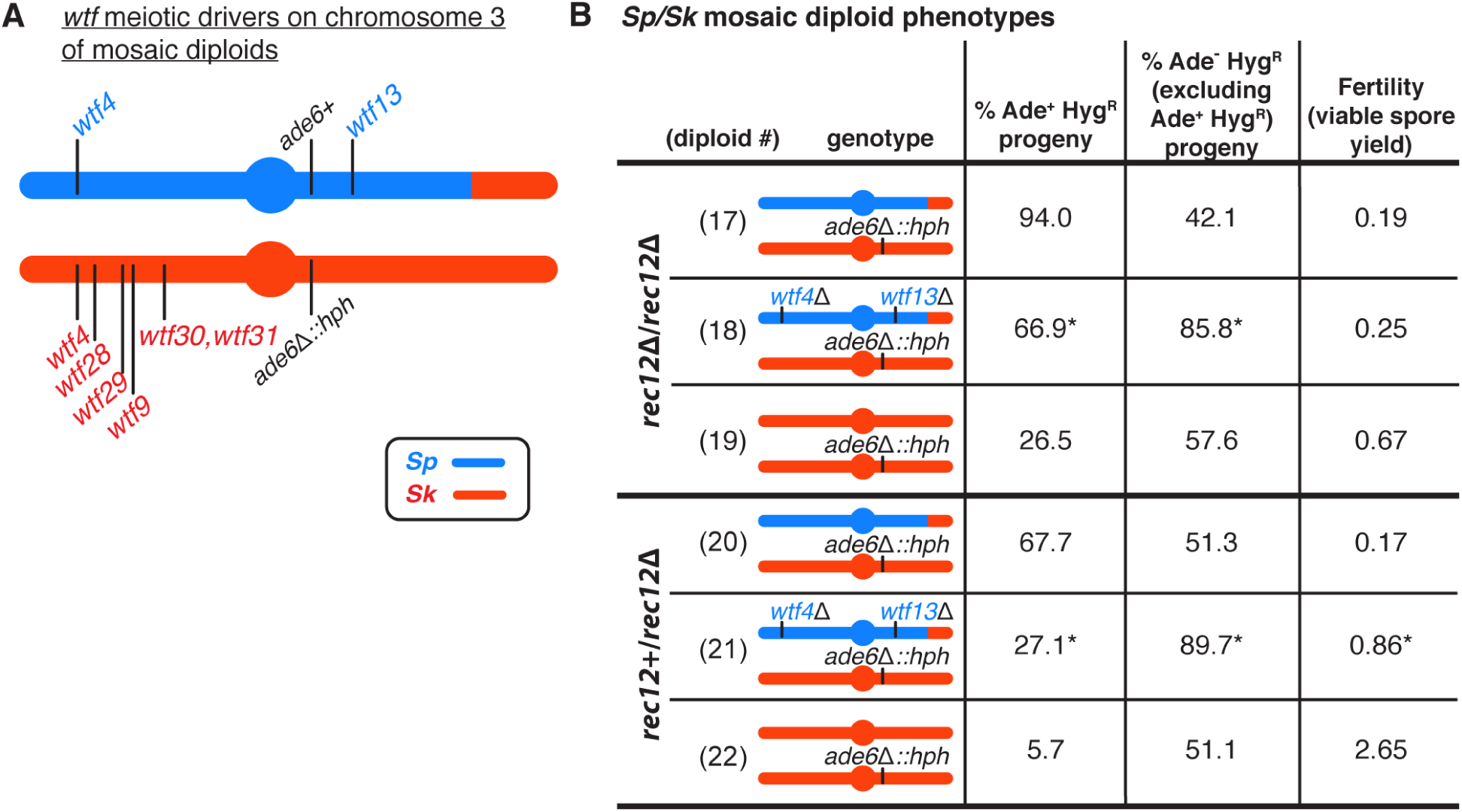
*wtf* meiotic driver competition contributes to the high disomy in *Sp/Sk* mosaic diploids. (A) Schematic of the predicted *wtf* meiotic drivers found on chromosome 3 of the *Sp/Sk* mosaic diploid*. Sp-*derived DNA is depicted in blue and *Sk-*derived DNA in red. (B) Phenotypes of mosaic and control diploids in *rec12+*/*rec12Δ* and *rec12Δ*/*rec12Δ* backgrounds. Allele transmission of chromosome 3 was assayed using markers at *ade6* (linked to centromere 3). To determine the contribution of *wtf* meiotic drivers to the frequency of disomic progeny and fertility, diploid 18 was compared to diploid 17, and diploid 21 was compared to diploid 20. To determine if there was biased allele transmission, diploids 17 and 18 were compared to control diploid 19, and diploids 20 and 21 were compared to control diploid 22. More than 300 viable spores were scored for each diploid. * indicates p-value <0.05 (G-test [allele transmission and Ade+ Hyg^R^ progeny] and Wilcoxon test [fertility]). Raw data can be found in Figure 3—figure supplement 3 and Figure 3—figure supplement 4.

Consistent with our previous observations in a similar mosaic diploid, we saw that the viable spores generated by this diploid were almost exclusively (94%) heterozygous disomes for chromosome 3 (Figure 3B, diploid 17) (Nuckolls et al., 2017). Recombination promotes faithful segregation of chromosomes, so the lack of recombination in this mosaic diploid likely contributed to the high disomy we observed amongst the viable spores. Lack of recombination is, however, insufficient to explain the majority of the phenotype, as *rec12*Δ *Sk* diploids generate only 27% disomic gametes (Figure 3B, diploid 19).

To test if the extremely high frequency of disomic progeny in the *Sp*/*Sk* mosaic diploid was dependent on the competition between *wtf* meiotic drivers, we deleted the predicted *Sp* drivers, *Sp wtf13* and *Sp wtf4.* This eliminated *wtf* driver competition as the remaining *wtf* drivers were either on the same haplotype or homozygous. Consistent with our hypothesis, deleting both *Sp* drivers significantly decreased the frequency of chromosome 3 heterozygous disomes (from 94% to 67%; Figure 3B, diploids 17 and 18), although it surprisingly did not significantly increase fertility. Deleting only one of the two *Sp* drivers was also sufficient to significantly decrease the frequency of disomic progeny (Figure 3—figure supplement 1, diploids 46 and 47). However, deleting only one of the six predicted *Sk* drivers (*wtf4*) had no effect (Figure 3—figure supplement 1, diploid 48).

We also performed analogous experiments in the presence of meiotic recombination by mating the mosaic haploid strain to a *rec12*+ *Sk* strain. Meiotic recombination will produce chromosomes with new combinations of *Sp* and *Sk wtf* drivers. Our model predicts that heterozygous disomic spores will still have a fitness advantage as they are more likely to inherit every *wtf* driver. We observed that this *rec12+*/*rec12*Δ diploid had low fertility, similar to that of the *rec12*Δ *Sp*/*Sk* mosaic diploid (Figure 3B, compare diploid 17 to diploid 20). To assay disomy amongst the progeny of the *rec12+*/*rec12*Δ mosaic, we genotyped the *ade6* locus (Figure 3A). We found that 68% of the viable spores generated by this mosaic diploid were heterozygous at *ade6* (Ade^+^ Hyg^R^). Deleting both *Sp wtf4* and *Sp wtf13* in the *rec12+*/*rec12*Δ mosaic diploid significantly increased fertility and decreased disomy at *ade6* amongst the viable spores from 68% to 27% (Figure 3B, diploids 20 and 21).

We also examined single deletions of *Sp wtf4* or *Sp wtf13* in a Rec12+ mosaic diploid. Deleting *Sp wtf4* or *Sp wtf13* individually decreased disomy amongst the viable spores, but only the *Sp wtf4* deletion significantly increased fertility (Figure 3—figure supplement 1, diploids 49 and 50). These results demonstrate that *wtf* driver competition contributes to the extremely high frequency of disomes amongst the surviving spores and can contribute to low spore viability in these mosaic strain backgrounds. However, *wtf* competition alone was insufficient to explain the total increase in disomy relative to the *Sk* homozygotes (Figure 3, compare diploid 18 to diploid 19; and diploid 21 to diploid 22). Overall, our results support the model that high disomy observed in the progeny generated by outcrossed *S. pombe* diploids is partially due to competing *wtf* meiotic drivers.

### Driver landscapes affect observed recombination rates on colinear haplotypes

Meiotic drivers are often associated with regions of suppressed recombination, such as chromosomal inversions (Dobzhansky and Sturtevant 1938; Dyer et al., 2007; Hammer et al., 1989; Larracuente and Presgraves 2012; Pieper and Dyer 2016; Stalker 1961; Svedberg et al., 2018). This state of recombination suppression is thought to be indirectly caused by the driver as linked loci also enjoy a transmission advantage. However, there is little empirical evidence about how the presence or absence of drivers can directly affect recombination landscapes. Fortuitously, our experiments also allowed us to assay the effects of meiotic drivers on recombination rates. We first assayed recombination between the *ade6* and *ura4* loci of the double driver heterozygote and control diploid described in Figure 2B. The *ade6* and *ura4* loci are over 100 cM apart in the *Sk* control (Figure 2—figure supplement 6). In the *Sk* double driver heterozygotes, this distance fell to 56 cM. We hypothesize this is because recombination can uncouple two of the strongest drivers, *Sp wtf13* and FY29033 *wtf35*, which are found on the same haplotype.

We also analyzed the effect of drivers on recombination in the in *rec12+/rec12*Δ mosaic diploids described in Figure 3. In the mosaic diploids with all drivers intact, we observed *ade6* and *ura4* were 43 cM apart, similar to the 62 cM previously observed in *Sp*/*Sk* hybrids (Figure 3—figure supplement 2) (Zanders et al., 2014). However, when *Sp wtf4* and *Sp wtf13* were deleted from the mosaic strain, the observed genetic distance decreased dramatically to 11 cM (Figure 3—figure supplement 2). We hypothesize this drop is due to preferential death of recombinants, as recombination would lead to haploids failing to inherit all the *Sk* drivers. Overall, our results demonstrate that the meiotic drivers can directly affect recombination landscapes.

### Meiotic driver competition at a single locus selects for atypical meiotic products

The experiments above test scenarios with at least two sets of competing *wtf* drive genes. However, when more closely related isolates mate, the number of heterozygous *wtf* driver loci will be reduced. We modeled this scenario by analyzing the impact of competing one set of meiotic drivers in a diploid. To do this, we analyzed an *Sp* diploid heterozygous for *Sk wtf4* and *Sk wtf28* transgenes integrated at the *ade6* locus (Figure 4A). We compared this diploid to a control heterozygote (empty vectors at *ade6*). Consistent with our hypothesis, *Sk wtf4/Sk wtf28* heterozygous diploids had decreased fertility (13% of the control diploid, Figure 4C) and 77% of the viable progeny inherited both *Sk wtf4-* and *Sk wtf28-*linked drug resistance markers (G418^R^ Hyg^R^ spores, Figure 4C). These phenotypes are not specific to those drivers, the *Sp* strain background, or the *ade6* locus. We also observed an increase in disomic spores in the *Sk* background with different *wtf* drivers, and when we competed *wtf* drivers at the *ura4* locus (Figure 2—figure supplement 1 and Figure 4—figure supplement 1).

**Figure 4.**
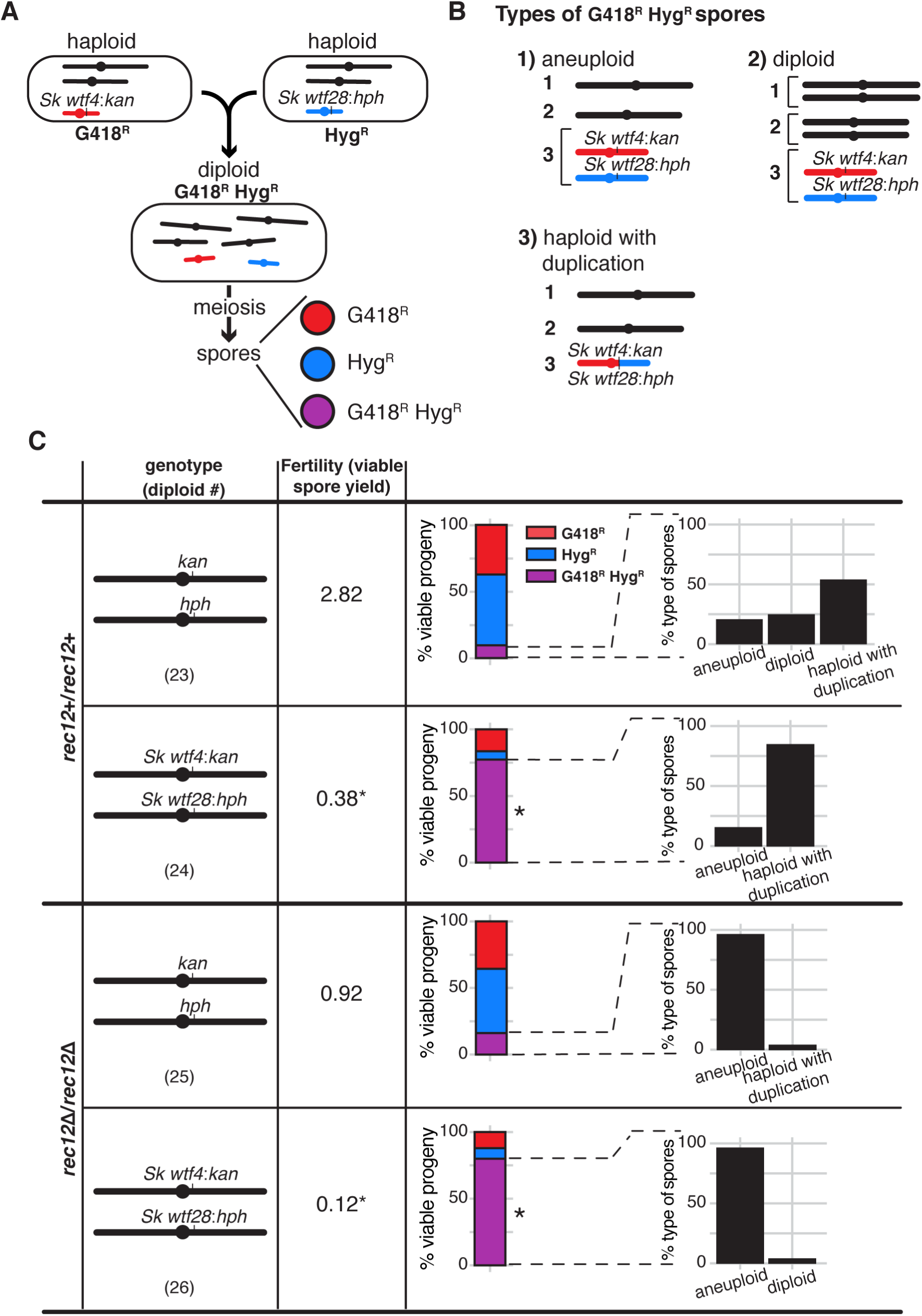
*wtf* competition at a single locus selects for aneuploids, diploids, and spores with a *wtf* duplication. (A) Schematic of diploid heterozygous for the Hyg^R^ and G418^R^ markers at the *ade6* locus. In the experimental diploids (24 and 26), those markers are linked to competing *wtf* drivers (*Sk wtf4/Sk wtf28*). In the control diploids (23 and 25), the drug markers are linked to empty vectors. (B) Types of G418^R^ Hyg^R^ spores. We distinguished these classes using a series of phenotypic and molecular tests (see methods, Figure 4—figure supplement 2, Figure 4—figure supplement 3, and Figure 4—figure supplement 4) (C) Viable progeny observed in control crosses (vector/vector) or with competing *wtf* meiotic drivers at the *ade6* locus in a *rec12*+ (top) or *rec12*Δ (bottom) background. Percentages of G418^R^ Hyg^R^ (aneuploid, diploid, haploid with a duplication event) progeny are shown. For statistical analyses, we compared diploid 24 to control diploid 23, and diploid 26 to control diploid 25. * indicates p-value <0.05 (G-test [G418^R^ Hyg^R^] and Wilcoxon test [fertility]). Raw data can be found in Figure 4— figure supplement 4, Figure 4—figure supplement 5, and Figure 4—figure supplement 6. The data for diploid 23 [excluding inset analyses of G418^R^ Hyg^R^ spores] were previously published in Bravo Núñez et al., 2020.

Although the progeny of diploids with one set of competing meiotic drivers (*Sk wtf4*/*Sk wtf28*) often inherited both *wtf* driver-linked drug resistance alleles, they generally did not exhibit a colony morphology typical of aneuploids. We therefore investigated the ploidy of the progeny of this cross more thoroughly using colony morphology, sporulation phenotypes, phloxin B staining, and a genetic marker loss assay (see methods, Figure 4—figure supplement 2). We were surprised to discover that amongst the G418^R^ Hyg^R^ progeny, only 15.4% appeared to be aneuploid and none appeared to be diploid. Instead, the majority (84.6%) of the G418^R^ Hyg^R^ progeny appeared to be haploid (Figure 4C, diploid 24). We reasoned that an unequal interhomolog crossover event at the *ade6* locus could have led to duplication of the *wtf* driver found on the opposite haplotype (Figure 4—figure supplement 3). Consistent with this idea, we found that the frequency of G418^R^ Hyg^R^ progeny that appeared to be haploid fell in the absence of recombination (*rec12*Δ*/rec12*Δ; Figure 4C, diploid 26). Additionally, we directly tested the unequal crossover hypothesis using PCR. We amplified a potential duplication junction in 22 haploid G418^R^ Hyg^R^ spore colonies and found that unequal crossovers moved both of the competing *wtf* transgenes to the same haplotype in 19 of the 22 tested colonies (Figure 4— figure supplement 3 and Figure 4—figure supplement 4). We detected similar unequal crossover products amongst the G418^R^ Hyg^R^ haploid progeny of the control diploid as well, but at much lower frequencies (Figure 4C, diploid 23 and Figure 4—figure supplement 4). Therefore, we concluded that this type of atypical meiotic product (duplications) was enriched amongst the progeny of the *Sk wtf4/Sk wtf28* heterozygote due to the death of spores that did not inherit both *wtf* genes.

### Fitness costs of meiotic mutants are mitigated or eliminated in diploids with competing *wtf* drivers

Our results demonstrate that when *wtf* drivers compete, such as when *S. pombe* outcrosses, the atypical spores that inherit more drivers are more fit. Disomic spores that inherit two copies of chromosome 3 most likely inherit the maximal number of *wtf* drivers. Therefore, we hypothesized that the fitness costs of decreasing the fidelity of meiotic chromosome segregation might be offset by the fitness benefits of generating more disomic spores when *wtf* drivers compete (Figure 5A and 5B). Consistent with this idea, we previously observed that deleting *rec12* imposed no fitness cost on *Sp*/*Sk* heterozygotes compared to the *Sp/Sp* or *Sk/Sk* homozygotes (Zanders et al., 2014) (Figure 5C). We wanted to know if this was specific to *Sp/Sk* heterozygous diploids or if it might apply more generally to outcrossed *S. pombe* strains. To address this, we compared fertility in the presence and absence of Rec12 in CBS5680/*Sp,* JB844*/Sp*, CBS5680/*Sk,* and JB844/*Sk* heterozygotes. Interestingly, we observed that Rec12 does not significantly promote fertility in heterozygotes as it does in homozygotes (Figure 5C). These results demonstrate that the meiosis fitness optima in inbred strains differs from what is optimal when strains outcross.

**Figure 5.**
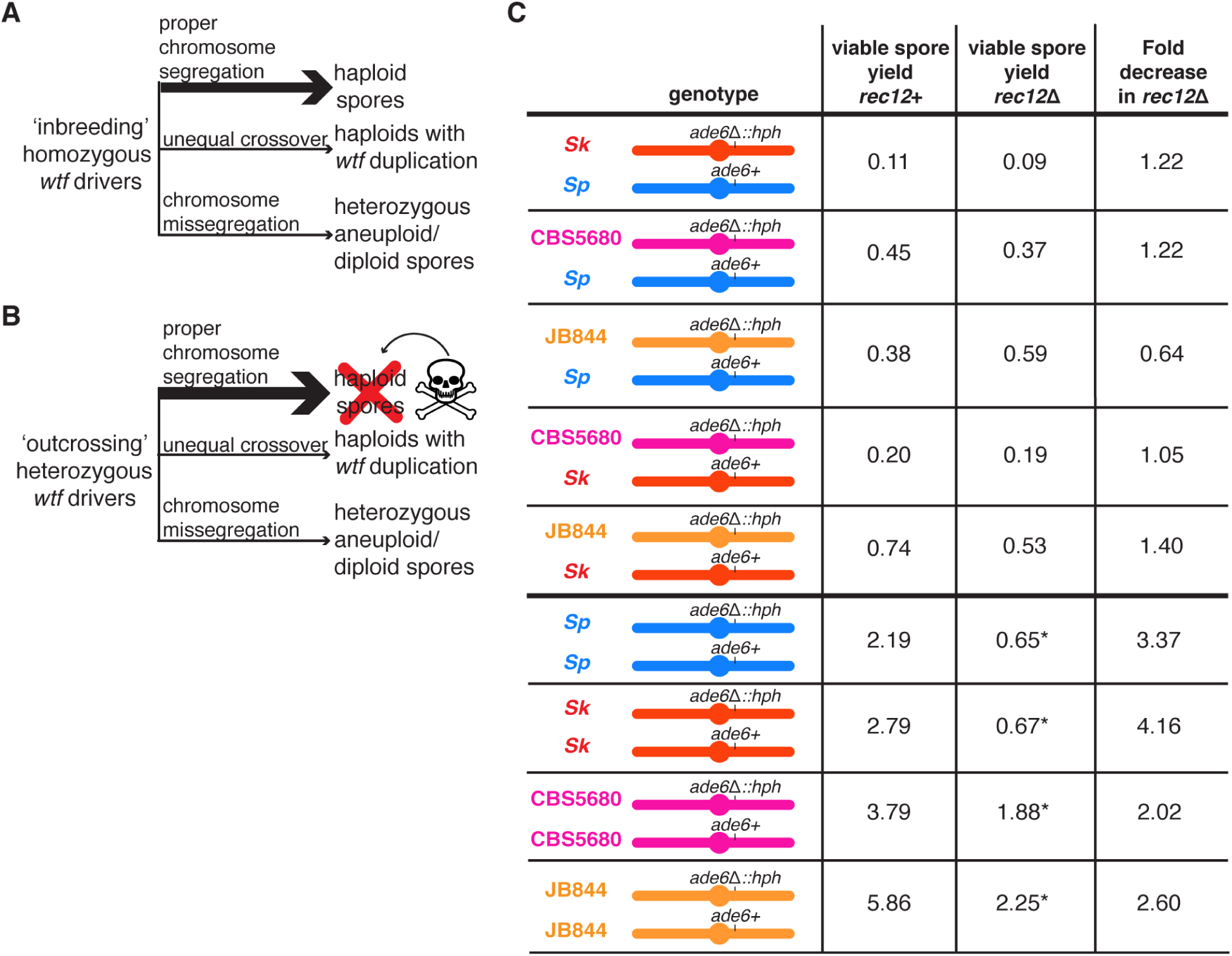
The Rec12 protein does not promote fertility in heterozygous *S. pombe* diploids. (A) Schematic of viable progeny resulting from an ‘inbreeding’ scenario. The three indicated types of spores are viable when *wtf* drivers are homozygous. (B) Schematic of viable progeny resulting from an ‘outcrossing’ scenario where one set of *wtf* drivers is heterozygous. Haploid spores that only inherit one *wtf* allele will be killed by the poison of the *wtf* they did not inherit. Spores that inherit both *wtf* drivers due to a *wtf* duplication or disomy (aneuploidy or diploidy) will survive. Other outcomes of meiosis are not represented in this figure. (C) Viable spore yield values of heterozygous and homozygous *S. pombe* diploids in *rec12+* and *rec12*Δ backgrounds. * indicates p-value < 0.05 (Wilcoxon test) when comparing the *rec12+* to *rec12*Δ fertility values. We compared the viable spore yield of each diploid in *rec12+* and *rec12*Δ backgrounds. At least three, but usually more independent diploids were used to calculate viable spore yield. The data for *rec12+* diploids is repeated from Figure 1. The data for *Sk/Sk rec12*Δ diploid (diploid 19) is repeated from Figure 3. The raw data is reported in Figure 5— figure supplement 1.

We reasoned that competing *wtf* meiotic drivers were contributing to the dispensability of *rec12* in the outcrossed diploids. To test that idea, we assayed the fitness costs of deleting *rec12* in strains with heterozygous *wtf* drivers at one or two loci. We found that in a diploid with one set of heterozygous drivers (*Sk wtf4*/*Sk wtf28* at *ade6*), the cost of deleting *rec12* (*rec12*Δ/*rec12*Δ) was similar to that observed in the wild-type background (3-fold decrease in fertility). However, we found that deleting *rec12* in a genetic background with *wtf* drivers competing at both *ade6* and *ura4* had no cost (Figure 2C, compare diploid 15 to diploid 13). These results support our model that the costs of disrupting chromosome segregation can be offset by the fitness benefits of disomic gametes in the presence of *wtf* driver competition.

We next tested the fitness costs of deleting other genes that promote accurate meiotic chromosomes segregation (*rec10*, *sgo1, moa1,* and *rec8*) in the presence and absence of competing *wtf* drivers (at *ade6*). Rec10 is a component of the meiotic chromosome axis (linear elements) that is required for the formation of most meiotic DSBs (Lorenz et al., 2004; Prieler et al., 2005). The Sgo1 and Moa1 proteins both act at centromeres to promote the disjunction of homologs, rather than sister chromatids, in the first meiotic division (Kitajima et al., 2004; Yokobayashi and Watanabe 2005). Sgo1 protects centromeric cohesion from cleavage and Moa1 promotes monopolar kinetochore attachment of sister chromosomes (Kitajima et al., 2004; Yokobayashi and Watanabe 2005). Finally, Rec8 is the meiotic kleisin that plays key roles in recombination and ensuring proper chromosome segregation in both meiotic divisions (Krawchuk et al., 1999; Watanabe and Nurse 1999; Yoon et al., 2016).

Deleting *sgo1* and *rec10* had a lower fitness cost in the background with heterozygous *wtf* drivers than in a background without *wtf* competition (∼3-fold decrease compared to a 6-7-fold decrease) (Figure 6—figure supplement 1 and Figure 6—figure supplement 2). Remarkably, deleting *moa1* or *rec8* had no effect on fitness in diploids with heterozygous *wtf* drivers, despite the fact that these mutations decrease fertility by 4- and 6-fold, respectively, in the absence of *wtf* driver competition (Figure 6—figure supplement 1 and Figure 6—figure supplement 2).

We reasoned that heterozygous *moa1* (*moa1*Δ/*moa1+*) or *rec8* (*rec8*Δ/*rec8+*) mutants might slightly increase meiotic chromosome missegregation and thus provide a selective advantage in the presence of *wtf* competition (*Sk wtf4*/*Sk wtf28* at *ade6*) by producing disomic spores. Deleting one copy of *moa1* did not significantly alter the frequency of disomic spores in a wild-type background (Figure 6B, diploid 28). In addition, heterozygosity for *moa1* did not suppress the fitness costs of *wtf* competition (Figure 6B, diploid 30). Deleting one copy of *rec8*, however, significantly increased the production of disomic spores in a background without heterozygous *wtf* drivers (Figure 6B, diploid 27). This suggests that *rec8* exhibits haploinsufficiency and reducing Rec8 protein levels may lead to chromosome segregation errors during meiosis. Consistent with this observation, mutations that reduce the levels of the meiotic cohesin can lead to meiotic defects in mice and flies (Murdoch et al., 2013; Subramanian and Bickel 2008). As hypothesized, *rec8* heterozygosity also increased the fertility of diploids with competing drivers (Figure 6B, diploid 29). Overall, these results suggest that the costs of disrupting chromosome segregation can be partially or totally alleviated by the increased protection against *wtf* drivers gained by generating more heterozygous disomic spores.

**Figure 6.**
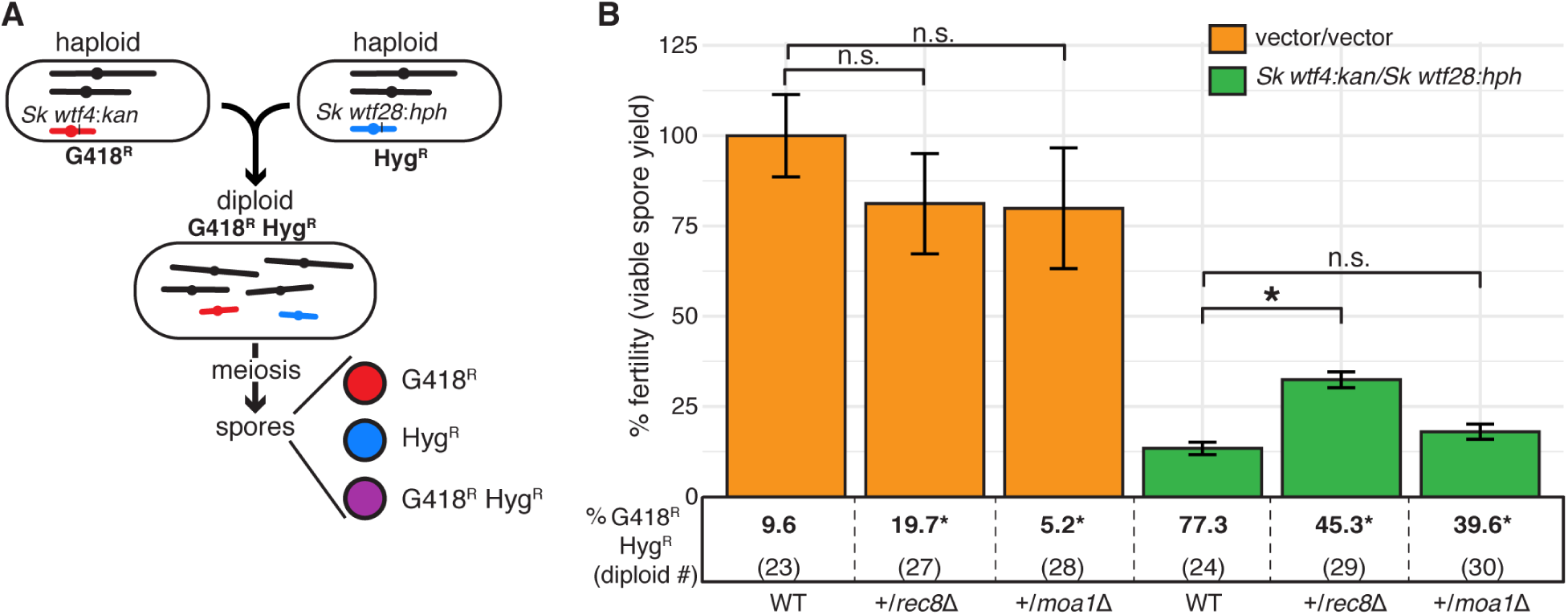
A heterozygous *rec8* mutation increases fitness when *wtf* meiotic drivers are in competition. (A) Schematic of diploid heterozygous for the Hyg^R^ and G418^R^ markers at the *ade6* locus. (B) Fertility was measured using the viable spore yield assay in diploids with markers linked to competing *wtf* drivers (*Sk wtf4/Sk wtf28*) or empty vectors (vector/vector). Error bars represent the standard error of the mean. Underneath each bar graph is the % of G418^R^ Hyg^R^ (aneuploid, diploid, or haploid with duplication event) progeny for each diploid. * indicates p-value <0.05 (G-test [G418^R^ Hyg^R^] and Wilcoxon test [fertility]). For statistical analyses, diploids 27 and 28 were compared to diploid 23, and diploids 29 and 30 were compared to diploid 24. Data for diploids 23 and 24 are repeated from Figure 4. Raw data are found in Figure 6—figure supplement 3 and Figure 6—figure supplement 4.

### Driver competition can facilitate the maintenance or spread of alleles that disrupt meiotic chromosome segregation fidelity in a population

Our experiments demonstrate that the effects of meiotic mutants can be quite different in heterozygous *S. pombe* wherein *wtf* drivers are competing. To explore this idea further, we turned to population genetic modeling to analyze how drivers affect the evolution of variants that decrease the fidelity of meiotic chromosome segregation.

Our model analyzes the evolutionary fate of a hypothetical mutation that disrupts the segregation of chromosome 3, which houses the majority of *wtf* drivers. For the sake of simplicity, our model assumes that chromosome 3 exhibits whole-chromosome drive. The model also considers six parameters (Figure 7A). The first two parameters relate to the *wtf* drivers. We varied the number of driving alleles in the population (*n*) and the strength of their drive (*t*). Each driving allele was assumed to be at an equal frequency in the population and have the same strength of drive. The next parameters relate to the meiotic mutation. We varied the level of chromosome missegregation caused by the mutation (*f*) from 0 (no mutant phenotype) to 1 (50% of the resultant spores are heterozygous disomes and the remaining 50% of the spores lack chromosome 3 and are thus inviable). We considered the dominance of the mutation (*h*) and any additional fitness costs (*s_m_*) the mutation may incur, such as potential costs relating to the missegregation of other chromosomes. Finally, we considered additional fitness costs (*s_s_*) disomic spores might bear. The full description of the model and additional analyses are presented in Supplementary File 1. We found that a mutation with no fitness costs (*s_m_* and *s_s_*=0) that disrupts meiotic segregation could invade a population when:

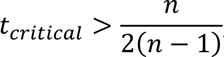

**Figure 7.**
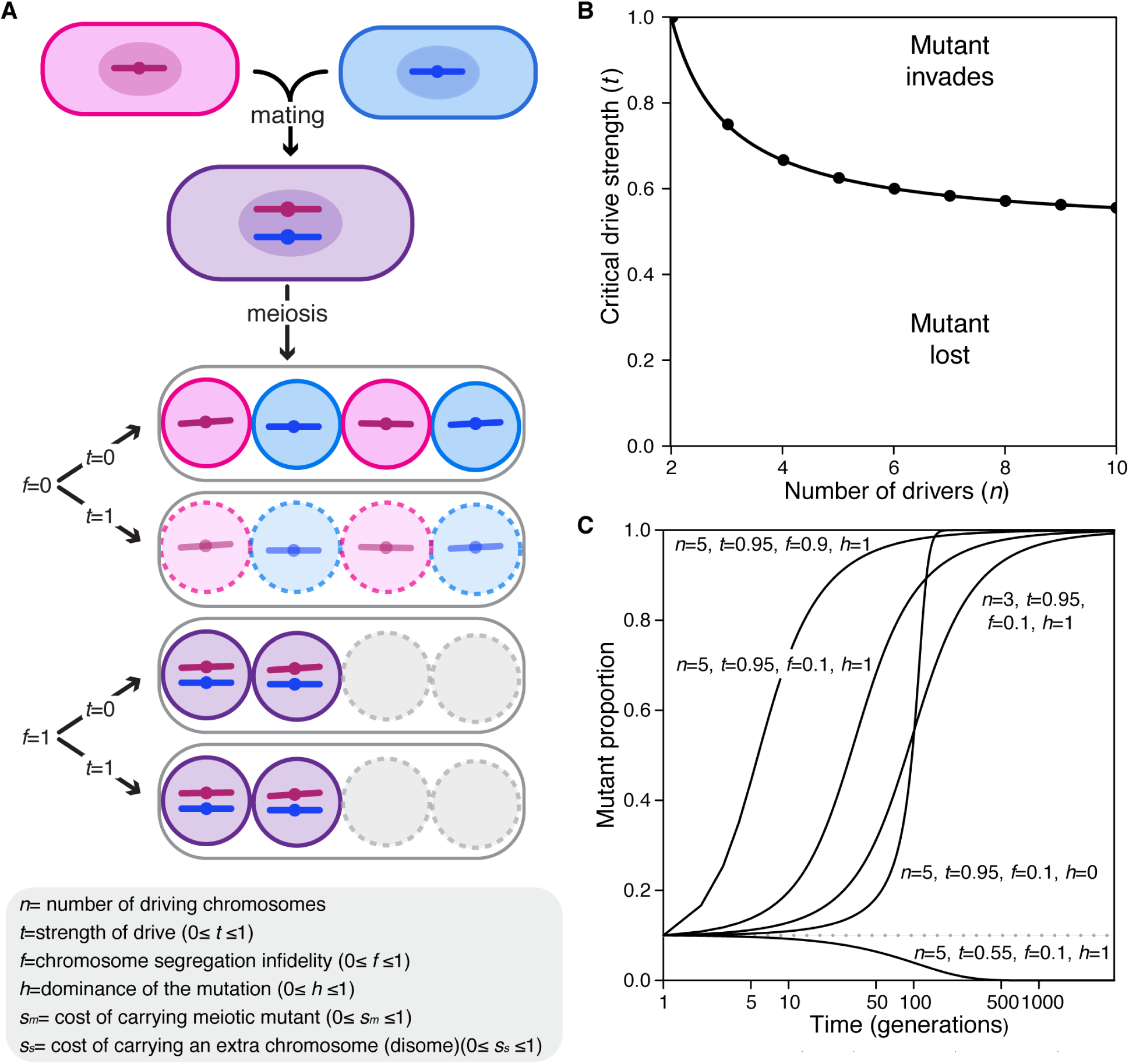
Population genetics of a meiotic mutant in response to meiotic drive. (A) Schematic of the spore progeny generated *S. pombe* diploids when the infidelity of chromosome segregation (*f*) is 0 or 1 and the strength of drive (*t*) is 0 or 1. (B) Critical drive strength (*t_critical_*) for invasion of the segregation infidelity mutant given *n* drivers. This assumes that there is no cost of the meiotic mutant. (C) Trajectories of meiotic mutants starting at a frequency of 0.1. Note that the X-axis is a log scale.

where *t_critical_* is the value of drive strength necessary for such invasion in a population of *n* drivers. Interestingly, the higher the strength of drive (*t*), the lower the number of drivers required for mutant invasion (Figure 7B). Importantly, our empirical work demostrates that drive strength is generally high (*t* >0.9) and that there are ample *wtf* drivers (n>5) leaving parameter space for mutants to invade even if they incur considerable costs (*s_m_* and *s_s_*) (Figure 7—figure supplement 1) (Bravo Núñez et al., 2018; Bravo Núñez et al., 2020; Eickbush et al., 2019; Hu et al., 2017; Nuckolls et al., 2017). With fitness costs applied to the equation above, increasing the number of drivers in the population and decreasing the value of these associated costs both increase the likelihood that the mutation can invade the population (Figure 7—figure supplement 1 and Supplementary File 1).

To get a broader perspective on the potential evolutionary trajectories of the segregation fidelity mutant, we varied the described parameters and plotted the results. These analyses all started with the mutant at a frequency of 0.1 in the population. We found that changes in some parameters dramatically influenced the trajectories. For example, with all other parameters fixed, at *t =* 0.55 the mutant is lost, while at *t* = 0.95, the mutant is fixed in the population (Figure 7C). With the parameters plotted, the dominance of the mutation, *h*, has little influence on the fate of the mutation but does influence the rate at which that fate is reached. Finally, in our model where the costs are applied, the cost of the mutation, the cost bore by disomic spores, and the degree of segregation infidelity also influence whether or not invasion occurs (Figure 7—figure supplement 1).

Overall, the results of the model are consistent with our experimental results. Both types of analyses support the idea that meiotic drivers can change the selective landscape of meiosis in outcrossed *S. pombe*. Instead of meiotic mutants being removed by negative selection due to fitness costs, variants that decrease the fidelity of chromosome segregation can be advantageous due to pervasive meiotic drivers.

### Natural isolates greatly vary in their propensity to produce disomic gametes

Given that mutations that increase the frequency of disomic gametes can be beneficial in scenarios that model outcrossing, we questioned if disomy-promoting variants are present in natural populations. To address this question, we assayed the frequency at which 17 different homozygous natural *S. pombe* isolates generated disomic spores using centromere 3-linked markers (at *ade6*) (Figure 8A). We found several natural isolates produce a similar fraction of disomes as the lab strain (<10%), but others produced as many as 32% heterozygous disomic spores (Figure 8B). This result is consistent with the idea that it is not strongly deleterious for *S. pombe* strains to generate non-haploid spores at a high frequency.

**Figure 8.**
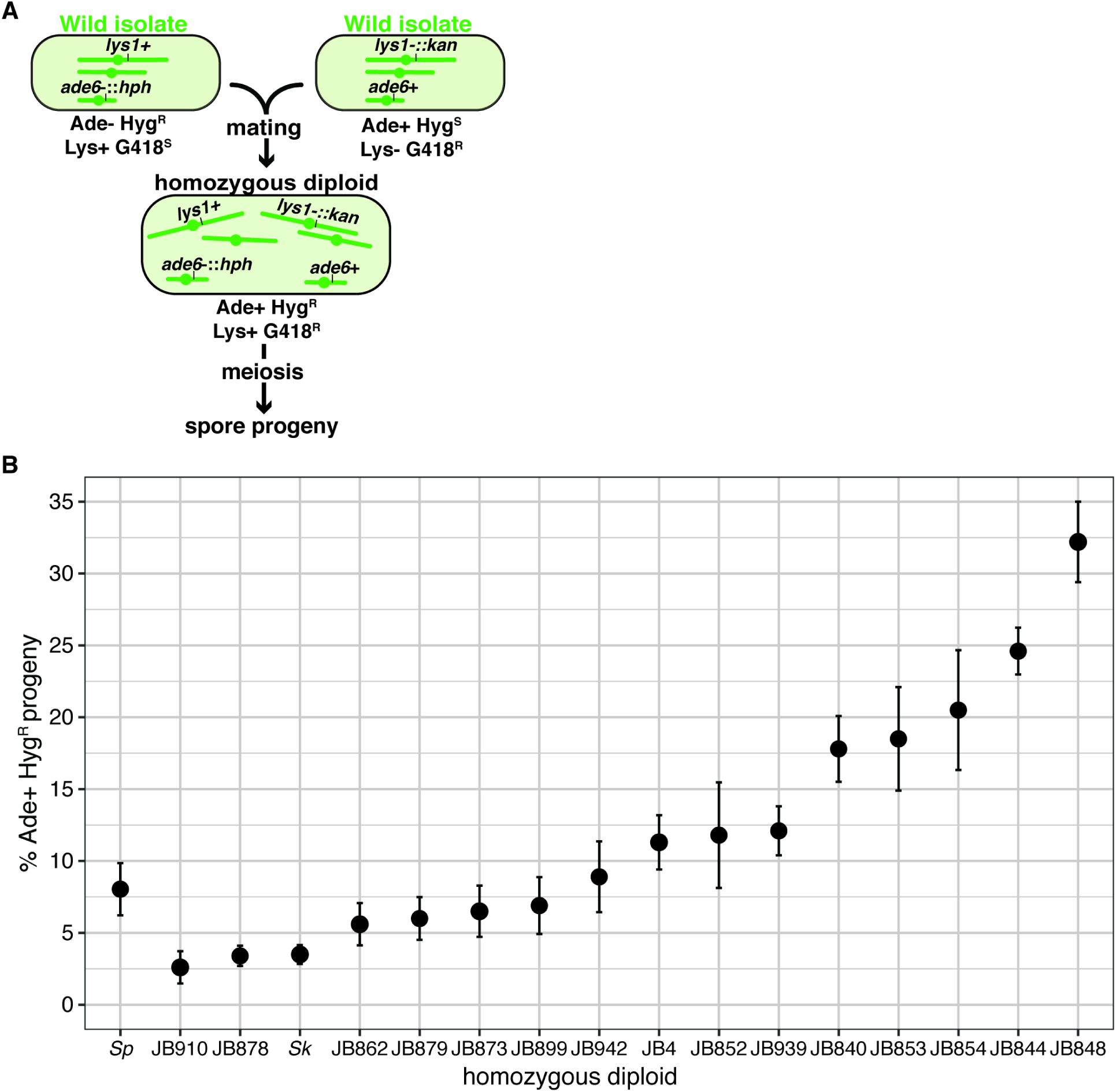
Homozygous *S. pombe* diploids generate variable frequencies of disomic progeny. (A) Schematic of the experimental approach used in Figure 8B. The *ade6* marker is linked to centromere 3. (B) Phenotypes of homozygous *S. pombe* diploid strains. Allele transmission of chromosome 3 was assayed using co-dominant markers at *ade6* (*ade6+* and *ade6-*::*hphMX6*). The *ade6+* allele confers an Ade+ phenotype, while the *ade6*-::*hphMX6* cassette provides resistance to Hygromycin B (Hyg^R^). The errors bars represent the standard error of the mean for each strain. Heterozygous aneuploid or diploid spores inherit both markers and are thus Ade+ Hyg^R^. More than 200 viable spores were scored for each diploid and more than three independent diploids were assayed. Raw data are shown in Figure 8—figure supplement 1.

## Discussion

### Meiotic drivers shape the evolution of *S. pombe* meiosis

Most genetic analyses of meiosis are performed in inbred (homozygous) organisms. This approach has been incredibly powerful. It enabled Mendel to establish the founding principles of genetics with his true-breeding peas and facilitating countless discoveries over the last 150 years (Abbott and Fairbanks 2016). However, studying inbred model systems with very low genetic variability has limitations in that phenotypes that can only be observed in heterozygotes remain hidden. These phenotypes can include those caused by selfish genetic elements, like meiotic drivers, that act primarily in heterozygous or outcrossed organisms. This work highlights the stark differences in fitness optima that can exist between gametogenesis in inbred homozygotes and in heterozygotes generated by outcrossing.

When *S. pombe* inbreeds to generate homozygotes, the effects of *wtf* meiotic drivers are largely invisible as every resulting spore inherits the necessary Wtf antidotes required to neutralize the Wtf poisons generated during gametogenesis. Granted these homozygotes faithfully segregate their chromosomes during meiosis and generate four haploid spores, the fitness of these diploids will be relatively high. However, mutations such as *rec12Δ*, *rec8Δ,* or *moa1Δ* will decrease the relative fitness of such diploids, as these mutations decrease the fidelity of chromosome segregation and spore viability (Davis and Smith 2003; Martín-Castellanos et al., 2005; Yokobayashi and Watanabe 2005).

When *S. pombe* outcrosses, the observed outcomes of meiosis are different from those observed under inbreeding, at least in part due to *wtf* meiotic drivers. These differences include changes in fertility, spore ploidy, recombination frequencies, and the fitness costs (or benefits) of inaccurate meiotic chromosome segregation. It is important to stress that the effects *wtf* drivers exert on meiosis are entirely indirect. There is no evidence that Wtf proteins directly participate in or affect the molecular mechanisms of recombination or chromosome segregation. Rather, the effects of the Wtf proteins are observed after the completion of meiosis. Despite this, the Wtf proteins have the power to indirectly affect the molecular steps of meiosis as changes in the fidelity of chromosome segregation or other aspects of gametogenesis can alter the number of spores destroyed by drive.

The *S. pombe* isolates in which *wtf* genes have been assembled carry between 4-14 intact *wtf* drive genes (Eickbush et al., 2019; Hu et al., 2017). The *wtf* genes are amongst the most rapidly evolving genes in *S. pombe*, which means outcrossing often generates extensive *wtf* driver heterozygosity. This leads to extensive death of haploid spores in outcrossed strains because 1) Wtf^antidote^ proteins appear to neutralize only the Wtf^poison^ proteins with highly similar or identical C-termini and 2) it is unlikely that haploid spores will inherit all drivers and thus encode all Wtf^antidote^ proteins (Bravo Núñez et al., 2018; Bravo Núñez et al., 2020). Hence, generating the maximal number of haploid spores does not maximize the fitness of outcrossed *S. pombe*.

Instead, fitness is maximized in outcrossed *S. pombe* when the spores inherit as many drivers as possible. If there is just one locus with heterozygous drivers, an unequal crossover event can place both drivers on the same haplotype and allow a haploid spore to survive. It is possible this type of selection may be occurring in nature, as all assayed strains contain multiple loci with 2-3 *wtf* genes in tandem (Eickbush et al., 2019).

Since *wtf* drivers are numerous in outcrossed *S. pombe*, it is very unlikely for a haploid spore to inherit every driver. In this scenario, disomic spores that inherit the two different copies of chromosome 3, which carries nearly every *wtf* gene, are most likely to inherit every driver and survive. Importantly, an extra copy of chromosome 3 is the only aneuploidy tolerated in *S. pombe* (Niwa et al., 2006). We have previously speculated that the *wtf* gene family specifically expanded on chromosome 3 as aneuploid spores provide an avenue to mitigate the fitness costs of multiple drivers (López Hernández and Zanders 2018; Zanders et al., 2014). The results of this study support and expand on that model. Specifically, we now show that drivers can create a selective landscape wherein variants that decrease the fidelity of chromosome segregation to generate more disomic gametes can be favorable. In these cases, the fitness costs of mutating genes like *rec12*, *moa1,* or *rec8* can be offset by the fitness benefits of increased disomy. Our work also adds to previous work demonstrating meiotic drive-independent adaptive potential of aneuploid spores in other fungi (Chuang et al., 2015; Ni et al., 2013).

It is not clear how often *S. pombe* outcrosses in the wild, and population genetics estimates are confounded by drive and repressed recombination in hybrids (Farlow et al., 2015; Fawcett et al., 2014; Jeffares et al., 2015; Tusso et al., 2019). Many *S. pombe* isolates can switch mating type during clonal growth and thus mate with nearby clonal cells when starved for nutrients (Egel 1977). This undoubtably leads to frequent inbreeding in *S. pombe* and could thus promote selection against mutations that increase the frequency of disomic gametes. However, when outcrossing occurs, mutations that increase disomy can have a selective advantage. A mix of inbreeding/outcrossing strategies could lead to the maintenance of variation in the frequency at which meiosis generates disomic spores. Consistent with this, we observed such variation amongst the natural isolates assayed in this study (Figure 8). Strikingly, the strains with the “highest” meiotic fidelity still make ∼5% disomic spores, suggesting that chromosome 3 missegregates during the first meiotic division in one out of ten meioses.

### The effects of drive on the evolution of gametogenesis outside of *S. pombe*

A growing body of evidence indicates that meiotic drive is pervasive in eukaryotes, and more drivers are identified each year. This includes the gamete-killing type of meiotic drivers described in this work (Bauer et al., 2012; Bravo Núñez et al., 2018; Bravo Núñez et al., 2020; Burt and Trivers 2006; Didion et al., 2015; Grognet et al., 2014; Hammond et al., 2012; Hu et al., 2017; Larracuente and Presgraves 2012; Long et al., 2008; Nuckolls et al., 2017; Pieper et al., 2018; Rhoades et al., 2019; Vogan et al., 2019; Xie et al., 2019; Yang et al., 2012; Yu et al., 2018), but also extends to other drivers that use completely different methods to gain a transmission advantage. For example, biased gene conversion favoring unbroken DNA during meiotic recombination is a form of meiotic drive tied to the mechanisms of double-strand break repair (Marais 2003). This type of drive shapes recombination landscapes and likely promotes the rapid evolution of at least one key recombination protein found in many mammals, including humans (Grey et al., 2018; Úbeda et al., 2019). Other meiotic drivers exploit the asymmetry of female meiosis to promote their transmission into the one viable meiotic product (i.e. the oocyte) (Akera et al., 2017; Akera et al., 2019; Dawe et al., 2018; Kato Yamakake 1976; Rhoades 1942). This type of bias has been hypothesized to drive the widespread rapid evolution of karyotypes, centromere sequences, and centromeric proteins (de Villena and Sapienza 2001; Henikoff et al., 2001; Rosin and Mellone 2017). In addition, drive during female meiosis in mice can even generate selective pressure to alter the timing of the first meiotic division (Akera et al., 2017; Akera et al., 2019).

It may be tempting to disregard the *wtf* genes within *S. pombe* as an anomaly. However, meiotic drivers are ubiquitous, and drive represents an incredibly powerful evolutionary force. Appreciating how *wtf* genes affect *S. pombe* will likely provide important insights into how genetic parasites can shape the evolution of meiosis in other eukaryotes.

## Materials and Methods

### Strain construction: *S. pombe* natural isolates

All yeast strain names and genotypes are described in Supplementary file 2. We made the *lys1*Δ::*kanMX4* and *ade6*Δ::*hphMX6* alleles used in Figure 1 as described in Zanders et al., 2014. Using the standard lithium acetate protocol, we independently transformed the cassettes into seven different *S. pombe* natural isolates (JB844, JB1172, CBS5680, JB873, JB939, JB929, and NBRC0365). However, we were only successful at transforming both markers into JB844, JB1172, CBS5680, and NBRC0365. We could not find conditions in which to mate and sporulate NBRC0365.

To generate a *rec12*Δ::*ura4+* deletion in the CBS5680 strain background, we first made a *ura4-D18* mutation in SZY2111 (*ade6*Δ::*hphMX6* in CBS5680). We amplified the *ura4-D18* allele from SZY925 using oligos 35 and 38 and transformed it into SZY2111 to generate SZY3949. We then amplified the *rec12*Δ::*ura4+* cassette from SZY580 using oligos 1194 and 1077 and transformed the cassette into SZY3949 to generate SZY3995. We confirmed the *rec12* deletion via PCR using oligos (1120 and 1108) that bind 730 bases upstream and 224 bases downstream of the deletion cassette. We generated the *rec12*Δ::*ura4+* deletion in the *lys1*Δ::*kanMX4* background of the CBS5680 isolate via crosses. We generated the *rec12*Δ strain in JB844, similarly to how we generated it in the CBS5680 strain.

We found it difficult to make gene deletions in many of the natural isolates used in this study. We had more success, however, making mutations using integrating vectors. Because of this, we used integrating vectors to generate the genetic markers used in Figure 8. We used pSZB386 to generate haploid strains with a *hphMX6* marker at *ade6*, without deleting the *ade6* gene. We cut this plasmid with KpnI and transformed it into different natural isolates, selecting for transformants that were resistant to Hygromycin B and red on media with low adenine (Bravo Núñez et al., 2018). To generate strains with a *kanMX4* marker at *lys1,* we first ordered a gBlock from IDT (Coralville, IA). This gBlock contained ∼1000 bp from the middle of the gene in which we replaced 50 bp from the center with a KpnI site. We then cloned the gBlock into the BamHI and SalI sites of pFA6 to generate pSZB816. We then digested pSZB816 with KpnI and transformed it into different *S. pombe* isolates. We then screened for transformants that grew on plates containing G418 and were not able to grow on media lacking lysine.

### Plasmid construction: integrating vectors with *wtf* alleles

Most of the integrating vectors containing *wtf* alleles were previously described in Bravo Núñez et al., 2018; Bravo Núñez et al., 2020; and Nuckolls et al., 2017. To generate the additional *ade6-* and *ura4-*integrating vectors unique to this work, we cloned the *wtf* genes of interest into the integrating plasmid backbones and confirmed them via sequencing. The DNA templates, oligos, and restriction enzymes used are described in Supplementary file 4.

To generate pSZB923 (*Sk wtf4:natMX4*), we first digested pSZB189 (which contains *Sk wtf4*) with SacI to release the *Sk wtf4* cassette. We then cloned the cassette into the SacI site of pSZB849.

### Deletions of the *moa1, rec10,* and *sgo1* genes in *Sp*

We made *moa1*, *rec10*, and *sgo1* gene deletions, using standard deletion cassettes and transformation. To make the *moa1*Δ::*natMX4* cassette, we amplified the upstream region of *moa1* with oligos 1673+1187 and the downstream region with oligos 1190+1191 (or 1190+1674) using SZY643 as a template. We also amplified the *natMX4* gene (with oligos 1675+1189) using pAG25 as a template (Goldstein and McCusker 1999). We then stitched all the PCR fragments together using overlap PCR and transformed this fragment into SZY44 and SZY643 to make strains SZY2479 and SZY2481, respectively. We confirmed the integration of the deletion cassette at the *moa1* locus using oligos AO638+1192, AO1112+1191, and 1701+1702. We also checked that the *moa1* gene was not present somewhere else in the genome by using two oligos (1703+1704) within *moa1*.

To generate a *rec10*Δ::*natMX4* strain, we first amplified the upstream region and the downstream region of *rec10* from SZY643 using oligos 1723+1724 and oligos 1727+1728, respectively. We also amplified the *natMX4* cassette from pAG25 using oligos 1725+1726 (Goldstein and McCusker 1999). Using overlap PCR, we stitched the three PCR fragments together and then transformed the final deletion cassette into SZY643 and SZY44 to make strains SZY2517 and SZY2519, respectively. To confirm the integration of the cassette at the correct locus, we used oligo pairs 1731+AO638 and 1732+AO1112. We also confirmed the absence of the wild-type *rec10* gene by using internal oligos (1729+1730).

To make the *sgo1*Δ*:: hphMX6* allele, we amplified the sequences upstream and downstream of *sgo1* from SZY643 using oligos 1224+1225 and 1228+1229. We also amplified the *hphMX6* cassette from pAG32 using oligos 1226+1227 (Goldstein and McCusker 1999). We then stitched all the PCR fragments together using overlap PCR and transformed the cassette into yeast to generate strains SZY1735 and SZY1736. We confirmed the *sgo1* deletion using oligos AO638+1230, AO1112+1231, and 1230+1231. We then confirmed the absence of the wild-type gene using an internal oligo pair (2088+2089).

### *Sp wtf4* deletion

To delete *Sp wtf4,* we utilized the CRISPR/Cas9-based method described in Rodriguez-Lopez et al., 2016. We first cloned a plasmid (pSZB570) encoding Cas9 and a guide RNA targeting *Sp wtf4*. To do that we first amplified pMZ379 (plasmid containing Cas9) using oligos 1206+1207. These oligos contained the single guide RNA (sgRNA) sequence that targets the *Sp wtf4* gene. We then ligated the ends together to generate pSZB570.

We also made a deletion cassette to knockout the *Sp wtf3* and *Sp wtf4* locus. We used oligos 574+1138 and 1139+471 to amplify the upstream and downstream sequence of the locus using SZY580 as a template. We then stitched these two fragments together using overlap PCR. We then transformed the deletion fragment and pSZB570 into SZY1595 to generate SZY1699. We used oligos 1069+543 to confirm the *Sp wtf4* deletion. We also Sanger sequenced the PCR fragment and found that we had only knocked out the *Sp wtf4* gene, not the entire *Sp wtf3* and *Sp wtf4* locus.

### *Sp wtf13* deletions

The *Sp wtf13* deletions were made similar to the ones described in Bravo Núñez et al., 2018. Using SZY580 as a template, we amplified the upstream (with oligos 1048 and 1049) and downstream (with oligos 1052+1053) sequence of *Sp wtf13*. Additionally, we amplified the *kanMX4* cassette using oligos 1050+1051 using pFA6 as a template (Wach et al., 1994). We then stitched the upstream region, *kanMX4*, and the downstream region together using overlap PCR to make an *Sp wtf13*Δ::*kanMX4* cassette. After, we transformed this fragment into SZY580 to generate SZY1391-SZY1394.

To generate the *Sp wtf13*Δ::*kanMX4 Sp wtf4*Δ strain, we first digested pAG25 with EcoRV and BamHI to release the *natMX4* cassette (Goldstein and McCusker 1999). We then used this cassette to switch the *kanMX4* marker (at the *his5* gene) from the SZY1699 strain via transformation to generate SZY1981 and SZY1982. We then transformed the *Sp wtf13*Δ::*kanMX4* deletion cassette into SZY1981 and SZY1982 to generate strains SZY2008 and SZY2010, respectively. We confirmed these deletions using a series of PCR reactions. We used two oligo pairs with one oligo outside of the deletion cassette and one oligo internal to the *Sp wtf13* gene (1058+1059 and 1060+1061) and two oligo pairs in which one oligo was external to and one oligo was within the deletion cassette (1058+AO638 and 1061+AO112).

### Fertility and allele transmission

We assayed fertility and allele transmission as described in Bravo Núñez et al., 2020.

Some of the spore colonies from *ade6*Δ::*hphMX6/ade6+; lys1+/lys1*Δ::*kanMX4* CBS5680 diploids (SZY2213/SZY2111) were small and red. When we determined their genotype, the colonies were adenine auxotrophs and took five days to grow when replicated to fresh media. We supplemented the plates with more adenine, but the colonies did not grow faster. This slow growth phenotype was curiously not observed in the ade-parental haploid (SZY2111).

### Recombination frequency within the *ade6* and *ura4* interval

To determine the recombination frequency for diploids 20 and 21, we needed to distinguish the *ura4* allele (*ura4-294* or *ura4*Δ*::kanMX4*) via PCR. We amplified the *ura4* locus using two sets of oligos (34+37 and 34+AO638).

### Determining ploidy of spore colonies

In the various tests to assay the ploidy of the spore colonies for Figure 4, we compared the spore colony phenotypes to the following control strains: a homothallic haploid (SZY925), a heterothallic haploid (SZY1180), a diploid (SZY925/SZY1180), and aneuploid (irregular colonies generated by a cross between SZY1994 and SZY1770) controls. The ploidy of the strains was determined by how closely a test strain resembled one of the controls in the following tests:

#### Spore colony morphology

To determine the morphology of the spore colonies from different diploids, we diluted the spores to get isolated colonies on YEA+S (0.5% yeast extract, 3% dextrose, agar, and 250 mg/L adenine, histidine, uracil, leucine, and lysine). We then imaged the colonies using the Canon EOS Rebel T3i plate imager. We then marked each cell in ImageJ and determined the genotype of each spore colony via replica plating. These images allowed us to correlate the morphology of each spore colony with its genotype. For the spores that had resistance to both G418 and Hygromycin B, we assessed if the colonies were either large, medium, or small, and if the morphology was either round or irregular. Round and large colonies are typical of diploid colonies, while round and medium colonies are typically haploid. Small and irregular-shaped colonies are characteristic of ‘sick’ colonies or aneuploids (Niwa et al., 2006).

#### Chromosome loss assay

To determine the ploidy of the spore colonies, we determined the frequency at which each of the G418^R^ Hyg^R^ strains lost one of the drug markers during vegetative growth. Aneuploids frequently lose their extra chromosome randomly during vegetative growth (Niwa et al., 2006). Diploids and haploids are expected to be more stable. We began by culturing ∼26 G418^R^ Hyg^R^ spore colonies produced by each experimental diploid (diploids 23-26), along with six haploid controls (deemed haploid due to the presence of only one drug marker) in 5 mL of YEL (0.5% yeast extract, 3% dextrose, and 250 mg/L adenine, histidine, uracil, leucine, and lysine) with shaking at 32°C for 24 hours. The next day (day 1), we diluted the cultures and plated the cells on YEA to assay the presence or absence of the drug resistance markers. We also made 1:10 dilutions into 650 μL of YEL in 96-well plates and grew them for 24 hours. We repeated this for five days. Aneuploids readily lost one marker randomly around day 1 or 2, so that a high fraction (17-100%) of the colonies generated by plating the culture were no longer resistant to both drugs. Haploids and diploids, however, maintained both markers for all 5 days because almost all (∼90%) of the colonies generated by plating the culture were still resistant to both drugs. The marker maintenance value was calculated by dividing the number of G418^R^ Hyg^R^ colonies by the total number of colonies that grew on YEA+S.

#### Phloxin B staining

Phloxin B was used to differentiate between diploid strains and haploids. Phloxin B is a dye that enters cells with compromised membranes, staining them red (Forsburg and Rhind 2006). Diploids have a higher concentration of dead cells within a colony than haploids due to their lower stability resulting in a red stain. Haploids are much more stable, which reduces the ability of phloxin B to enter cells, leading haploids to appear white. Haploid strains that are homothallic (h^90^) look pink (Figure 4—figure supplement 2). We spotted 10 μL of the saturated culture of day 1 (described above) onto YEA+S plates containing 5 mg/L phloxin B and grew the cells at 32°C overnight. We then determined if the spots were red, pink, or white by comparing them to the controls.

#### Microscopy and iodine staining from SPA plates

Using the cultures from day 1 (described above), we also spotted 10 μL of the saturated culture of each strain and controls onto SPA two plates. We then placed these plates at 25°C for 20 hours. From the first SPA plate, we imaged the cells on a Zeiss (Germany) Observer Z.1 widefield microscope with a 40X (1.2 NA) water-immersion objective and acquired the images using the μManager software. Twenty hours was enough time for diploids to sporulate but not enough for homothallic haploid strains to mate, form diploids, and sporulate, allowing for further distinction between these ploidies. Although this assay did not reliably allow us to distinguish aneuploid cells that could not sporulate (as they would resemble the heterothallic haploid control), we were able to score some homothallic ‘aneuploid’ strains due to the presence of asci with an abnormal shape or number of spores.

The second SPA plate was stained with iodine (Forsburg and Rhind 2006) after 20 hours at 25°C. Iodine vapors stain the starch present in spore walls a dark brown color, while heterothallic haploid cells that cannot sporulate appear yellow. Diploids stain dark brown, while homothallic haploids that first needed to mate in order to sporulate stained light brown (Figure 4—figure supplement 2).

#### PCR assay for duplications

Some of the G418^R^ Hyg^R^ progeny we tested appeared to be haploids based on the assays described above. We reasoned they could be the result of an unequal crossover putting both marker genes on one chromosome. To test this, we used two sets of oligo pairs (2415+2417 and 2416+2418). These PCR reactions will only work if the drug cassettes are found in tandem. To confirm that the presence of only one cassette would not lead to band amplification, we used the haploid parental strains (SZY925, SZY1180, SZY887, and SZY1293) as negative controls.

## Acknowledgments

We thank the members of the Zanders lab for their helpful comments on the manuscript. We are grateful to Gerry Smith for sharing the *rec8* mutant strain, and to Jeff Lange and Valeria Eliosa for technical support. This work was performed to fulfill, in part, requirements for MABN’s thesis research in the Graduate School of the Stowers Institute. Original data underlying this manuscript can be accessed from the Stowers Original Data Repository at http://www.stowers.org/research/publications/libpb-1514. This work was supported by the following awards to SEZ: The Stowers Institute for Medical Research (https://www.stowers.org), the March of Dimes Foundation Basil O’Connor Starter Scholar Research Award No. 5-FY18-58 (https://www.marchofdimes.org), the Searle Scholar Award, and the National Institutes of Health (NIH) under the award numbers R00GM114436 and DP2GM132936 (https://www.nih.gov). MABN was also supported by the National Cancer Institute of the NIH under award number F99CA234523. RLU was supported by funding from the University of Kansas. The funders had no role in study design, data collection and analysis, or manuscript preparation.

---

**Figure 1—figure supplement 1.**
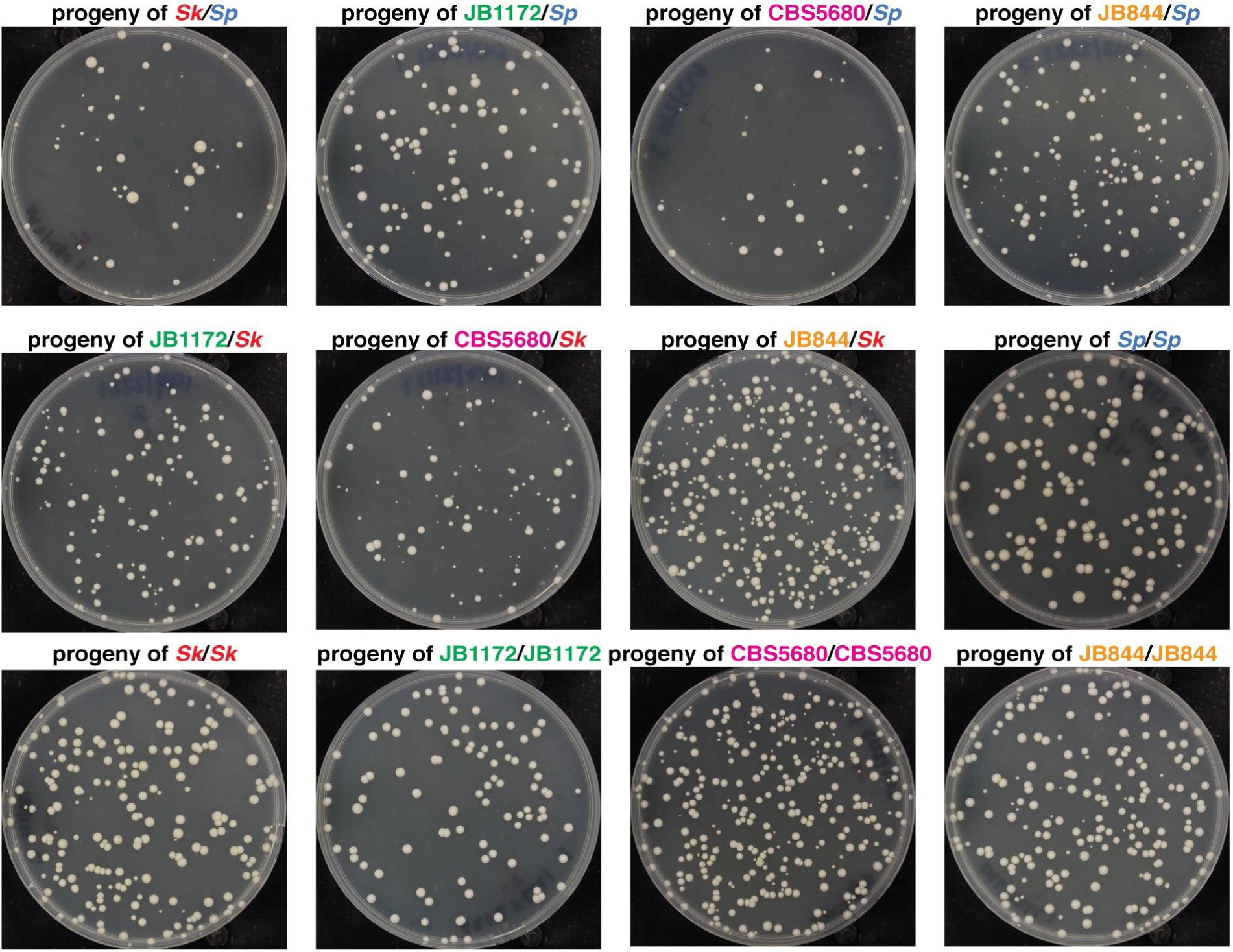
Colony phenotypes of spores produced by *S. pombe* heterozygous and homozygous diploids. Representative images of the spore colonies generated by the indicated diploids. *Sk/Sk,* JB1172/JB1172, and *Sk*/JB1172 images are the same as those presented in Figure 1C.

**Figure 1—figure supplement 2.**
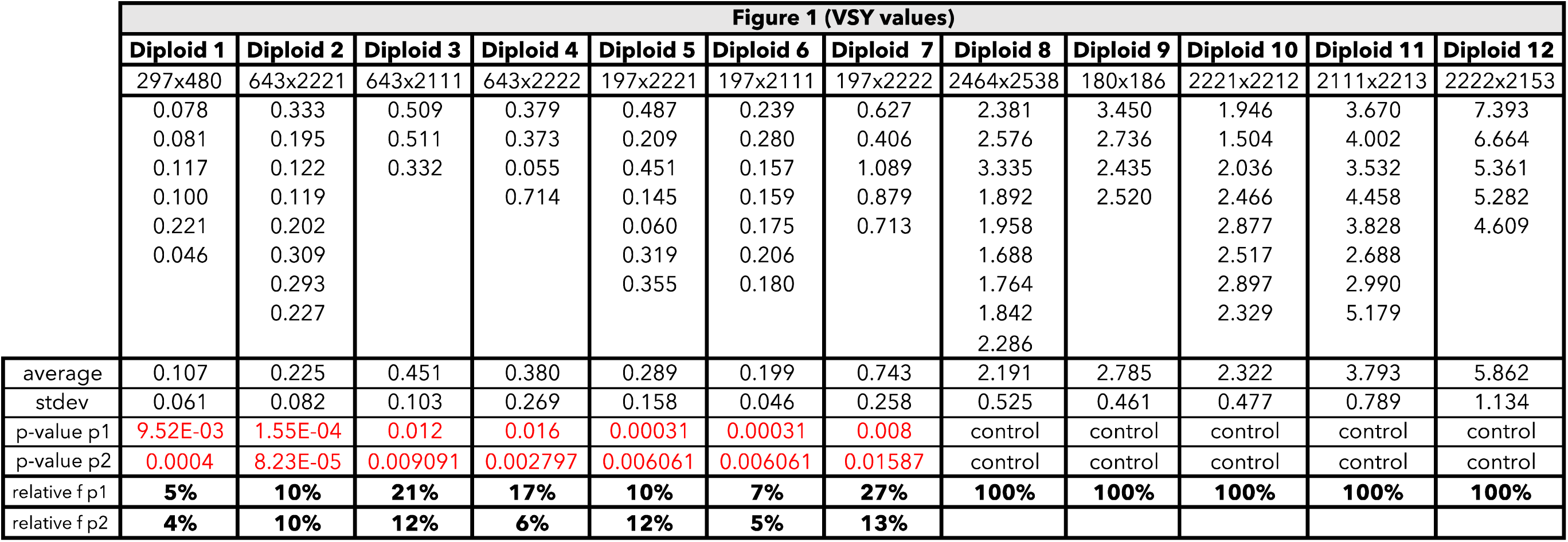
Raw data for the viable spore yield reported in Figure 1. Each column represents the diploid assayed, which matches the diploid number in Figure 1. The second row shows the diploid number. The third row shows the SZY strain numbers of both haploid parent strains. We present all the viable spore yield values from independent assays. We calculated the p-value using the Wilcoxon test by comparing the heterozygous diploid to the homozygous parent 1 (p1) and parent 2 (p2) strains. Diploid 1 was compared to control diploids 8 (p1) and 9 (p2); diploid 2 was compared to control diploids 8 (p1) and 10 (p2); diploid 3 was compared to control diploids 8 (p1) and 11 (p2); diploid 4 was compared to control diploids 8 (p1) and 12 (p2); diploid 5 was compared to control diploids 9 (p1) and 10 (p2); diploid 6 was compared to control diploids 9 (p1) and 11 (p2); and diploid 7 was compared to control diploids 9 (p1) and 12 (p2). The last two rows show the relative fertility (f) when compared to the homozygous parent 1 and parent 2.

**Figure 1—figure supplement 3.**
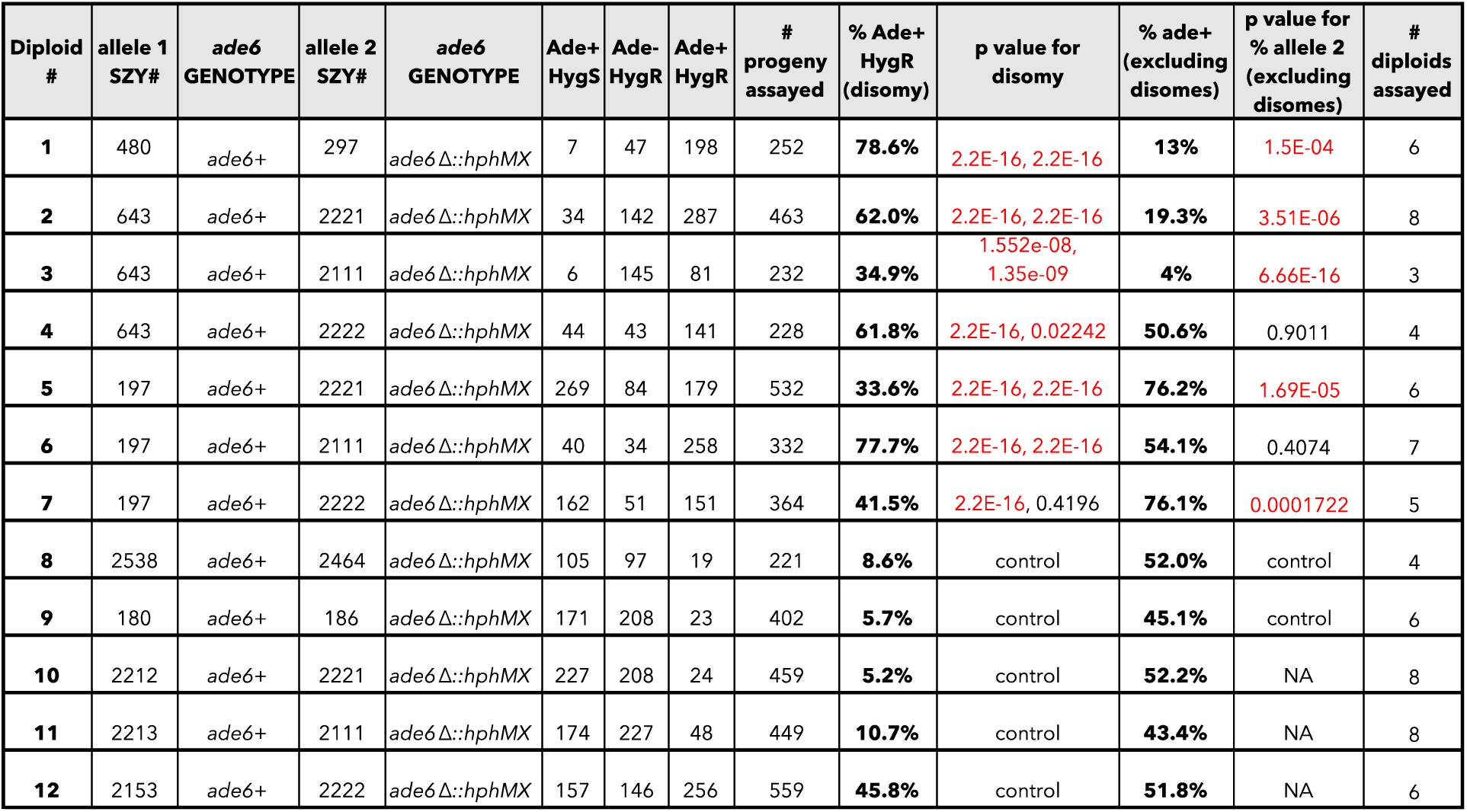
Raw data of allele transmission values reported in Figure 1. Each of the horizontal lines represents the relevant genotype and allele transmission of the indicated diploid. The first column matches the diploid number from Figure 1. Columns 2-5 contain the SZY strain number and relevant genotypes used to determine the allele transmission for chromosome 3. Columns 6-8 indicate the number of progeny that exhibited the indicated phenotype. The total number of progeny assayed is shown in column 9. Column 10 indicates the percentage of the progeny from column 8 that were likely disomes (Ade+ Hyg^R^). Column 11 indicates the p-values calculated when comparing the frequency of Ade+ Hyg^R^ progeny produced by heterozygous diploids to the frequency of Ade+ Hyg^R^ progeny produced by both homozygous diploid parent strains. Diploid 1 was compared to control diploids 8 and 9; diploid 2 was compared to control diploids 8 and 10; diploid 3 was compared to control diploids 8 and 11; diploid 4 was compared to control diploids 8 and 12; diploid 5 was compared to control diploids 9 and 10; diploid 6 was compared to control diploids 9 and 11; and diploid 7 was compared to control diploids 9 and 12. Column 12 shows the percentage of the progeny that were Ade+ Hyg^S^ (excluding Ade+ Hyg^R^ progeny). Column 13 indicates the p-value calculated when comparing diploids 2-4 to diploid 8 and diploids 5-7 to diploid 9. The last column shows the total number of independent diploids assayed for each cross.

**Figure 2—figure supplement 1.**
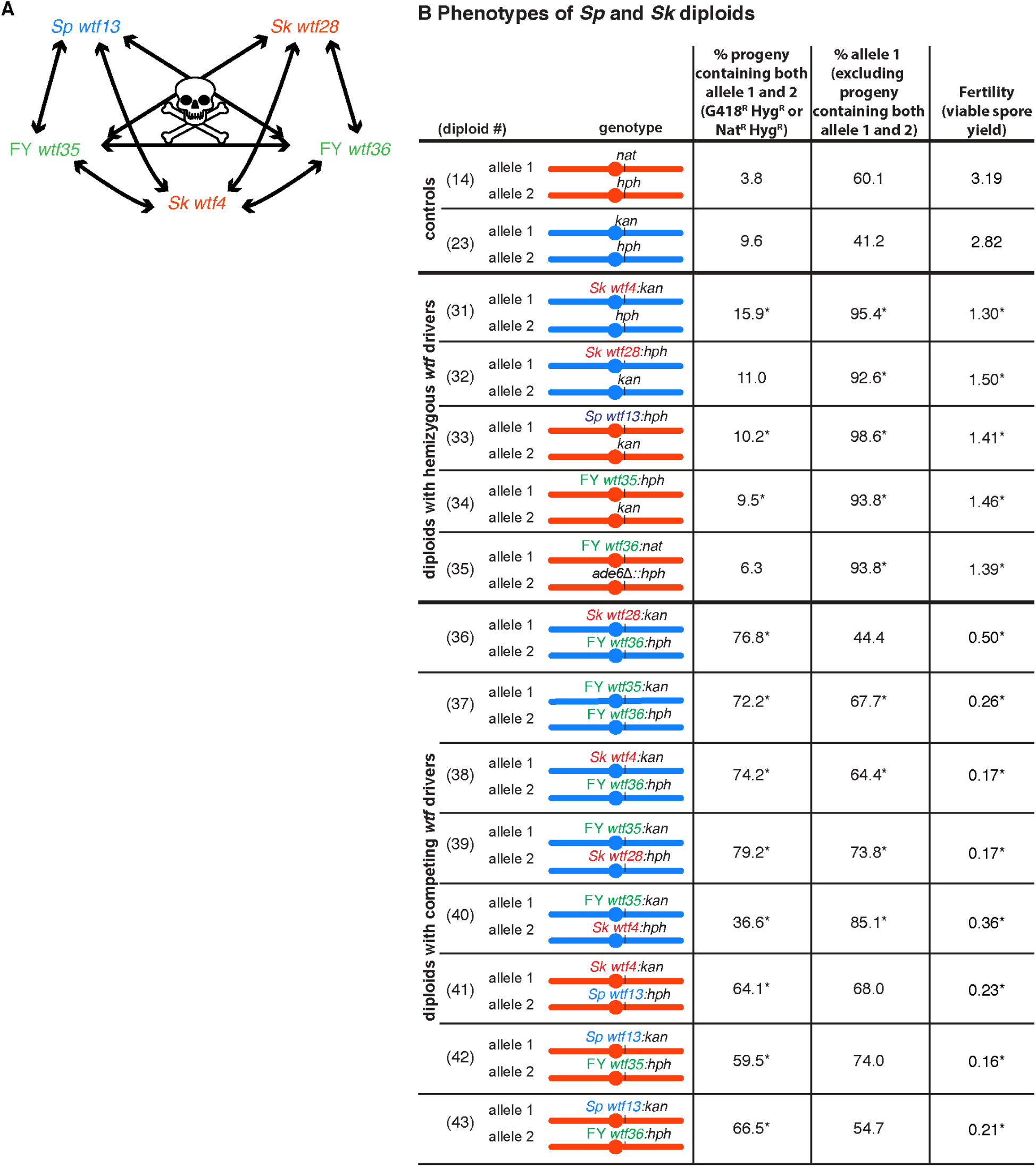
Wtf^antidote^ proteins are generally specific for a Wtf^poison^ and do not provide cross-resistance to other Wtf^poison^ proteins. (A) Summary of interactions between distinct *wtf* meiotic drivers. Due to the low fertility observed in the diploids with competing *wtf* meiotic drivers compared to the fertility of diploids with hemizygous *wtf* drivers, we concluded that the antidote of a given *wtf* driver does not provide resistance to a different poison. (B) Phenotypes of *Sp* and *Sk* diploids containing heterozygous *wtf* meiotic drivers or containing competing *wtf* meiotic drivers. For statistical analyses, the frequency of disomic progeny, allele transmission, and fertility in the diploids with heterozygous or competing *wtf* drivers were compared to the control diploids. Diploids 31, 32, and 36-40 were compared to control diploid 23, and diploids 33-35 and 41-43 were compared to control diploid 14. * indicates p-value <0.05 (G-test [allele transmission and Nat^R^ Hyg^R^ or G418^R^ Hyg^R^ progeny] and Wilcoxon test [fertility]). The data for diploids 14 and 23 were previously published in Bravo Núñez et al., 2020. The diploid numbers carry over between figures, meaning that the data for diploids 14 and 23 are repeated from Figure 2 and Figure 4, respectively. Raw data can be found in Figure 2—figure supplement 4 and Figure 2—figure supplement 5.

**Figure 2—figure supplement 2.**
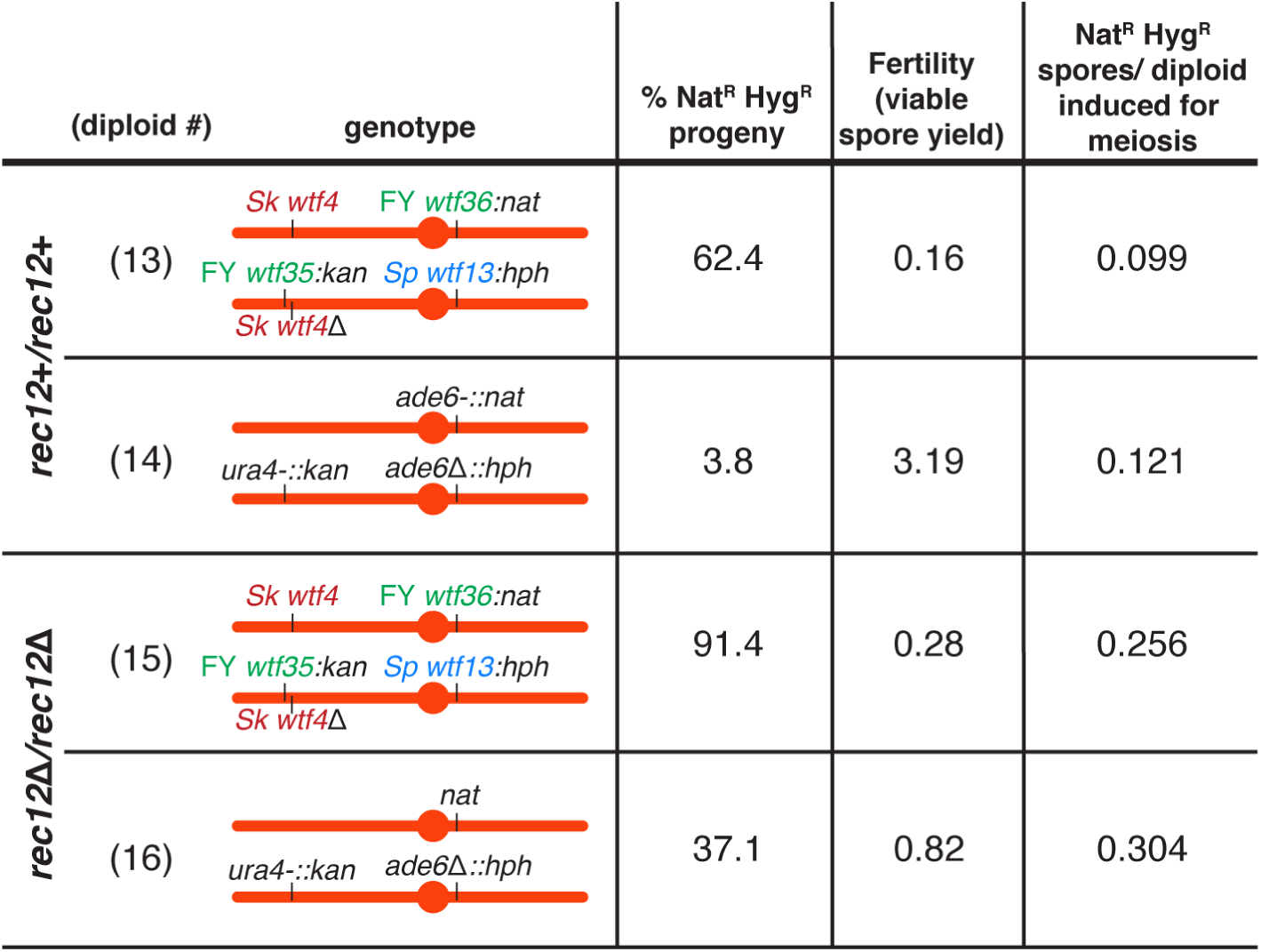
Competing *wtf* meiotic drivers do not affect the frequency of disomes generated per meiosis. To calculate the number of aneuploid and diploid gametes per meiosis from the diploids presented in Figure 2C, we multiplied the fraction of Nat^R^ Hyg^R^ progeny (disomes) by the viable spore yield (viable spores/viable diploid cells induced to undergo meiosis). This gave the number of Nat^R^ Hyg^R^ spores produced per diploid cell induced to undergo meiosis.

**Figure 2—figure supplement 3.**
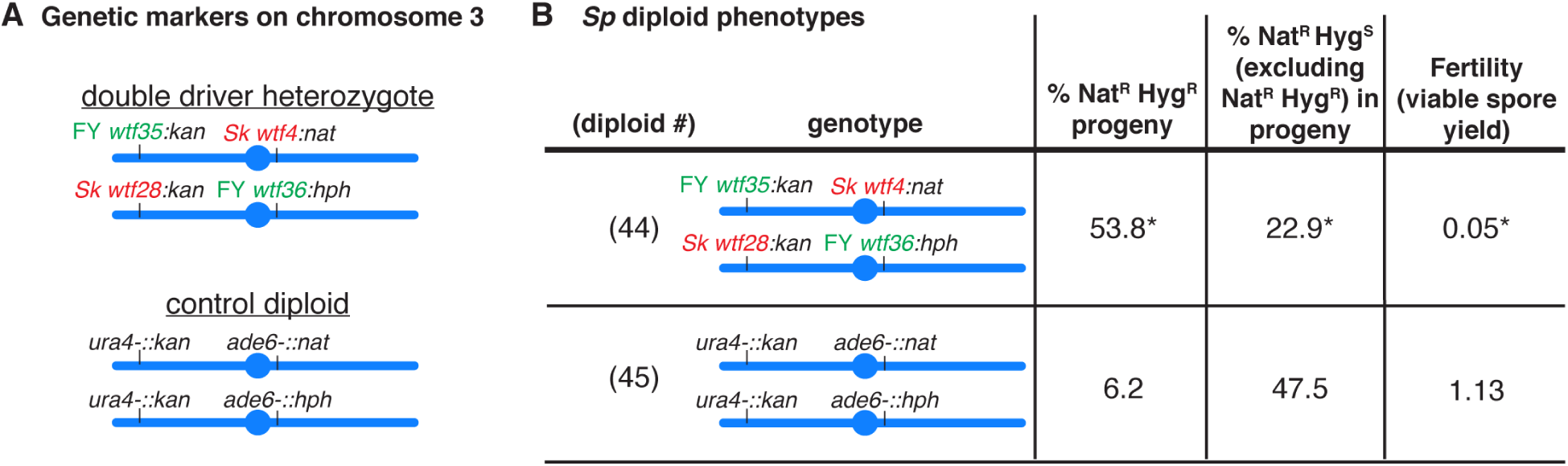
A high fraction of viable gametes are disomic in *Sp* strains with *wtf* competition at two loci. (A) Schematic of the genetic markers utilized in the control diploid and the double driver heterozygote in the *Sp* strain background. (D) Allele transmission and fertility of *Sp* diploids is shown. The allele transmission at *ade6* was determined using co-dominant markers (*natMX4* and *hphMX6*). For statistical analysis, the frequency of disomic progeny, allele transmission, and fertility from diploid 44 was compared to control diploid 45. * indicates p-value <0.05 (G-test [% NAT^R^ Hyg^R^ progeny and allele transmission] and Wilcoxon test [fertility]). Raw data can be found in Figure 2—figure supplement 4 and Figure 2—figure supplement 5. The low VSY of diploid 45 relative to diploid 8 is likely due to double auxotrophy of this strain and the fact that all the experiments used unsupplemented SPA media (see methods).

**Figure 2—figure supplement 4.**
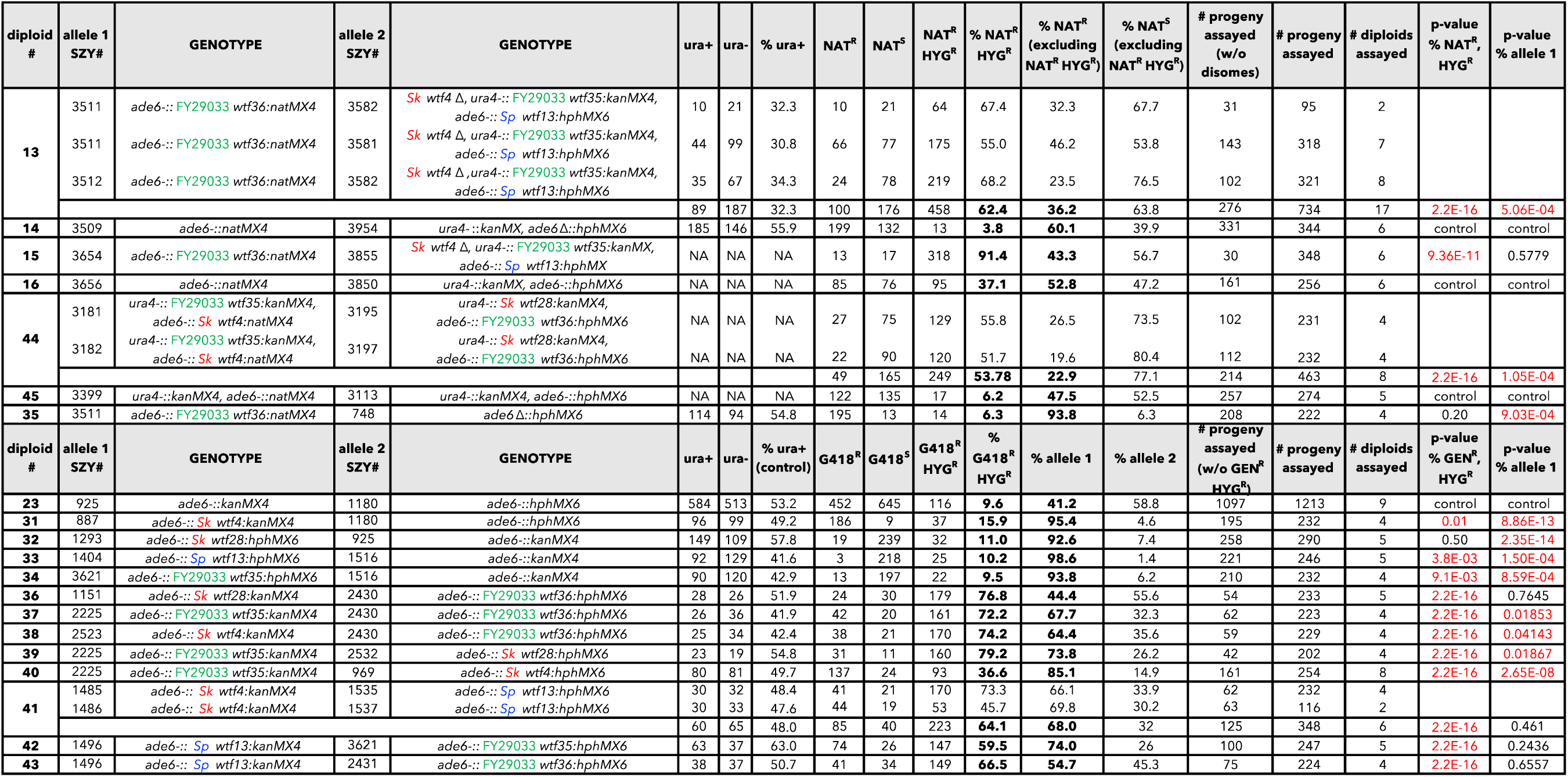
Raw data of allele transmission values reported in Figure 2, Figure 2—figure supplement 1, and Figure 2—figure supplement 3. Each of the horizontal lines represents the relevant genotype and allele transmission of the indicated diploid. The first column matches the diploid number from Figure 2, Figure 2—figure supplement 1, and Figure 2—figure supplement 3. In columns 2-5 are the SZY strain number and relevant genotypes used to determine the allele transmission for chromosome 3. Columns 6 and 7 show the indicated genotype (ura+ or ura- progeny excluding disomes). Columns 9-11 indicate the number of haploid progeny that exhibited the indicated phenotype. Column 15 indicates the total number of progeny assayed excluding Nat^R^ Hyg^R^ and G418^R^ Hyg^R^ progeny. Column 16 indicates the total number of progeny assayed. Column 17 shows the total number of independent diploids assayed for each cross. Columns 18 and 19 indicate the p-values calculated with a G-test when comparing diploids 13, 33-35, and 41-43 to control diploid 14; diploid 15 to control diploid 16; diploid 44 to control diploid 45; and diploids 31, 32, and 36-40 to control diploid 23. The diploid numbers carry over between figures, meaning that the raw data for diploid 23 is also presented in Figure 4—figure supplement 5 and Figure 6—figure supplement 3. The data for diploids 14 and 23 were previously published in Bravo Núñez et al., 2020.

**Figure 2—figure supplement 5.**
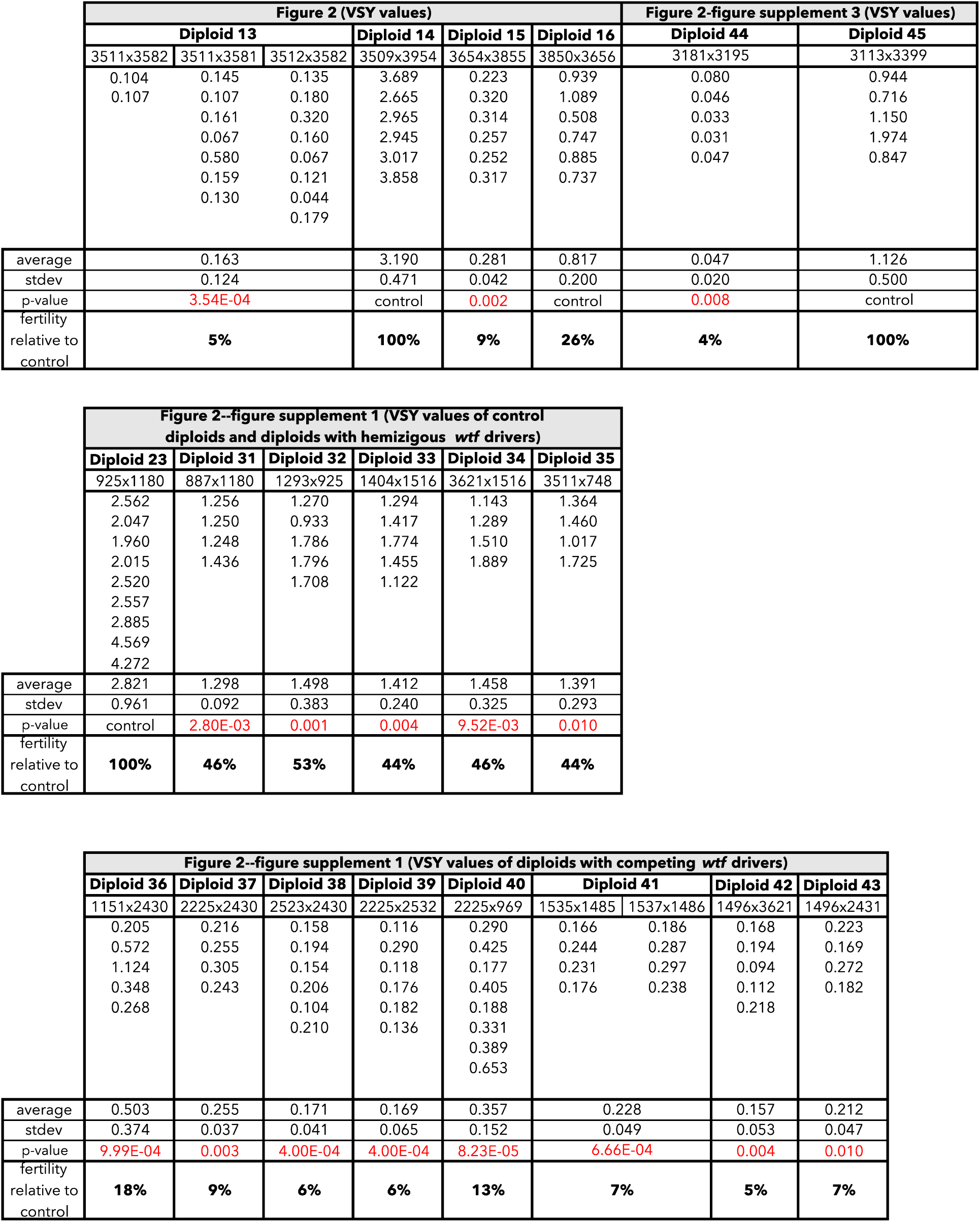
Raw data of viable spore yield assays reported in Figure 2, Figure 2—figure supplement 1, and Figure 2—figure supplement 3. Each column represents the diploid assayed, which matches the diploid number in Figure 2, Figure 2—figure supplement 1, and Figure 2—figure supplement 3. The first row indicates in which figure these data are reported. The second row shows the diploid number. The third row shows the SZY strain numbers of both haploid parent strains. We present all of the viable spore yield values from independent assays. We calculated the p-value using the Wilcoxon test by comparing diploids 13, 33-35, and 41-43 to control diploid 14; diploid 15 to control diploid 16; diploid 44 to control diploid 45; and diploids 31, 32, and 36-40 to control diploid 23. The diploid numbers carry over between figures, meaning that the raw data for diploid 23 is also presented in Figure 4—figure supplement 6 and Figure 6—figure supplement 4. The data for diploids 14 and 23 were previously published in Bravo Núñez et al., 2020.

**Figure 2—figure supplement 6.**
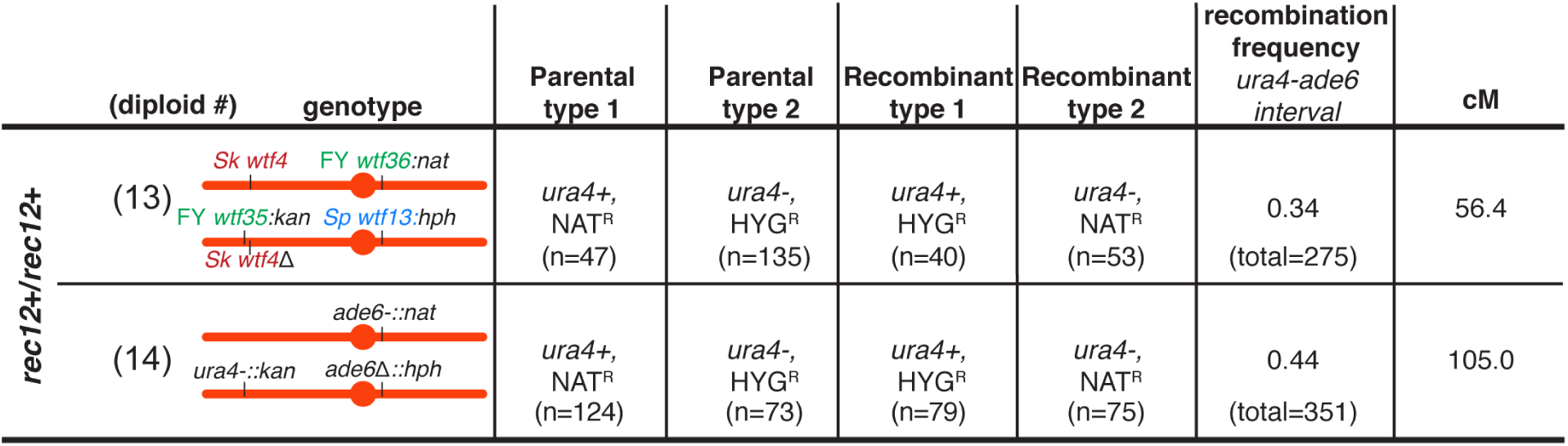
Recombination frequency is altered in *Sk* diploids with competing *wtf* meiotic drivers. To determine the recombination frequency (R) within the *ade6* and *ura4* interval in diploids 13 and 14 presented in Figure 2, we calculated the number of recombinants/(number of parental and recombinant progeny). To calculate the genetic distance (cM), we used Haldane’s formula x= −50 ln(1-2R) (Haldane 1919; Smith 2009).

**Figure 3—figure supplement 1.**
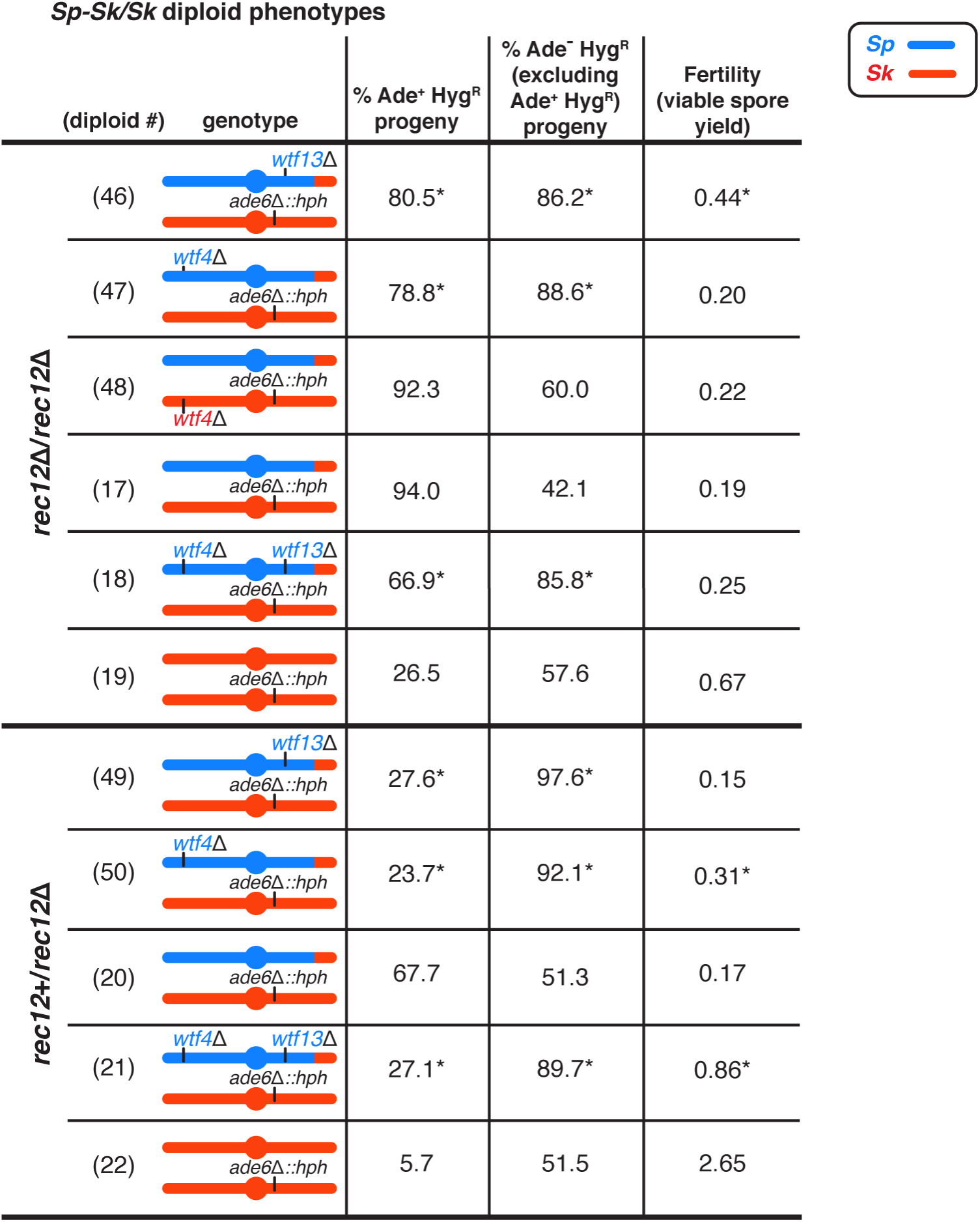
Single deletions of *Sp wtf* meiotic drivers partially decrease the frequency of disomic gametes in *Sp/Sk* mosaic diploid. Phenotypes of *Sp/Sk* mosaic and control diploids in *rec12+*/*rec12Δ* and *rec12Δ*/*rec12Δ* backgrounds. Allele transmission of chromosome 3 was assayed using co-dominant markers at *ade6* (*ade6+* and *ade6Δ::hphMX6*). For statistical analyses of disomy and fertility, we compared diploids 18 and 46-48 to diploid 17, and diploids 21, 49, and 50 to diploid 20. For statistical analyses of allele transmission, we compared diploids 17, 18, and 46-48 to control diploid 19, and we compared diploids 20, 21, 49, and 50 to control diploid 22. * indicates p-value <0.05 (G-test [allele transmission, Ade+ Hyg^R^] and Wilcoxon test [fertility]). Raw data can be found in Figure 3—figure supplement 3 and Figure 3—figure supplement 4. Diploid numbers get carried over between figures, meaning that the data for diploids 17-22 are also presented in Figure 3.

**Figure 3—figure supplement 2.**
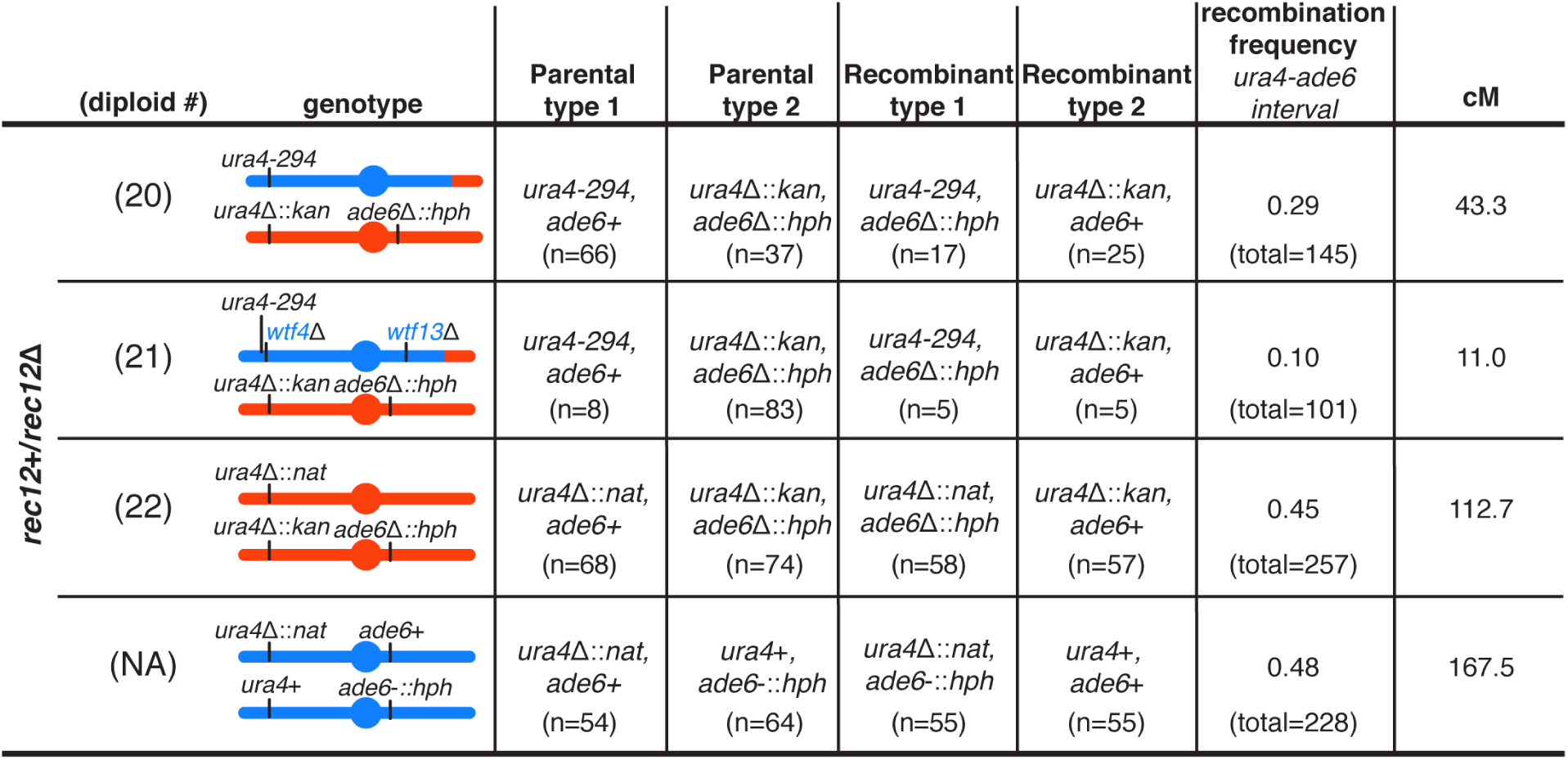
Observed recombination frequencies are altered by meiotic drive in *Sk/Sp* mosaic diploid. To determine the recombination frequencies (R) between *ade6* and *ura4* in Rec12+ diploids presented in Figure 3 (diploids 20, 21, and 22), and in an *Sp* homozygous control, we calculated the number of recombinant progeny/(number of parental and recombinant progeny). To calculate the genetic distance (cM), we used Haldane’s formula x= −50 ln(1-2R) (Haldane 1919; Smith 2009). The data for the *Sp* homozygous control was generated by crossing strains SZY2397 and SZY1180.

**Figure 3—figure supplement 3.**
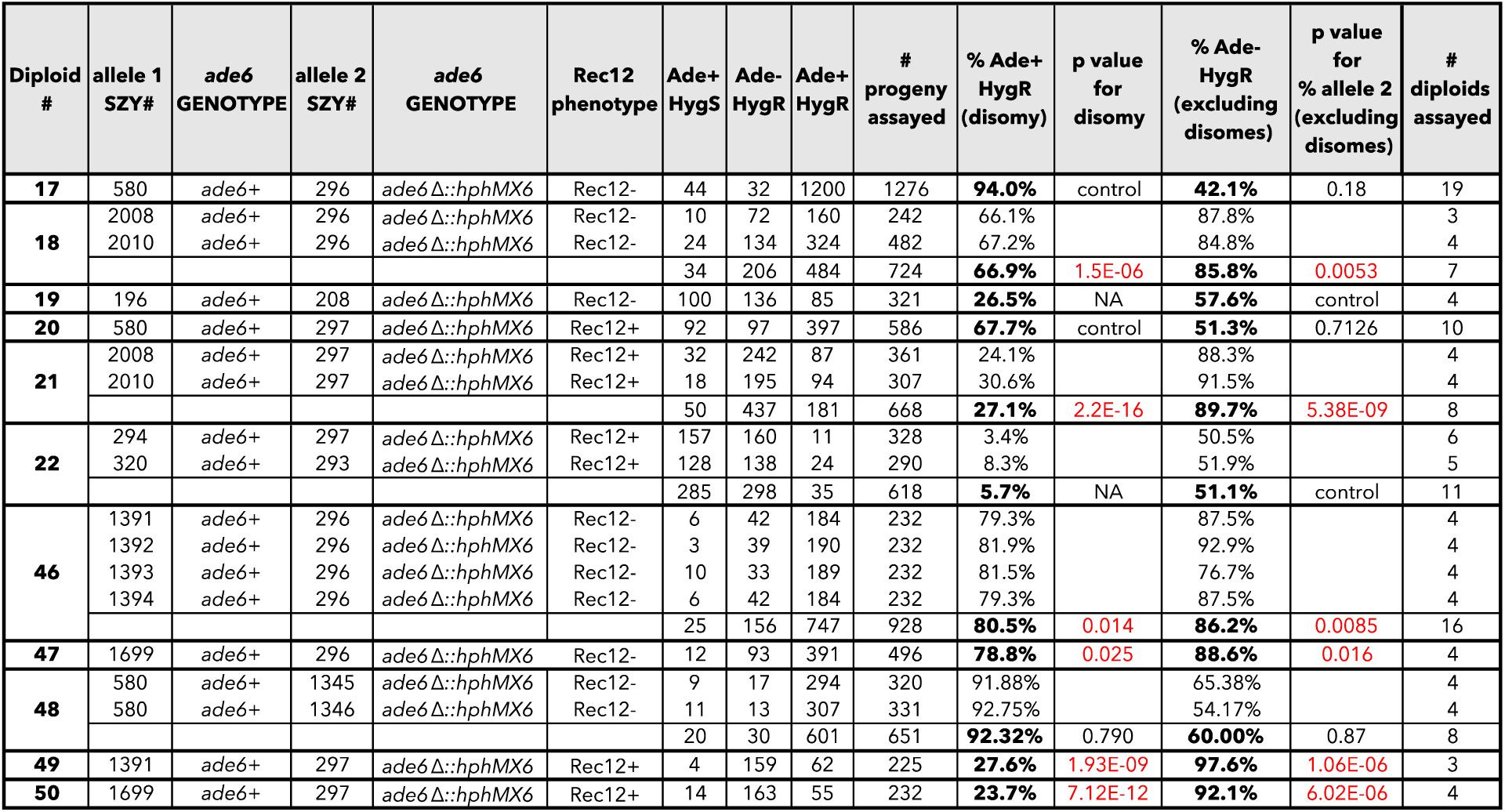
Raw data of allele transmission values reported in Figure 3 and Figure 3—figure supplement 1. Each of the horizontal lines represents the relevant genotype and allele transmission of the indicated diploid. The first column matches the diploid number from Figure 3 and Figure 3—figure supplement 1. In columns 2-5 are the SZY strain number and relevant genotypes used to determine the allele transmission for chromosome 3 in the *Sp/Sk* mosaic and control diploids. Column 6 shows the Rec12 phenotype for each diploid. Columns 7 and 8 indicate the number of progeny that exhibited the indicated phenotype (Ade+ Hyg^S^ or Ade-Hyg^R^, respectively). Column 9 shows the number of progeny that exhibited both phenotypes and is thus Ade+ and Hyg^R^. These progeny are likely disomic. The total number of progeny assayed is shown in column 10. Column 11 indicates the percentage of the progeny from column 9 that were both Ade+ and Hyg^R^. Column 12 indicates the p-values calculated when comparing diploids 18, 46, 47, and 48 to diploid 17, and diploids 21, 49, and 50 to diploid 20. Column 13 shows the percentage of the progeny (excluding Ade+ Hyg^R^ progeny) that were Ade-Hyg^R^. Column 14 indicates the p-value calculated when comparing diploids 17, 18, and 46-48 to control diploid 19, and diploids 20, 21, 49, and 50 to control diploid 22. The last column shows the total number of independent diploids assayed for each cross.

**Figure 3—figure supplement 4.**
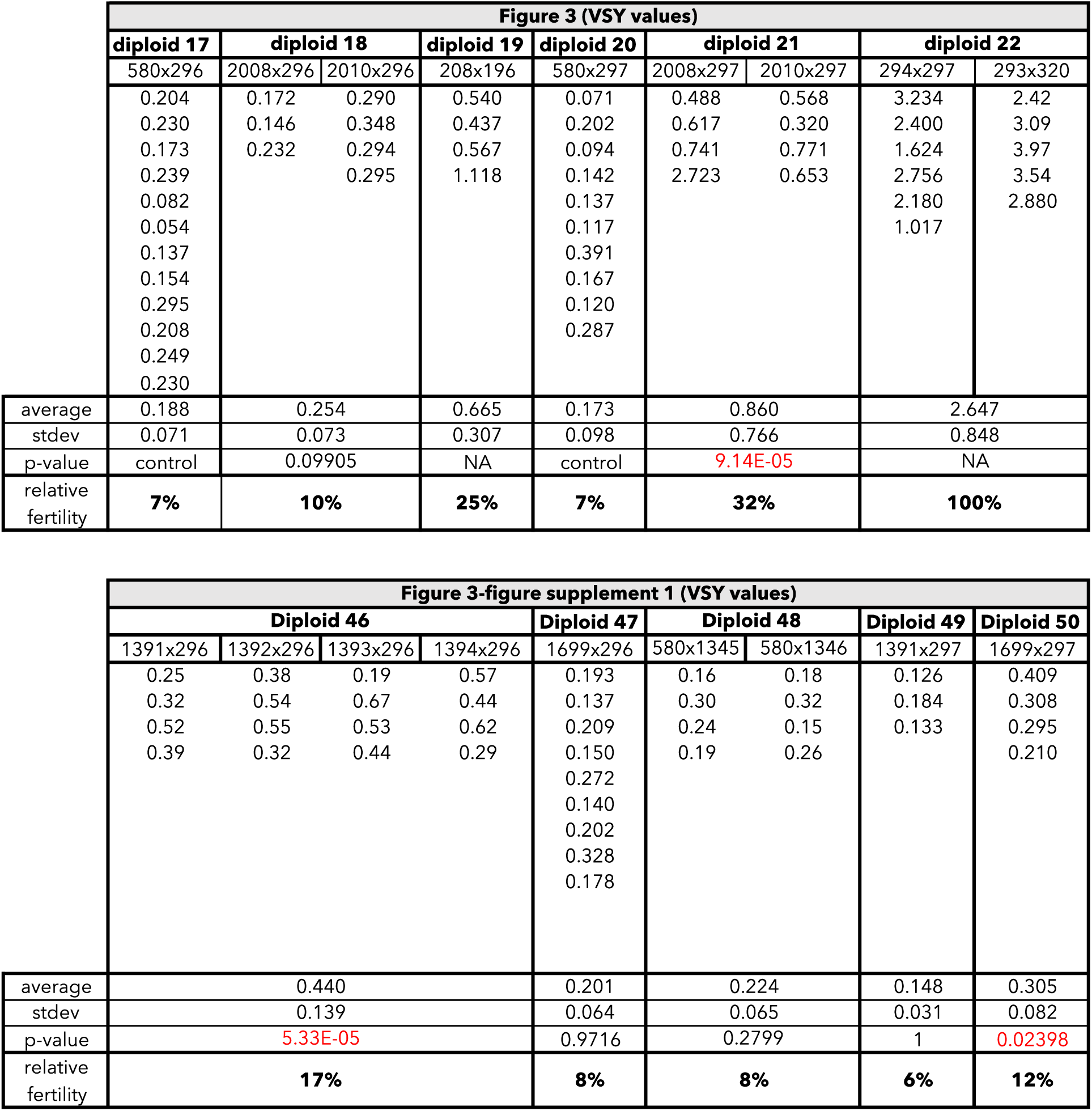
Raw data of viable spore yield reported in Figure 3 and Figure3—figure supplement 1. The top table shows the data for Figure 3. The bottom table shows the data for Figure 3—figure supplement 1. Each column represents the diploid assayed, which matches the diploid numbers in Figure 3 and Figure 3—figure supplement 1. The first row shows the figure where the data are reported. The second row shows the diploid number. The third row shows the SZY strain numbers of both haploid parent strains. We present all of the viable spore yield values from independent assays. We calculated the p-value using the Wilcoxon test by comparing diploids 18, 46, 47, and 48 to diploid 17, and diploids 21, 49, and 50 to diploid 20.

**Figure 4—figure supplement 1.**
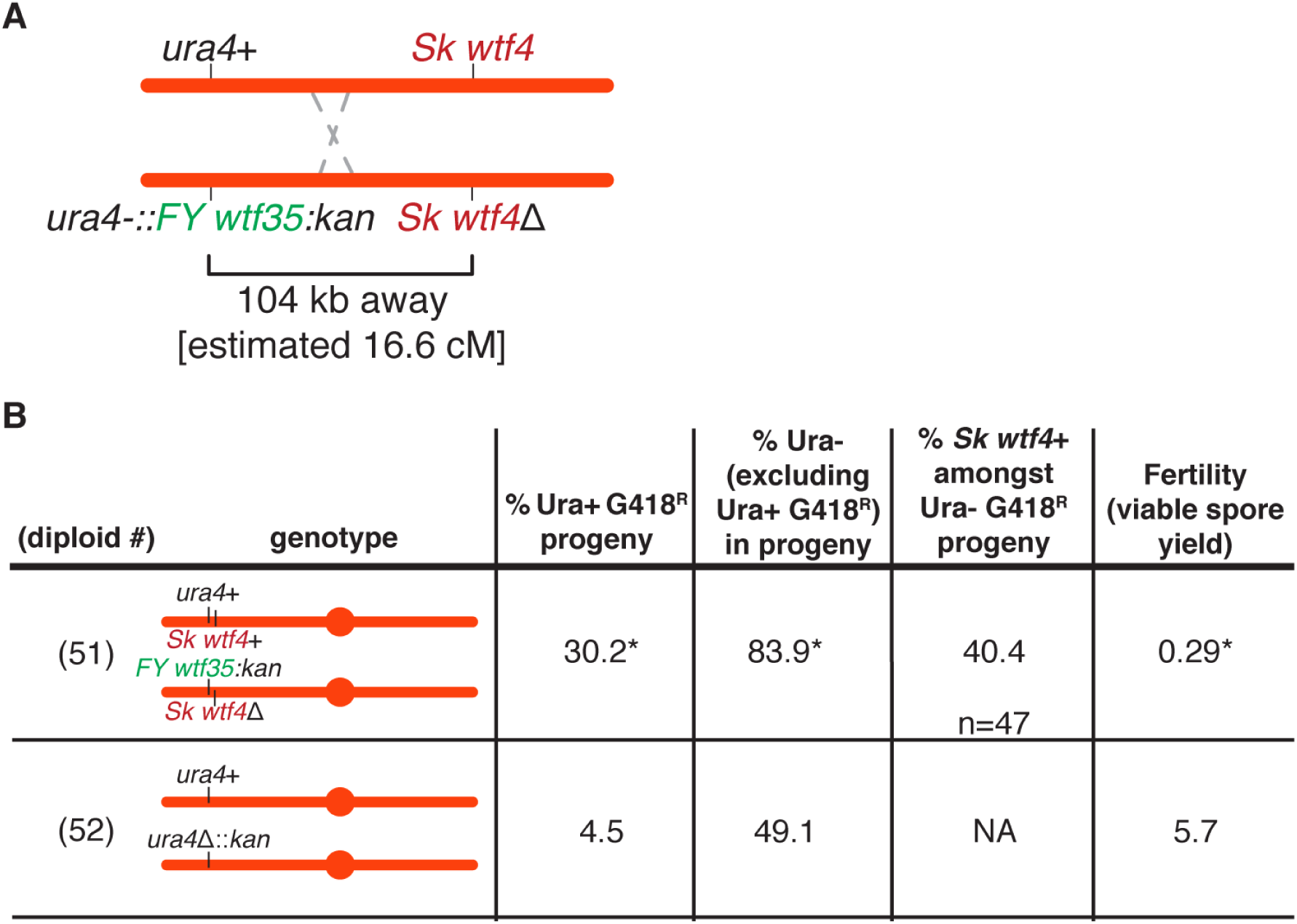
Competing *wtf* meiotic drivers enrich for spores that contain both *wtf* alleles in *Sk.* Phenotypes of *wtf* competition and control diploids. Alleles were followed using the *ura4+* and *kanMX4* markers. We determined the presence of the *wtf4+* allele amongst the ura- G418^R^ progeny via PCR with oligos 543+548. For statistical analyses of allele transmission, ura+ G418^R^ frequency, and fertility, we compared diploid 51 to control diploid 52. * indicates p-value < 0.05 (allele transmission and ura+ G418^R^ [G-test], fertility [Wilcoxon test]). The raw data can be found in Figure 4—figure supplement 5 and Figure 4—figure supplement 6.

**Figure 4—figure supplement 2.**
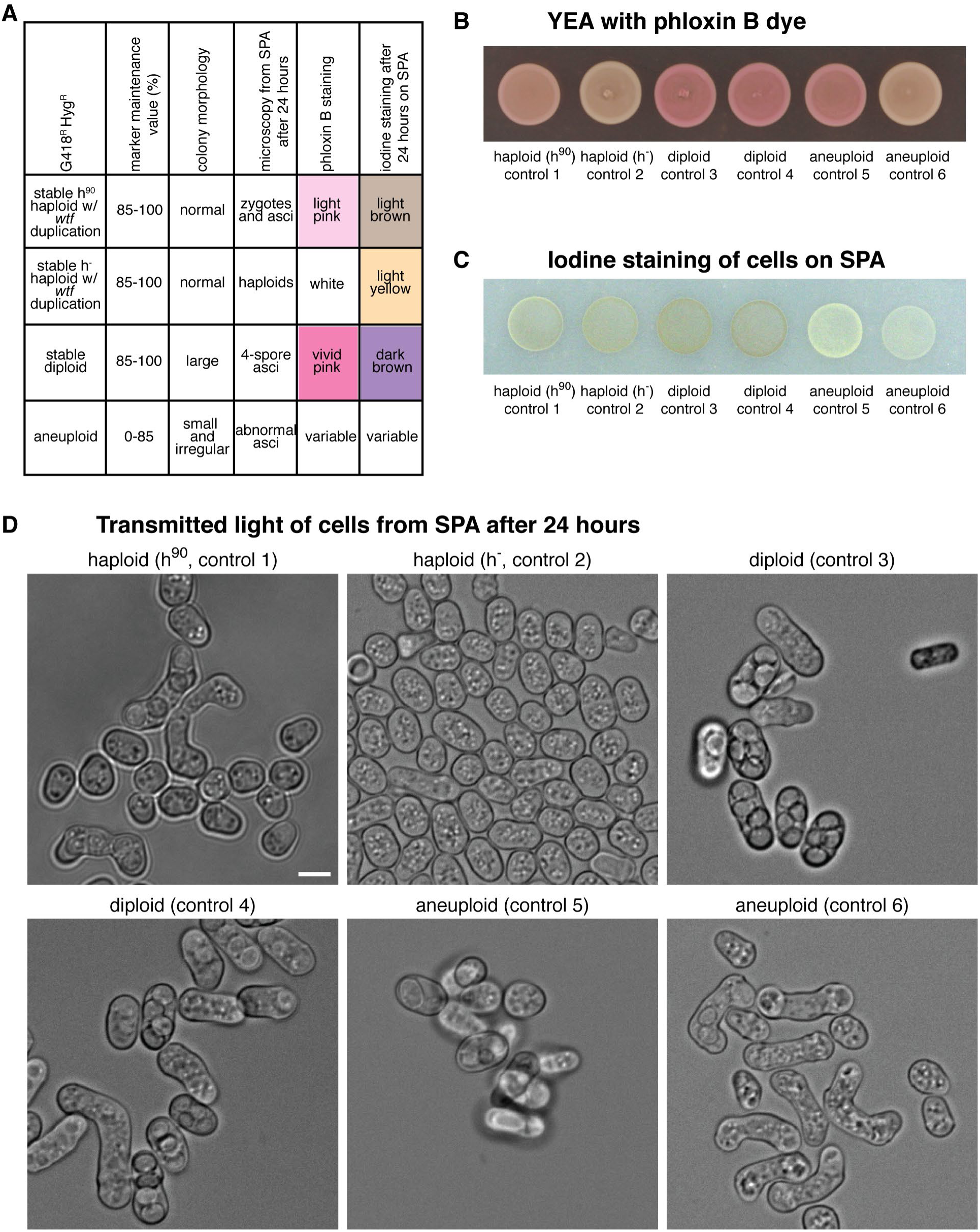
Methods to distinguish between haploid, diploid, and aneuploid spore colonies. (A) Summary table of how the ploidy of the spore colonies was determined based on the distinct phenotypes is shown. The marker maintenance value reflects the percentage of the population that showed the presence of the two markers (*kanMX4* and *hphMX6*) after passaging 1:10 diluted 650 μL cultures for 5 days. (B) Images of the controls spotted onto phloxin B-containing plates. Phloxin B is normally used in *S. pombe* to stain dead cells (Forsburg and Rhind 2006). Due to the instability of *S. pombe* diploids, patches of diploid cells stain bright pink. In contrast, haploid cells do not stain and remain white. (C) Images of the controls spotted onto sporulation plates (SPA) and then stained with iodine vapors after 24 hours. Iodine vapors stain the starch in the spore wall (Forsburg and Rhind 2006). Cells that made spores after 24 hours on sporulation plates, will stain brown. Otherwise, the cells will stain yellow. (D) Representative images of the haploid, diploid, and aneuploid controls. The cells were taken from the sporulation plates after 24 hours. Scale bar represents 5 μm.

**Figure 4—figure supplement 3.**
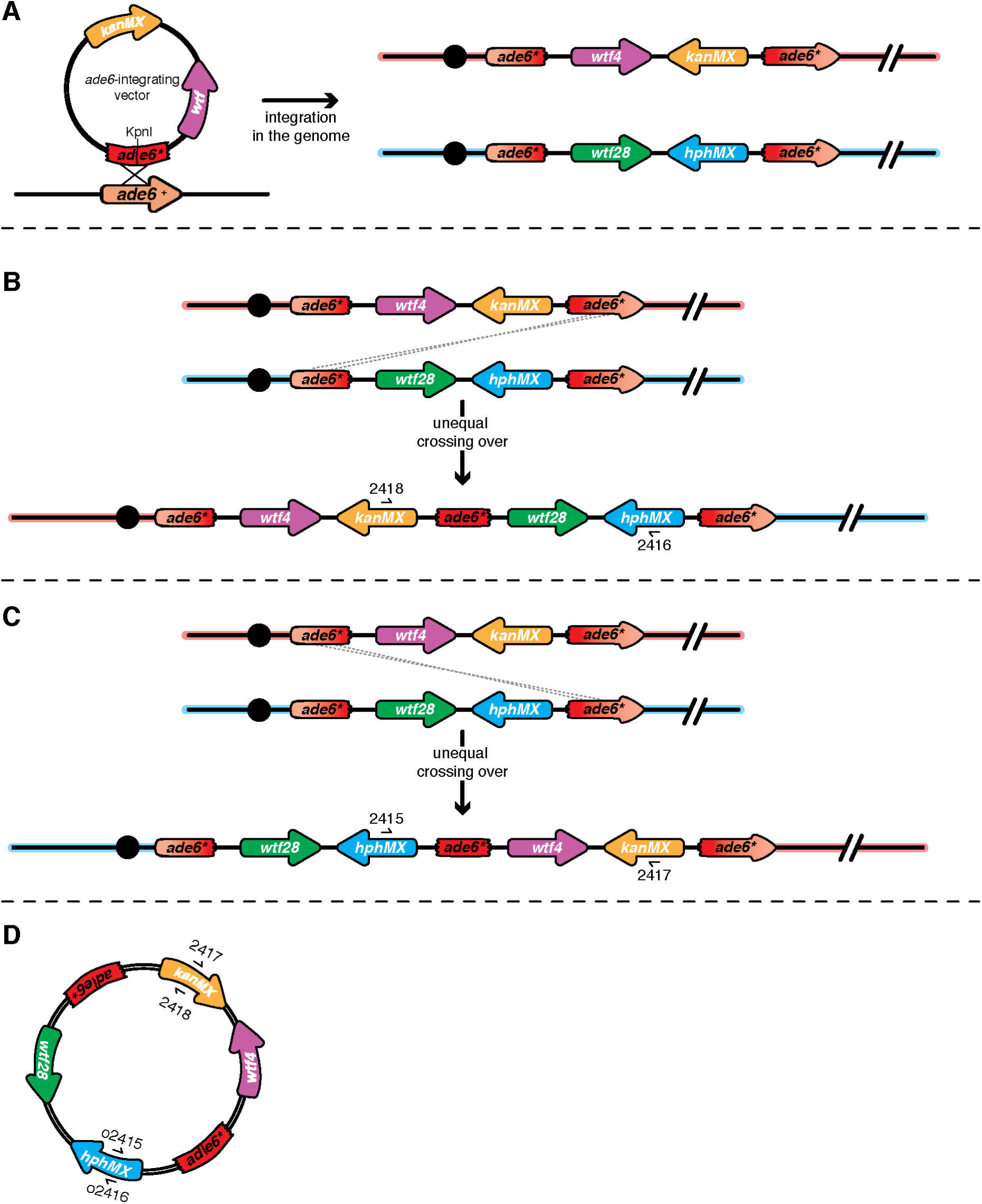
Unequal crossover event at *ade6* leads to *wtf* duplication. (A) Schematic of the *ade6* locus before and after transformation of the *ade6-*integrating vector into the genome. (B) An example of unequal crossing over leading to a *wtf* duplication event is shown. To determine the presence of this duplication event in the G418^R^ Hyg^R^ haploids, we used oligos 2416 + 2418 found within the drug resistance cassettes. (C) An example of a second scenario of unequal crossing over leading to a *wtf* duplication is shown. To determine the presence of the duplication event in the G418^R^ Hyg^R^ haploids, we used oligos (2415 + 2417) within the drug resistance markers. (D) Schematic of circularization of duplicated *ade6* transgenes. The *ade6-*integration vectors are prone to popping-out due to intrachromatid crossovers between the terminal *ade6* cassettes. After the duplication event happened, the entire construct could have popped-out of the genome. The PCR results (i.e. amplification of a product with both pairs of oligos presented in Figure 4—figure supplement 4) are consistent with these pop-outs forming in most cases.

**Figure 4—figure supplement 4.**
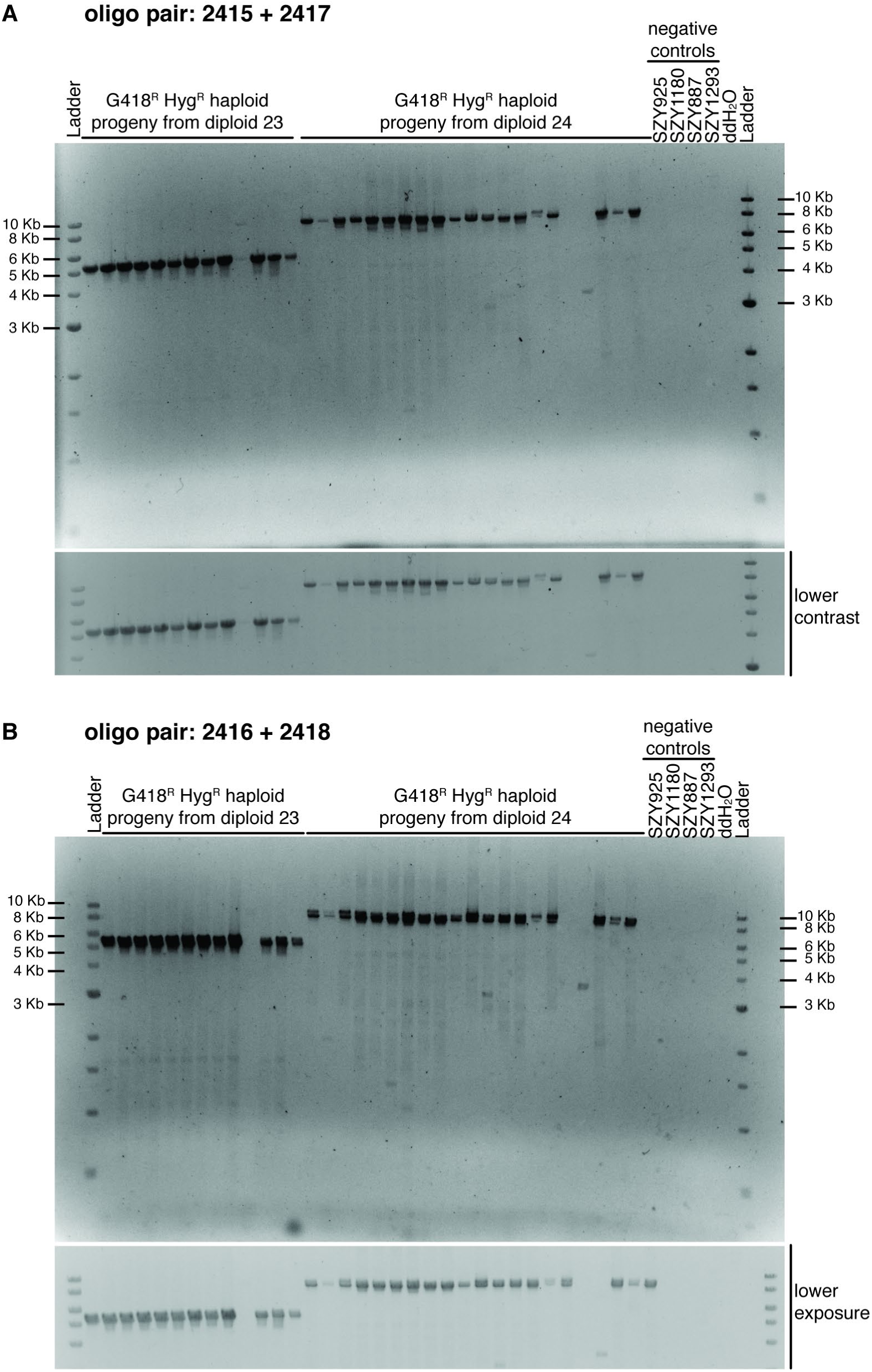
Most haploids with duplication events show evidence of unequal crossover scenarios. Ethidium bromide agarose gel (1%) of predicted haploids with duplication events using (A) oligos 2415 + 2417 and (B) oligos 2416 + 2418. Most of the haploids tested show both orientations of duplication events suggesting potential circularization of the duplicated transgenes as shown in Figure 4—figure supplement 3D. Below each gel, we show an image with lower contrast or exposure, respectively. The negative controls are the haploid parental strains.

**Figure 4—figure supplement 5.**
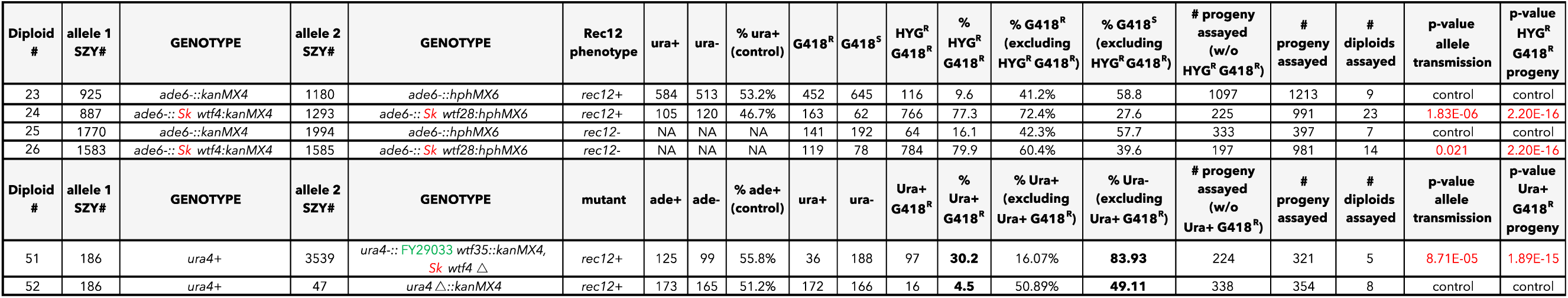
Raw data of allele transmission values reported in Figure 4C and Figure 4—figure supplement 1. Each of the horizontal lines represents the relevant genotype and allele transmission of the indicated diploid. The first column matches the diploid number from Figure 4C and Figure 4—figure supplement 1. In columns 2-5 are the SZY strain number and relevant genotypes used to determine the allele transmission of the *wtf* genes or empty vector. Column 6 indicates if the diploid has a *rec12+* or *rec12-* phenotype. Columns 7 and 8 indicate the indicated genotype (ura+ or ura- progeny). Columns 10 and 11 indicate the number of haploid progeny that exhibited the indicated phenotype (G418^R^ or G418^S^). Column 12 shows the number of progeny that exhibited the G418^R^ Hyg^R^ phenotype. Column 16 indicates the total number of progeny assayed excluding G418^R^ Hyg^R^ progeny. Column 17 indicates the total number of progeny assayed. Column 18 shows the total number of independent diploids assayed for each genotype. The last two columns show the p-values calculated using a G-test from the allele transmission at *ade6* and the frequency of G418^R^ Hyg^R^ progeny, respectively. To calculate the p-values, we compared diploid 24 to control diploid 23; diploid 26 to control 25; and diploid 51 to control diploid 52. The data for diploid 23 were previously published in Bravo Núñez et al., 2020. The raw data for diploid 23 is also presented in Figure 2—figure supplement 4 and Figure 6—figure supplement 3.

**Figure 4—figure supplement 6.**
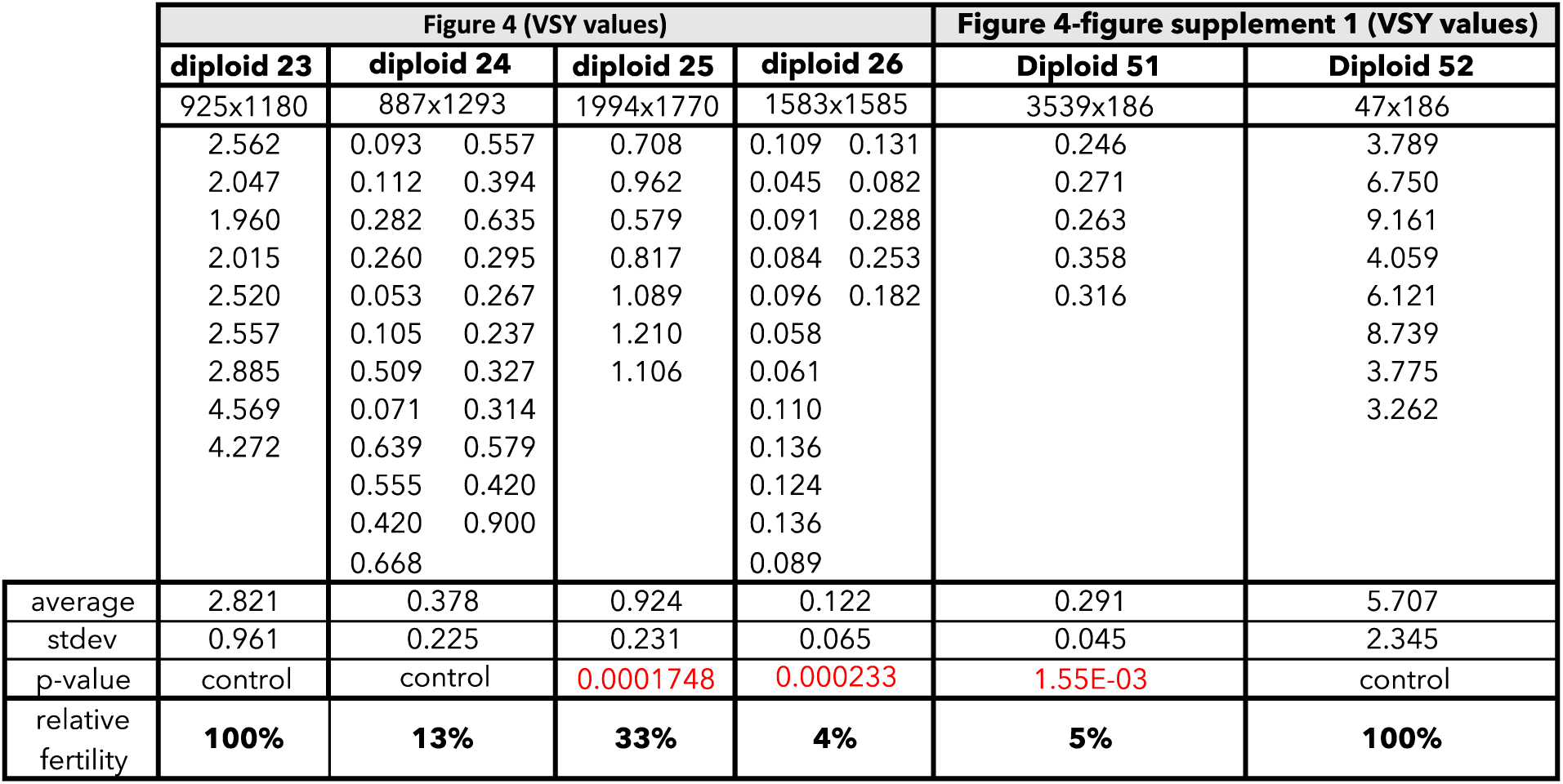
Raw data of viable spore yield reported in Figure 4 and Figure 4—figure supplement 1. Each column represents the diploid assayed, which matches the diploid number in Figure 4 and Figure 4—figure supplement 1. The first row shows the figure where the data are presented. The second row shows the diploid number. The third row shows the SZY strain numbers of both haploid parent strains. We present all of the viable spore yield values from independent assays. To calculate the p-value, we used the Wilcoxon test and compared diploid 24 to control diploid 23; diploid 26 to control diploid 25; and diploid 51 to control diploid 52. The data for diploid 23 were previously published in Bravo Núñez et al., 2020. The raw data for diploid 23 are also presented in Figure 2—figure supplement 5 and Figure 6— figure supplement 4.

**Figure 5—figure supplement 1.**
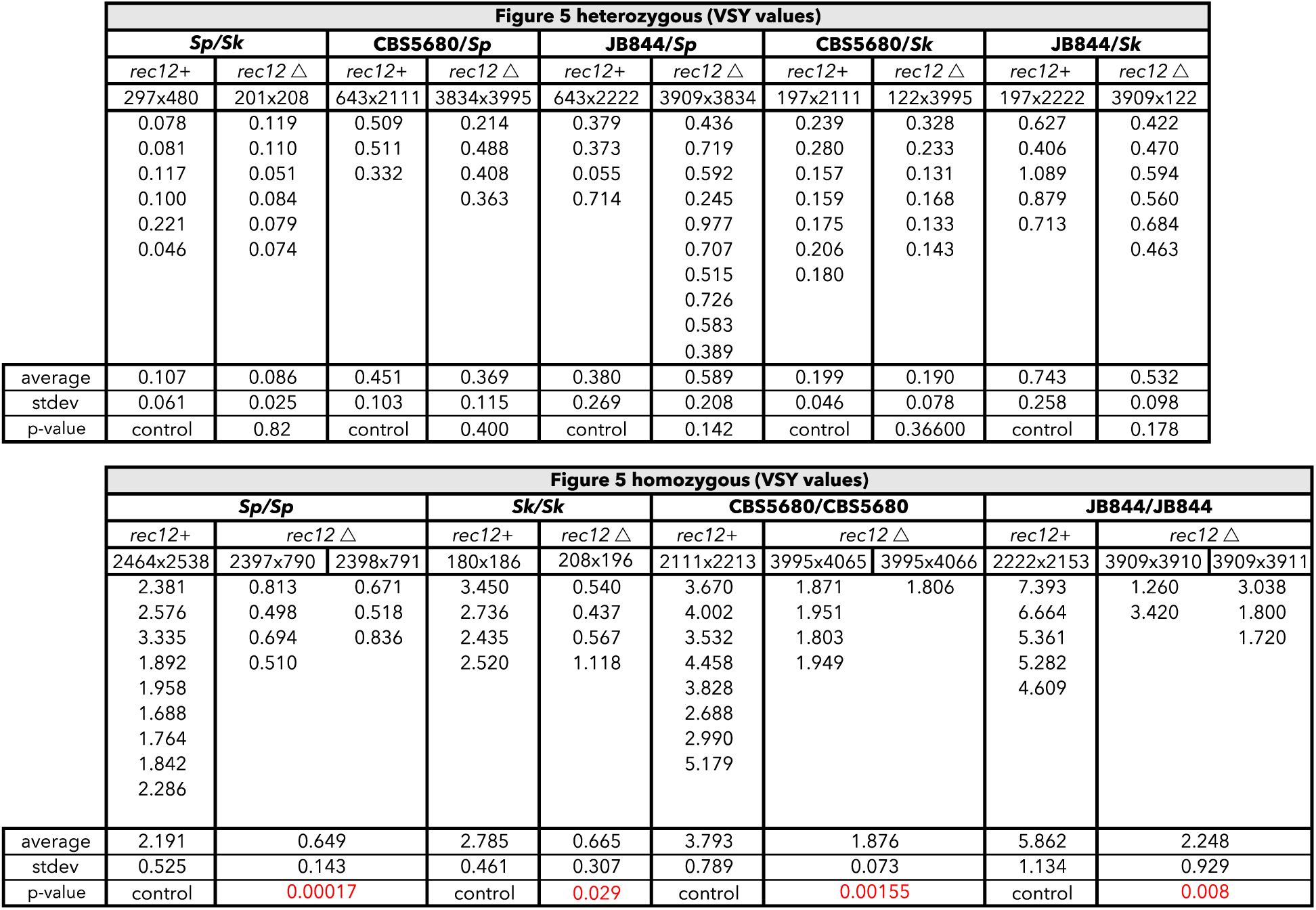
Raw data of the viable spore yield reported in Figure 5. The top columns represent the *S. pombe* diploid assayed and the backgrounds in which it was tested (*rec12+* or *rec12*Δ). The SZY strain numbers of both of the haploid parental strain are shown underneath the *rec12* genotype. We present all of the viable spore yield values from independent assays. For statistical analyses, in every case, the *rec12*Δ diploid was compared to the *rec12+* diploid. The raw data from the *rec12+* crosses are also shown in Figure 1—figure supplement 2. The raw data for the *Sk rec12*Δ is also presented in Figure 3—figure supplement 4.

**Figure 6—figure supplement 1.**
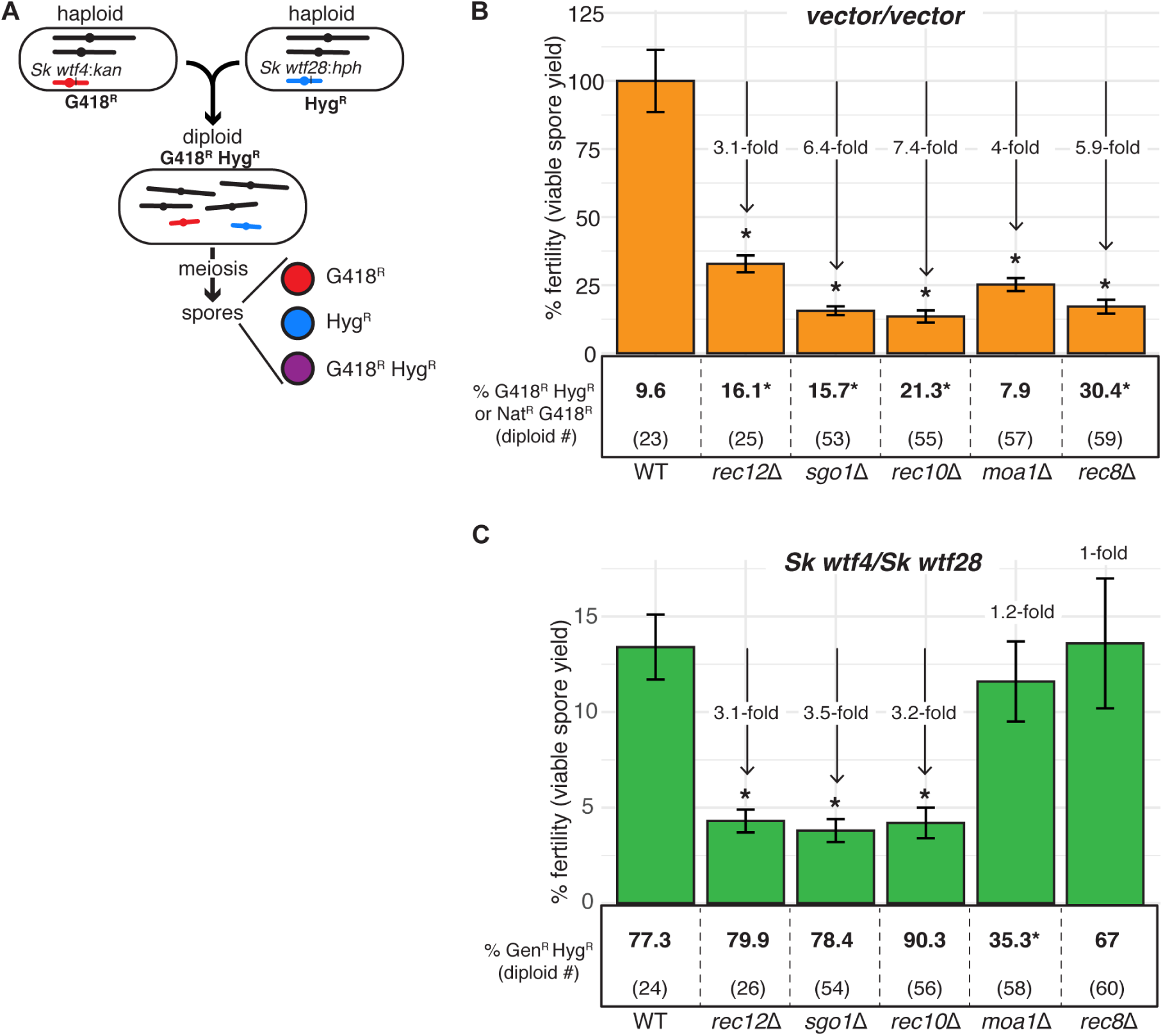
Fitness costs of some meiotic mutants is reduced in diploids with competing *wtf* meiotic drivers relative to a background without *wtf* competition. (A) Schematic of diploid heterozygous for the Hyg^R^ and G418^R^ markers at the *ade6* locus. Fertility measured by viable spore yield in (B) an inbreeding scenario (empty vector/empty vector) or (C) an outcrossing scenario with one set of heterozygous drivers (*Sk wtf28/Sk wtf4*) in wild-type, *rec12*Δ, *sgo1*Δ, *rec10*Δ, *moa1*Δ, and *rec8*Δ homozygous backgrounds. Underneath the bar graph is the % of G418^R^ Hyg^R^ (aneuploid, diploid, or haploid with duplication event) progeny for each diploid. In the *sgo1* mutant for the vector/vector diploid, we used the frequency of Nat^R^ G418^R^ progeny, instead of G418^R^ Hyg^R^. Error bars represent the standard error of the mean. * indicates p-value <0.05 (G-test [G418^R^ Hyg^R^, Nat^R^ G418^R^] and Wilcoxon test [fertility]). For statistical analyses of the frequency of G418^R^ Hyg^R^ (or Nat^R^ G418^R^) progeny and fertility, we compared diploids 25, 53, 55, 57, and 59 to control diploid 23, and diploids 26, 54, 56, 58, and 60 to control diploid 24. At least three, but usually more, independent diploids were used to calculate viable spore yield. Additionally, more than 200 viable spores were scored for each diploid. The data for diploid 23 were previously published in Bravo Núñez et al., 2020. Data for diploids 23-26 are also shown in Figure 4 and Figure 6. Raw data are shown in Figure 6—figure supplement 3 and Figure 6—figure supplement 4.

**Figure 6—figure supplement 2.**
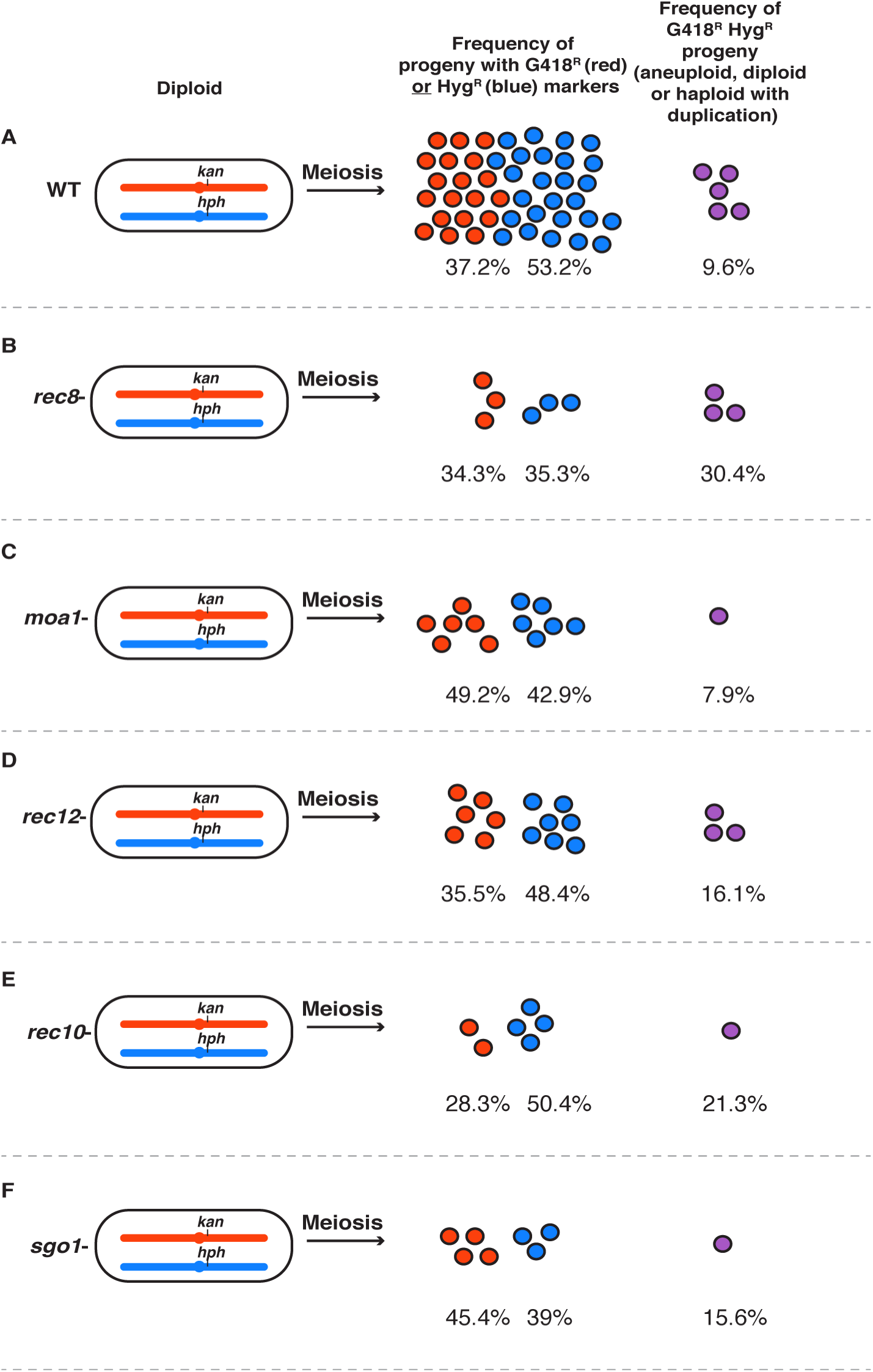
Pictorial description of mutant effects on fertility and the frequency of G418^R^ Hyg^R^ progeny. Summary figure depicting the percentage of haploids and abnormal progeny (aneuploids, diploids, and haploids with duplications of (A) wild type, (B) *rec8*, (C) *moa1*, (D) *rec12*, (E) *rec10,* and (F) *sgo1* based on viable spore yield and allele transmission values. If spore viability was wild type, there are 50 circles. Fewer circles represent a proportional drop in viable spore yield. The raw data used to generate this figure can be found in Figure 6—figure supplement 3 and Figure 6—figure supplement 4.

**Figure 6—figure supplement 3.**
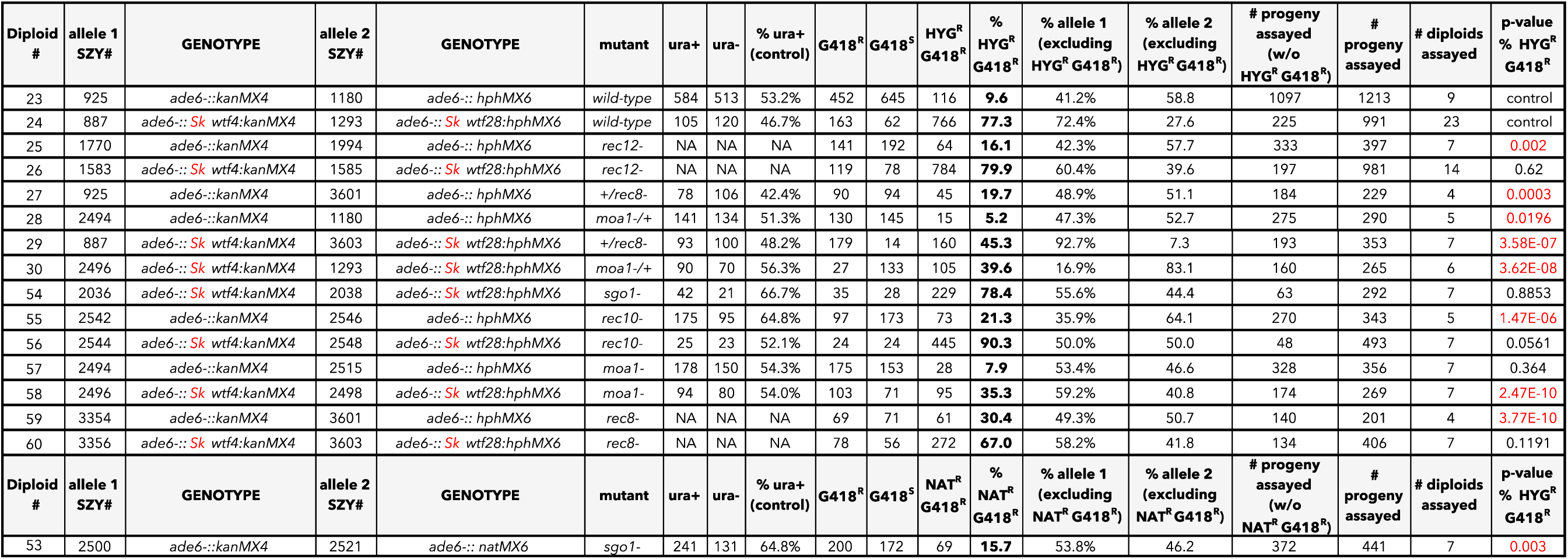
Raw data of allele transmission values reported in Figure 6 and Figure 6—figure supplement 1. Each of the horizontal lines represents the relevant genotype and allele transmission of the indicated diploid. The first column matches the diploid number from Figure 6 and Figure 6—figure supplement 1. In columns 2-5 are the SZY strain number and relevant genotypes used to determine the allele transmission of the *wtf* genes or empty vector. Column 6 indicates if the diploid has a mutant or wild-type genotype. Columns 7 and 8 indicate the indicated genotype (ura+ or ura- progeny) amongst haploids. Columns 10 and 11 indicate the number of haploid progeny that exhibited the indicated phenotype (G418^R^ or G418^S^) excluding G418^R^ Hyg^R^ progeny. Column 12 shows the number of progeny that exhibited the G418^R^ Hyg^R^ phenotype. Column 16 indicates the total number of progeny assayed excluding G418^R^ Hyg^R^ progeny. In diploid 53, we used the frequency of Nat^R^ G418^R^ progeny, instead of G418^R^ Hyg^R^. Column 17 indicates the total number of progeny assayed. Column 18 shows the total number of independent diploids assayed for each cross. The last two columns show the p-values calculated from the allele transmission at *ade6* and G418^R^ Hyg^R^ progeny, respectively. To calculate the p-values using a G-test, we compared diploids 25, 27, 28, 53, 55, 57, and 59 to control diploid 23; diploids 26, 29, 30, 54, 56, 58, and 60 to control diploid 24. For comparison purposes, we included the data for diploids 23-26, which are also shown in Figure 2—figure supplement 5 and Figure 4—figure supplement 5. The data for diploid 23 were previously published in Bravo Núñez et al., 2020.

**Figure 6—figure supplement 4.**
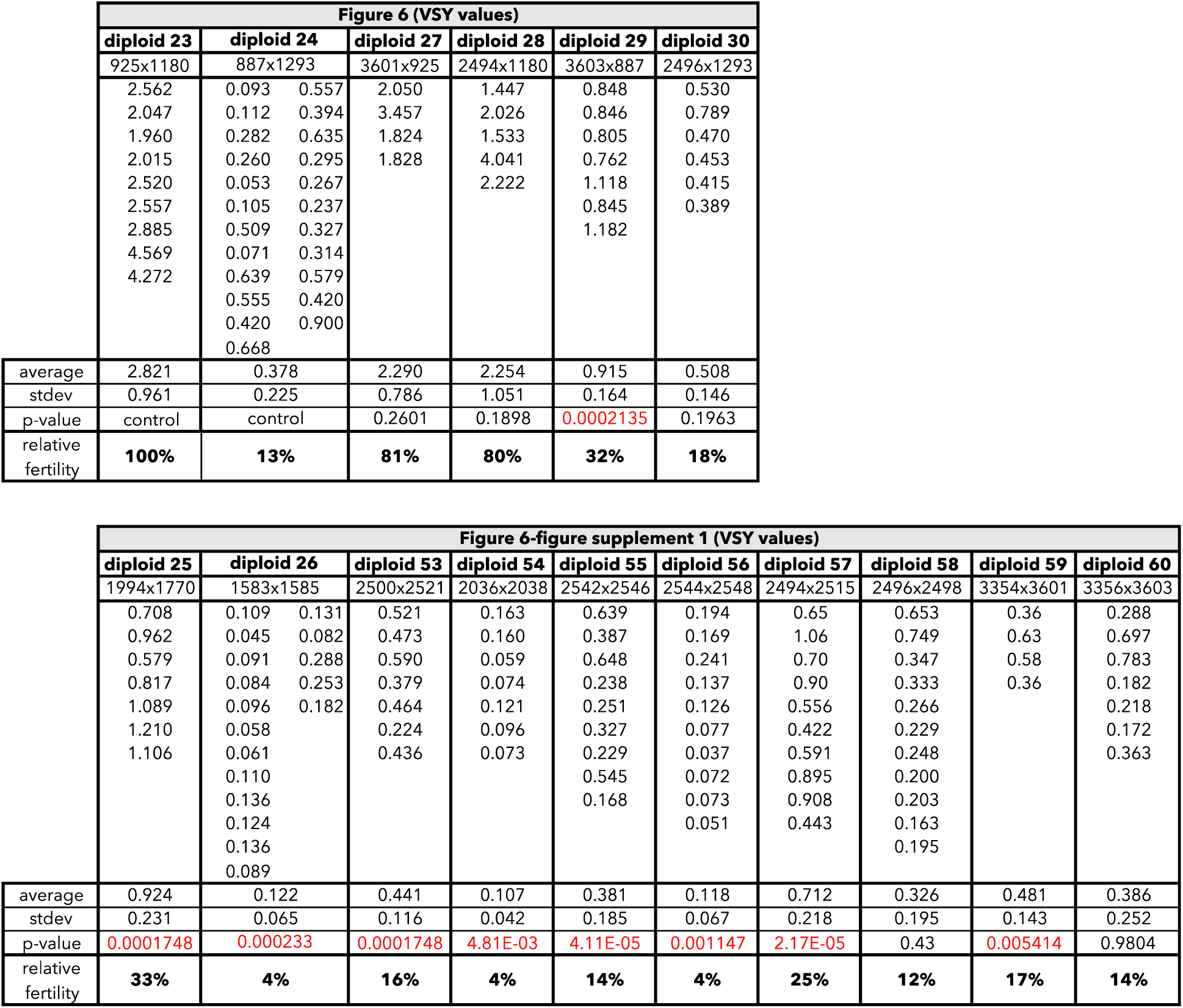
Raw data of the viable spore yield reported in Figure 6 and Figure 6—figure supplement 1. Each column represents the diploid assayed, which matches the diploid number in Figure 6 and Figure 6—figure supplement 1. The diploid number and SZY strain numbers of both haploid parental strains are shown at the top. We present all the viable spore yield values from independent assays. We calculated the p-value using the Wilcoxon test by comparing diploids 25, 27, 28, 53, 55, 57, and 59 to control diploid 23; and diploids 26, 29, 30, 54, 56, 58, and 60 to control diploid 24. For comparison purposes, we included the data for diploids 23-26, which is also shown in Figure 2—figure supplement 4 and Figure 4—figure supplement 6. The data for diploid 23 were previously published in Bravo Núñez et al., 2020.

**Figure 7—figure supplement 1.**
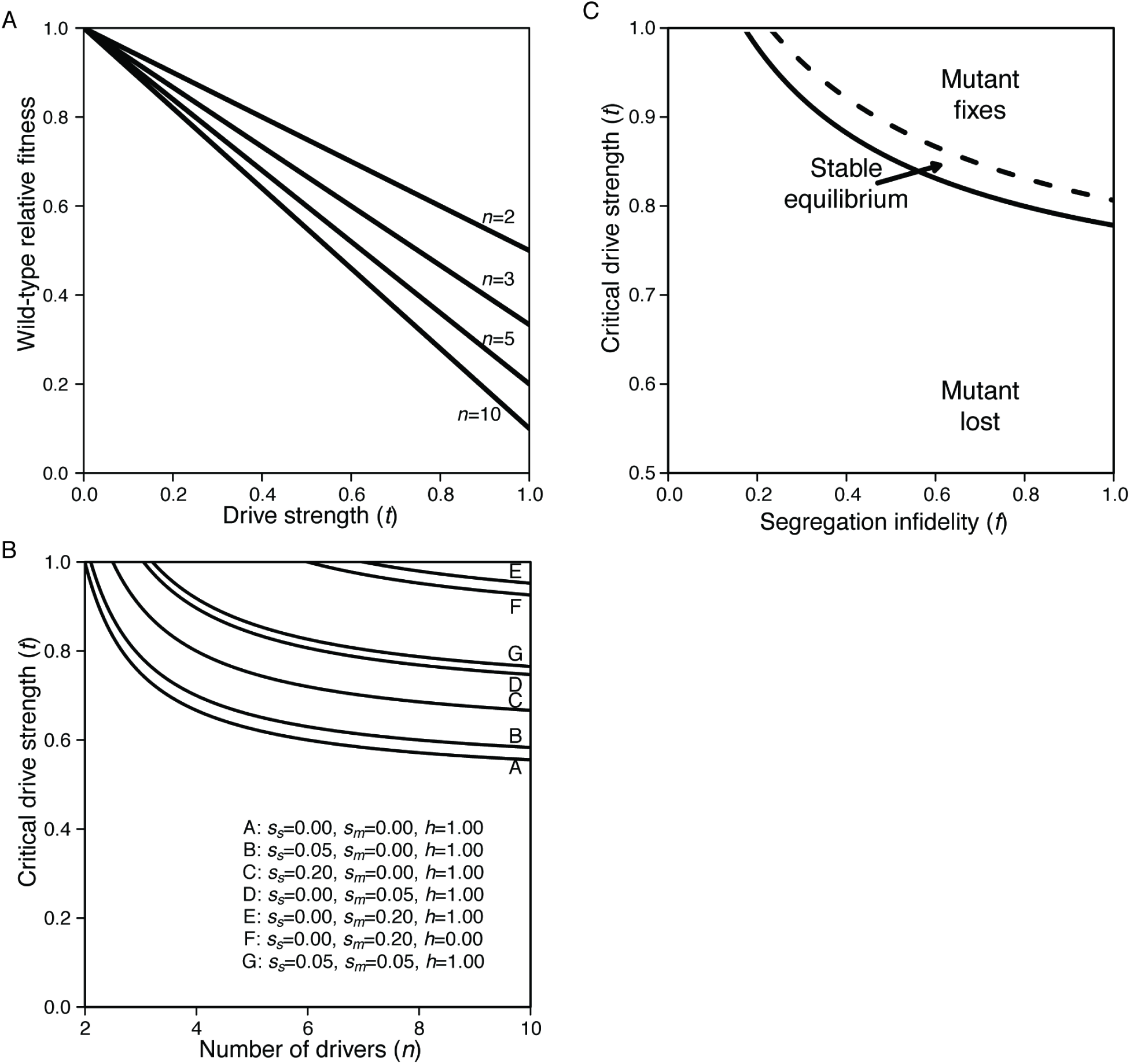
The drive load due to *wtf* segregating within populations. (A) From equation S.1, *n* is the number of drivers in the population, *t* is the strength of drive. Wild-type fitness is relative to a population without any drivers. (B) Meiotic mutant invasion criteria with extrinsic costs. All curves assume *f* = 0.1. Other parameters are listed in the plot. (C) Narrow parameter space allows for stable equilibrium. Solid line represents *t_critical_*, below which the segregation mutant cannot invade. Dashed line represents *t_fixation_*, above which the mutant fixes. In the space between the lines, the equilibrium ranges from zero (solid line) to one (dashed line). *n*=5, *s_m_*=0.2, *s_s_*=0.1, *h*=1.

**Figure 8—figure supplement 1.**
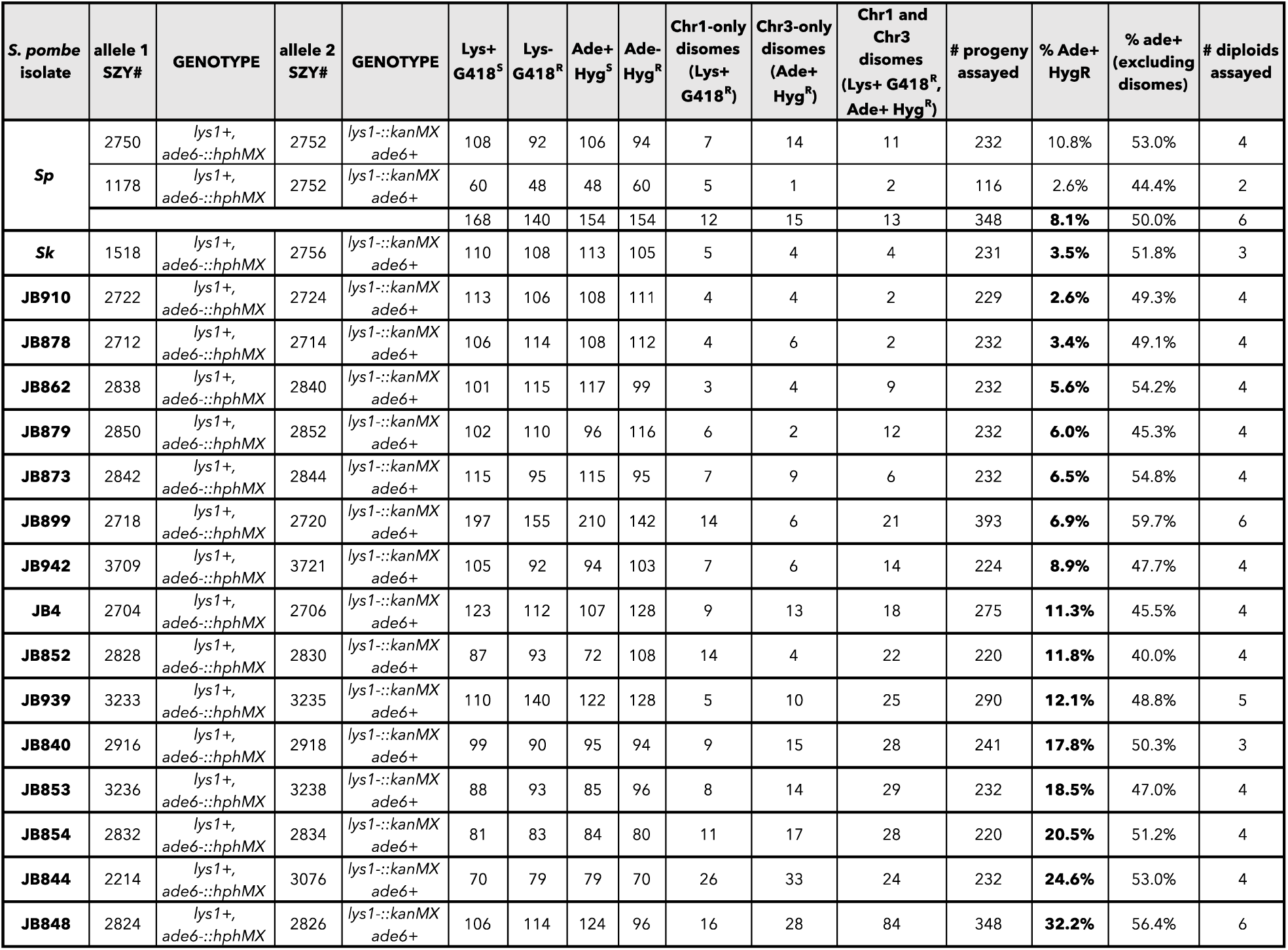
Raw data of allele transmission values reported in Figure 8. Each of the horizontal lines represents the relevant genotype and allele transmission of the indicated diploid. The first column shows the *S. pombe* diploid assayed. In columns 2-5 are the SZY strain number and relevant genotypes used to determine the allele transmission for chromosome 1 and chromosome 3. Columns 6-12 indicate the number of progeny that exhibited the indicated phenotypes. The total number of progeny assayed is shown in column 13. Column 14 indicates the percentage of the progeny that were likely disomes (Ade+ and Hyg^R^). Column 15 indicates the percentage of the progeny (shown in column 8) that were Ade+ Hyg^S^. Ade+ Hyg^R^ progeny were excluded to calculate these percentages. The last column shows the total number of independent diploids assayed for each cross.

## Supplementary File 1. Population genetics model

It is counterintuitive that a mutation causing reduced fidelity of chromosome segregation during meiosis could ever be beneficial and spread through a population. To determine whether such a segregation infidelity mutation could invade a population with *wtf-*like meiotic drivers, we created a simple mathematical model of the scenario.

### The model

As described in the main text, we start with a random mating population of infinite size with *n* distinct drivers – all at the same locus on the same homologous chromosome. Each driving chromosome has the same strength of drive (*t*) and is completely susceptible to all other driving chromosomes (the antidote of the 3^rd^ driver does not protect against the poison of the 2^nd^ driver). We assume that each drive chromosome is at equal frequency so the frequency of the *i*^th^ driver is *P_i_* = 1/*n*. This means that there are no chromosomes without a driver in the population – a reasonable assumption given observations in isolates of *S. pombe* (Bravo Núñez et al., 2018; Bravo Núñez et al., 2020; Eickbush et al., 2018; Hu et al., 2017; Nuckolls et al., 2017). With random mating, the proportion of the population that are homozygotes when it comes to the drive chromosome is 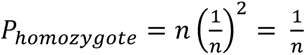 and the proportion that are heterozygotes is 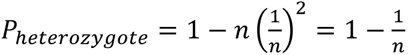 . Since all chromosomes contain *wtf* drive loci, all heterozygotes are likely to produce gametes that will be killed by the *wtf^poison^*.

The strength of drive (*t*) ranges from zero (*wtf^poison^* does not kill any gametes) to one (*wtf^poison^* kills all gametes that do not produce the specific *wtf^antidote^*). Therefore, the relative number of gametes produced by a heterozygous parent is 1-*t* and the average relative fitness of a population with *n* drivers all at equal frequency (*P_i_*) and strength of drive (*t*) is:

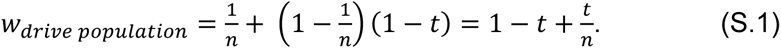

This means that a population with *n* drivers is, on average, *t+t/n* less fit than a population without drivers (Figure 7—figure supplement 1A). This “drive load” is quite severe with large *t* and large *n*. Based on our findings, natural populations show drive strength around *t=*0.9 and segregate for *n≈*5 drivers (this number varies considerably), suggesting the population fitness is 0.28 relative to a population without drive (Bravo Núñez et al., 2020; Eickbush et al., 2018; Hu et al., 2017). Such a burden would select for resistance mechanisms that might include positive assortative mating or inbreeding, resistance alleles such as *wtf18-2* or aneuploidy (Bravo Núñez et al., 2018). With aneuploidy, some fraction of gametes (*f*/2, see below) would inherit two drive chromosomes with both poisons but also both antidotes and therefore would be protected. However, an equal fraction (*f*/2) would receive zero copies of chromosome 3 and would therefore be inviable (Niwa et al., 2006).

We assume that a single locus controls segregation fidelity and that a mutation at that locus (termed a segregation infidelity mutant) results in an increase in aneuploid gametes due to a defect in meiosis I. We also assume that this mutation only influences segregation on chromosome 3 (where most *wtf* drivers reside) but is not on the third chromosome itself (Eickbush et al., 2018; Hu et al., 2017). However, reduced segregation fidelity of other chromosomes can be included in costs discussed below. The parameter *f* denotes the segregation infidelity and can range from 1 (all meioses result in 50% of gametes being disomic and 50% without a third chromosome) to 0 (all gametes are haploid). This segregation infidelity mutant, therefore, has an intrinsic fitness cost since 50% of gametes lack a third chromosome and are therefore inviable. The recursion for the frequency of the infidelity mutant is:

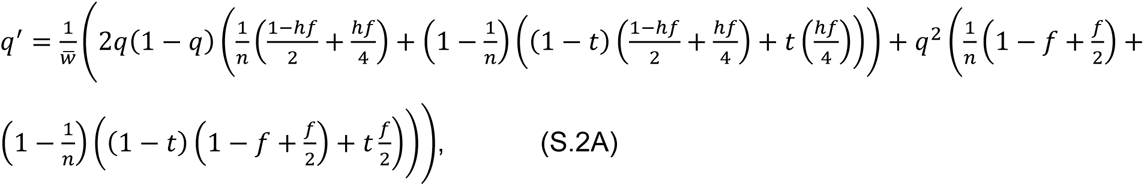

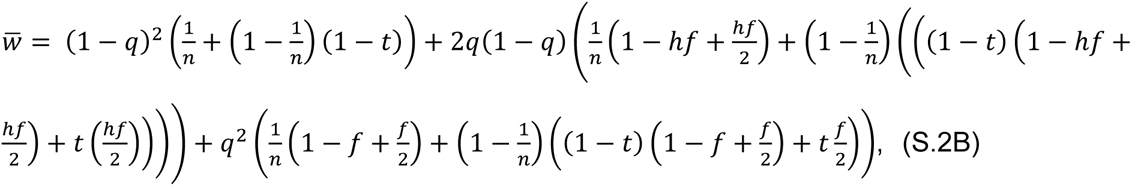

### Results

#### Invasion

For the mutant to invade, we must have *q*′> *q* (the frequency of the mutation in the next generation must be higher than in the current generation). Solving for *q*′ > *q* in terms of *t*, we find:

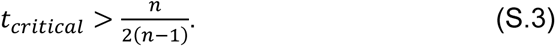

Surprisingly, the critical value of *t* allowing for invasion is independent of the degree of segregation infidelity (*f*) (Figure 7B).

In an attempt to get a more intuitive feel for the path a segregation infidelity mutant might take, we plotted trajectories of such mutants using variable parameters. Figure 7C shows that changes in some parameters influence the trajectories in predictable ways: a) increasing segregation infidelity from *f*=0.1 to *f*=0.9 allows for a more rapid spread; b) dominant mutations (*h*=1) spread more rapidly than recessive ones (*h*=0); c) more segregating drivers (*n*=5) leads to a more rapid spread than fewer segregating drivers (*n*=3); and d) lower strength of drive (*t*=0.55 compared to *t*=0.95) can lead to the loss of the infidelity mutant as predicted by equation S.3 and Figure 7B. Note that in this no-cost model, there is no stable internal equilibrium. All infidelity mutants are either lost or fixed. Accounting for additional costs of the infidelity mutant leads to the possibility of a stable equilibrium, though the parameters allowing this equilibrium are narrow (Figure 7-figure supplement 1C).

#### Allowing for additional costs of segregation infidelity mutants

Segregation infidelity mutants incur an intrinsic fitness cost since 50% of the gametes do not contain a chromosome but may also carry additional costs. We imagine two such costs. First, we allow for additional fitness costs (*s_s_*) experienced by disomic spores. There is ample evidence that spores disomic for the third chromosome jettison one copy relatively quickly (Niwa et al., 2006), but there may be some initial cost due to gene dosage imbalance or other factors. Second, we can imagine that the meiotic mutant might also bear a cost because it leads to other problems including missegregation of the other chromosomes. We denote this cost *s_m_* and it has an associated dominance (*h*). The dominance coefficient (*h*) is assumed to apply to both the cost of the segregation mutant (*s_m_*) and the segregation infidelity (*f*). The recursion for the frequency of the infidelity mutant is therefore:

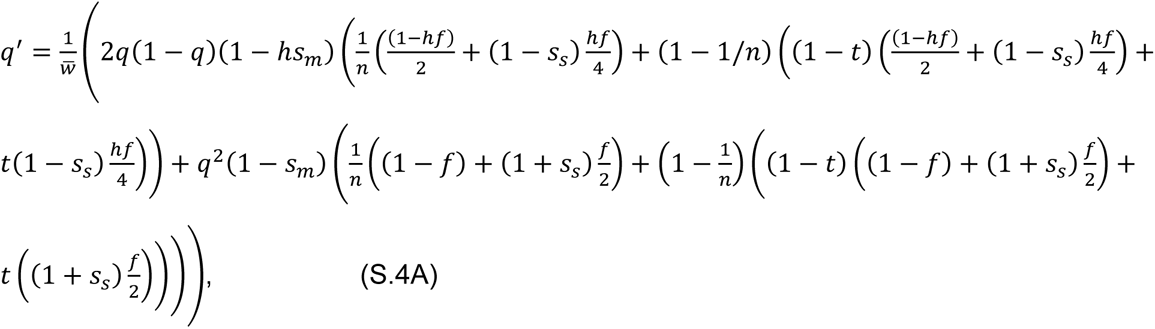

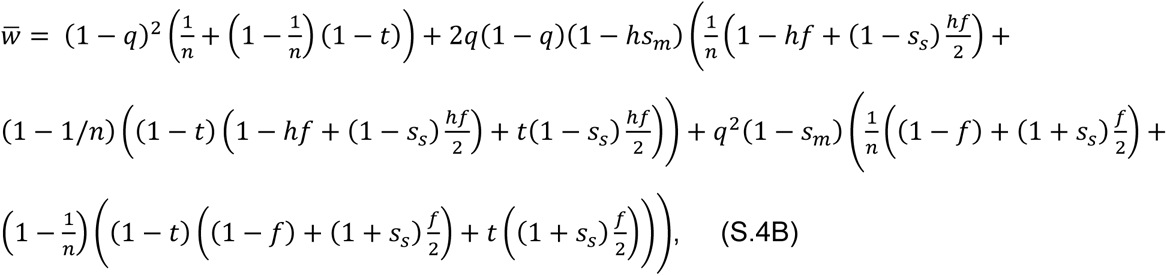

#### Invasion with additional costs

For the mutant to invade, we again must have *q*′ > *q*. Solving for *q′* > *q* in terms of *t*, we find:

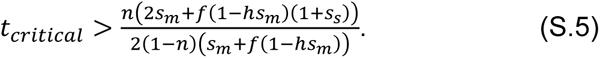

If we assume that there is *no* cost of being a disomic spore (*s_s_*=0), this becomes:

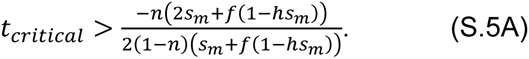

If we instead assume that there is *no* cost of carrying the meiotic mutation (*s_m_*=0), this becomes:

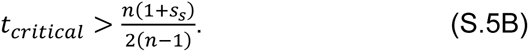

Figure 7—figure supplement 1B shows the effect of these costs on the critical value of drive strength required for a meiotic infidelity mutant to invade. Note that even with high cost (20%), strong drivers select for these meiotic infidelity mutants. In addition to reducing the parameter space allowing for invasion of segregation infidelity mutants, these additional costs also slow the invasion of the mutant. For example, a mutant without extrinsic cost (*s_m_* and *s_s_* are 0) takes 74 generations to climb from a frequency of 1% to 50% (*f*=0.33, *t*=0.9, *h*=1; *n*=5). With low cost (*s_m_*=0.05) this would take 25 generations and with higher cost (*s_m_*=0.20) this would take 755 generations. Note that with higher costs (e.g. *s_m_*=0.3), the mutant does not invade because our *t*=0.9 is less than the critical *t*=0.978 for invasion. However, with observed parameters in various isolates of *S. pombe* (*t*=0.9 and *n*=5), the system could tolerate considerable cost and still allow mutants to invade.

#### Equilibrium frequency of meiotic mutants with costs

Extrinsic costs of segregation infidelity mutants also allow for a narrow range of stable equilibria in some cases. Solving *q*^′^ = *q* for *q* provides the equilibrium values. There are three such values: 0, 1 and

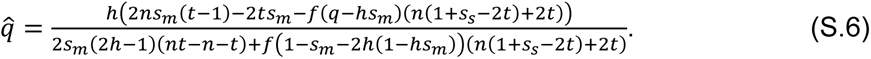

The value in equation S.6 is between zero and one for only a narrow range of parameter values. We can think of this in terms of a range of *t* allowing for a stable equilibrium. Below *t_critical_*, the mutation is lost. However, if *t* is high enough, the meiotic mutation will fix. This happens when *q* = 1, which is when

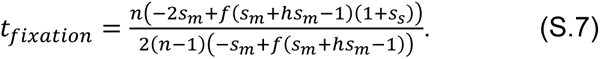

Figure 7—figure supplement 1C shows the narrow range allowing for stable equilibrium. Note that in the narrow space between the solid line for *t_critical_* and the dashed line for *t_fixation_*, the stable equilibrium frequency ranges from zero to one.

#### Limitations and caveats

Our model is simplistic in several ways but does suggest that a *wtf-*like poison-antidote system might select for mutations that disrupt “normal” chromosome segregation fidelity. The first caveat is that we assume the population is randomly mating. However, due to spatial proximity of cells, we expect that inbreeding may be common. Such inbreeding would reduce the proportion of matings between haploids with different drive chromosomes and therefore decrease the benefit of segregation infidelity. A second caveat is that we assume all drive strengths are equal and all drivers are at equal frequency in the population. This is a logical starting point but is unlikely to be the case in real populations. Increasing the variance of the frequencies of individual drive chromosomes within a single population will also reduce the proportion of matings between haploids with different drive chromosomes and will decrease the benefit of segregation infidelity. Finally, we do not allow for non-driving third chromosomes. Diploids with a wild-type and a driving chromosome would still benefit from segregation infidelity. This benefit would be due to the disomic spores generated, in which both poison and antidote would be produced, and thus the disomic spore would be saved. However, selection for the infidelity mutation would be weaker because the drive load in the population would be less since fewer gametes are killed each generation due to drive.

## Supplementary File 2. Yeast strains used in this study

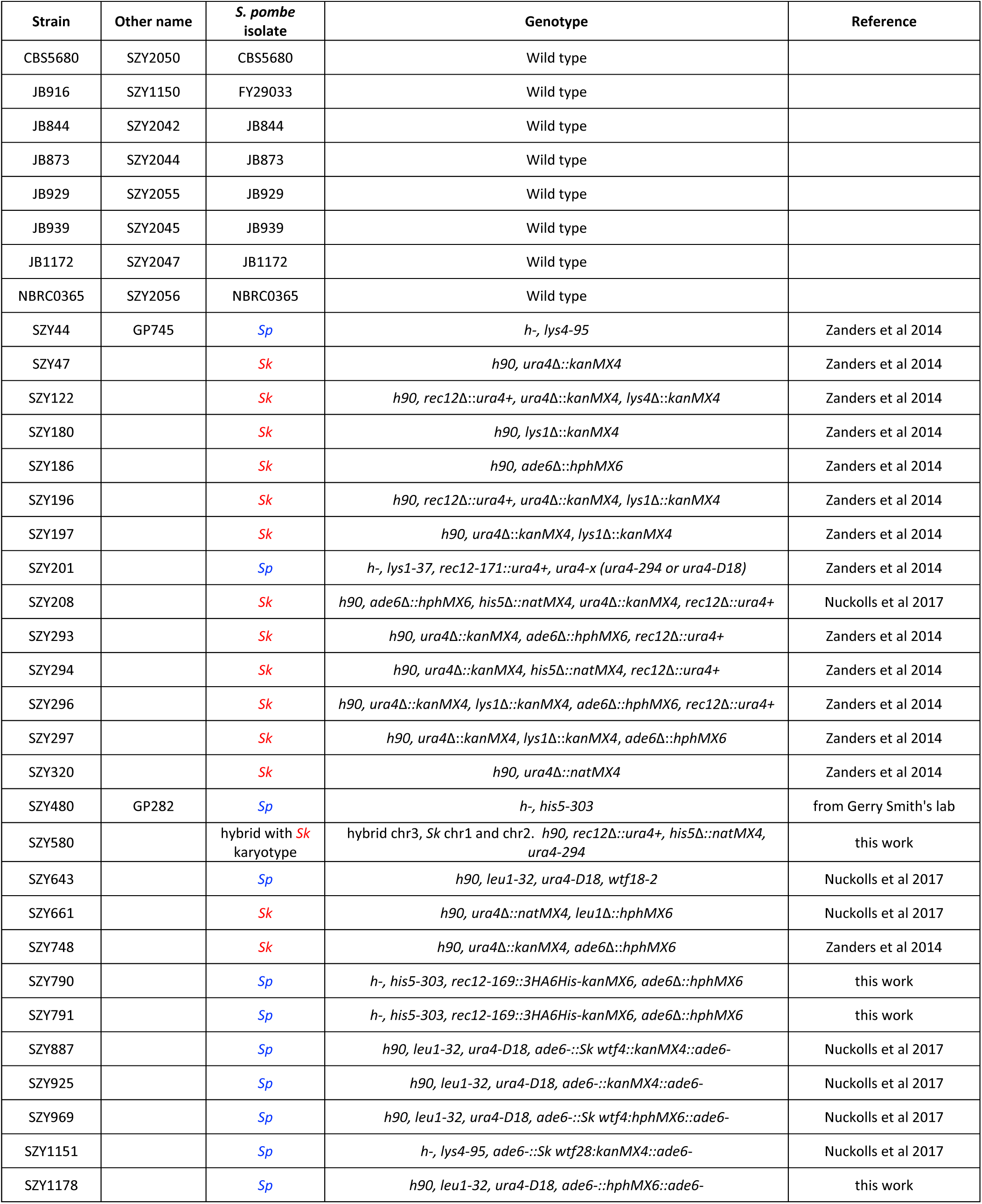
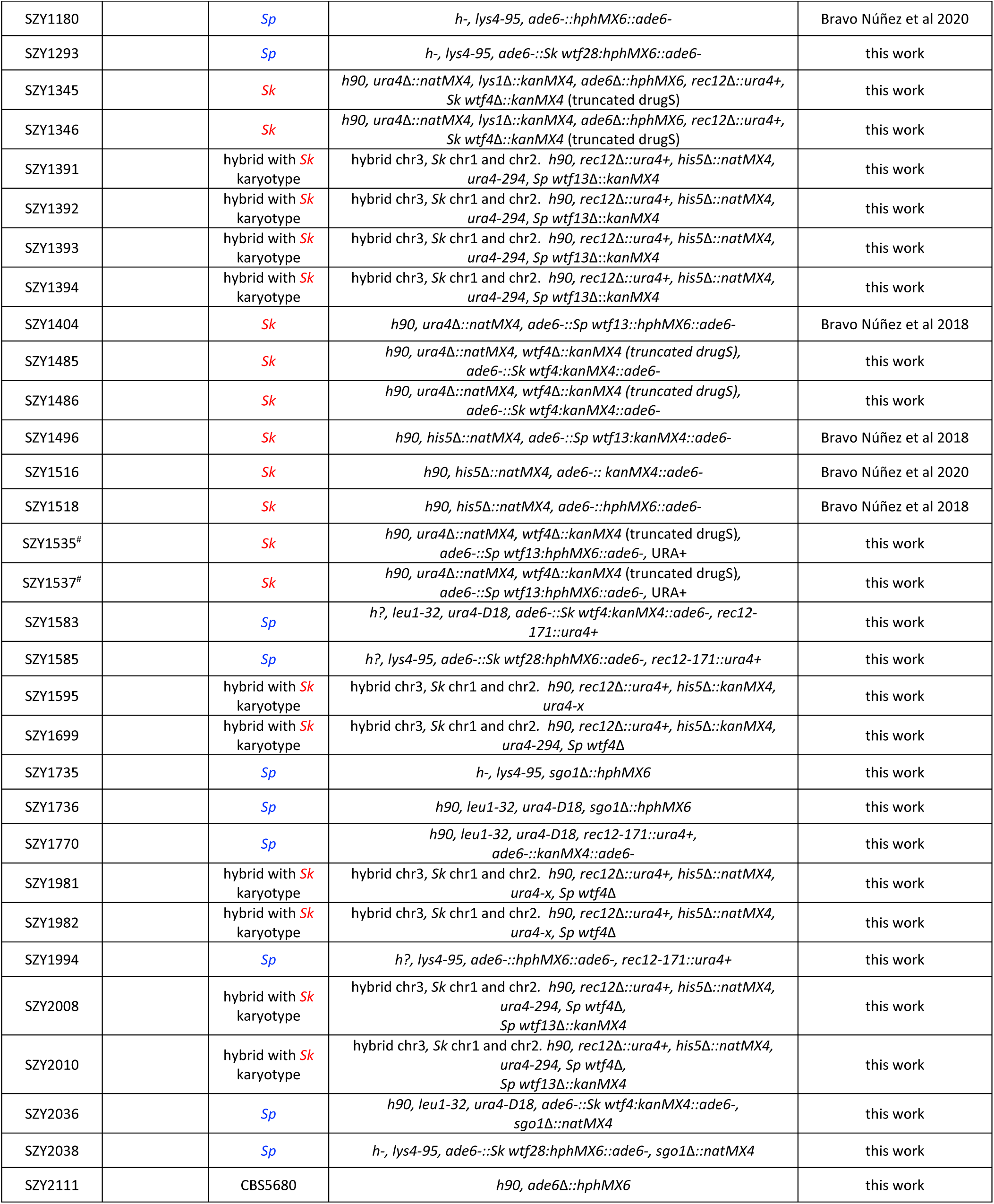
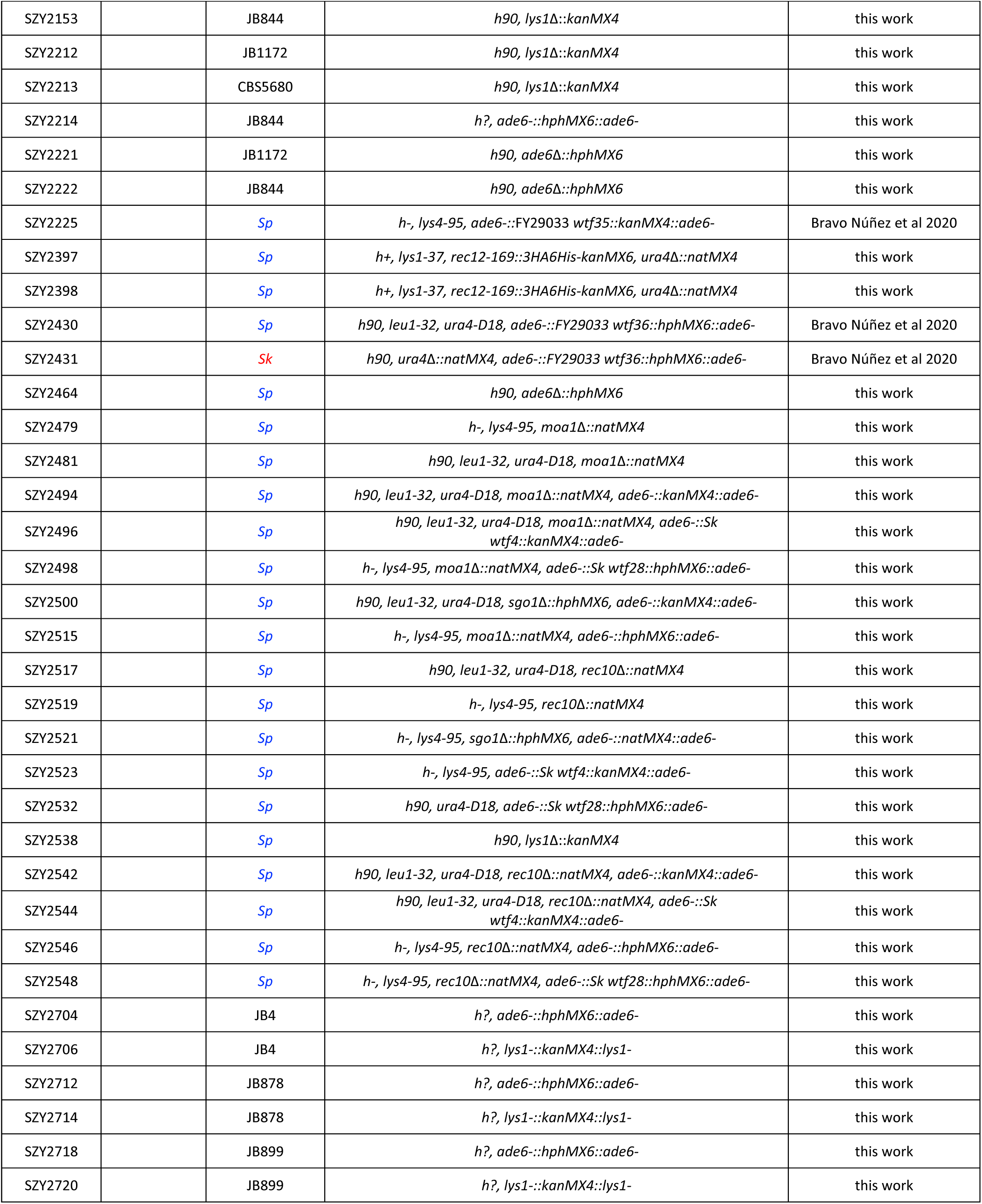
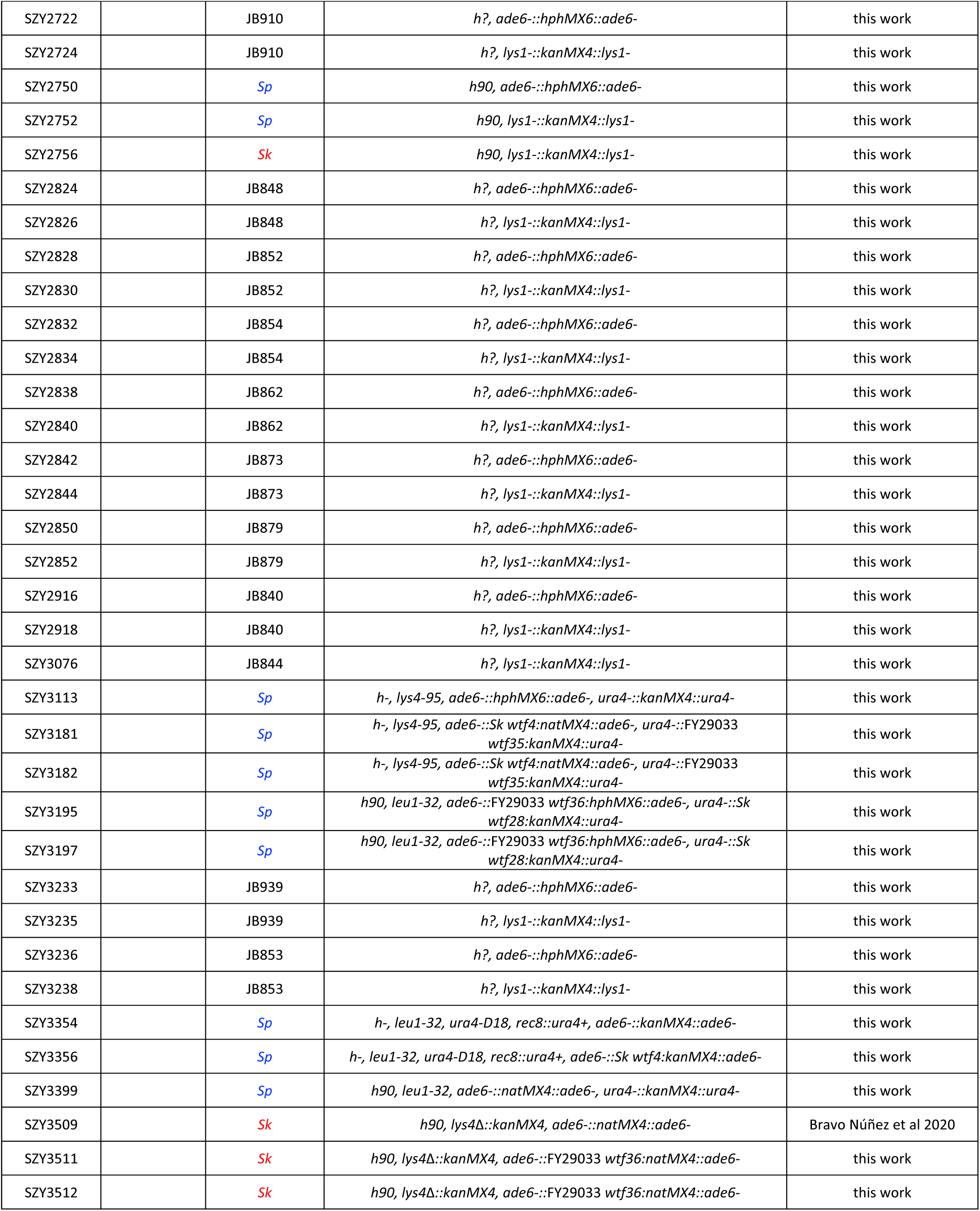
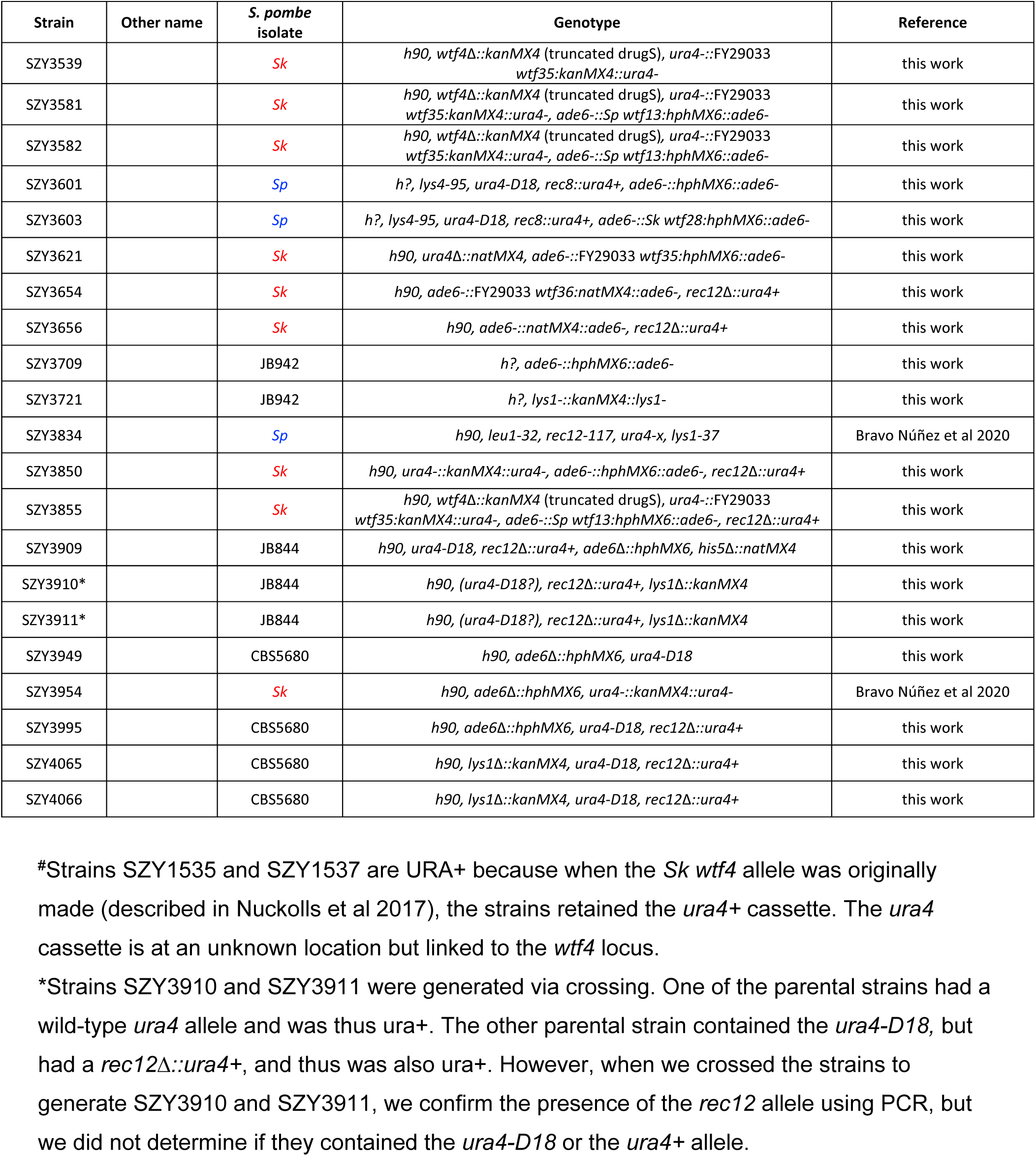

## Supplementary File 3. Plasmids used in this study

Supplementary File 3. Plasmids used in this study.
| Plasmids | short description | reference |
| --- | --- | --- |
| pFA6 | contains kanMX4 | Wach et al 1994 |
| pAG25 | contains natMX4 | Goldstein and McCusker 1999 |
| pAG32 | contains hphMX6 | Goldstein and McCusker 1999 |
| pMZ379 | contains Cas9 | Rodríguez López et al 2016 |
| pSZB188 | derivative of pFA6 that integrates at *ade6*, yielding *ade6*- | Nuckolls et al 2017 |
| pSZB189 | pSZB188 with *Sk wtf4* cloned into SacI site | Nuckolls et al 2017 |
| pSZB254 | pSZB188 with *Sk wtf28* into SacI site | Nuckolls et al 2017 |
| pSZB331 | derivative of pFA6 that integrates at *ura4*, yielding *ura*- | Bravo Núñez et al 2020 |
| pSZB332 | derivative of pFA6 that integrates at *ura4*, yielding *ura*- | Bravo Núñez et al 2020 |
| pSZB386 | derivative of pAG32 that integrates at *ade6*, yielding *ade*- | Bravo Núñez et al 2018 |
| pSZB412 | pSZB386 with *Sk wtf28* cloned into SacI site | this work |
| pSZB570 | derivative of pMZ379 containing the *Sp wtf4* gRNA | this work |
| pSZB718 | pSZB332 with *Sk wtf28* cloned into SacI site | this work |
| pSZB722 | derivative of pAG25 that integrates at *ade6*, yielding *ade*- | Bravo Núñez et al 2020 |
| pSZB723 | derivative of pAG25 that integrates at *ade6*, yielding *ade*- | Bravo Núñez et al 2020 |
| pSZB788 | pSZB188 with FY29033 *wtf35* cloned into SacI site | Bravo Núñez et al 2020 |
| pSZB816 | derivative of pFA6 that integrates at *lys1*, yielding *lys*- | this work |
| pSZB849 | derivative of pAG25 that integrates at *ade6*, yielding *ade*- | this work |
| pSZB923 | pSZB849 with *Sk wtf4* into SacI site | this work |
| pSZB1001 | pSZB331 with FY29033 *wtf35* into SacI site | this work |
| pSZB1060 | pSZB849 with FY29033 *wtf36* into SacI site | this work |

## Supplementary File 4. Table summary of plasmid construction

**Supplementary File 4. Table summary of plasmid construction.**
| *wtf* allele | *S. pombe* isolate | DNA template | PCR oligos used | PCR fragment digested with restriction enzyme | target site | Resulting plasmid number |
| --- | --- | --- | --- | --- | --- | --- |
| *wtf28* | *Sk* | pSZB254 | 733+651 | SacI | SacI site of pSZB386 | pSZB412 |
| *wtf28* | *Sk* | SZY661 | 733+651 | SacI | SacI site of pSZB332 | pSZB718 |
| *wtf35* | FY29033 | pSZB788 | 1036+1349 | SacI | SacI site of pSZB331 | pSZB1001 |
| *wtf36* | FY29033 | SZY1150 | 1351+1593 | SacI | SacI site of pSZB849 | pSZB1060 |

Column 1 lists the *wtf* gene cloned into each vector. Column 2 denotes the isolate from which each *wtf* was cloned. The DNA templates and oligos used in the PCR reactions to amplify the *wtf* alleles are shown in columns 3 and 4, respectively. We digested each of the amplified fragments with the enzymes reported in column 5 and then integrated into the target site listed in column 6. The strain number of each of the plasmids that we generated is reported in column 7. The description of each plasmid can be found in Supplementary File 3.

## Supplementary File 5. Oligo table

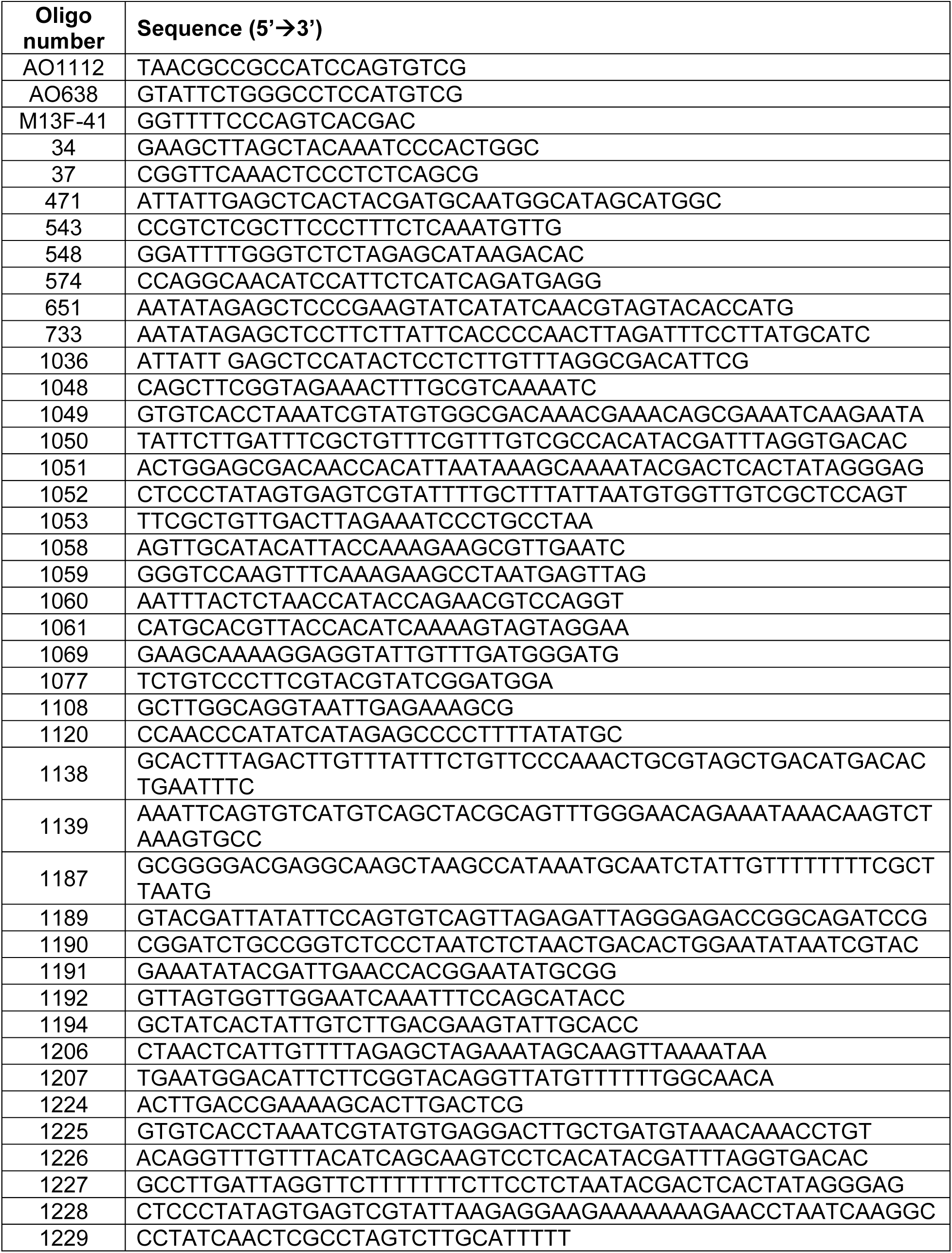
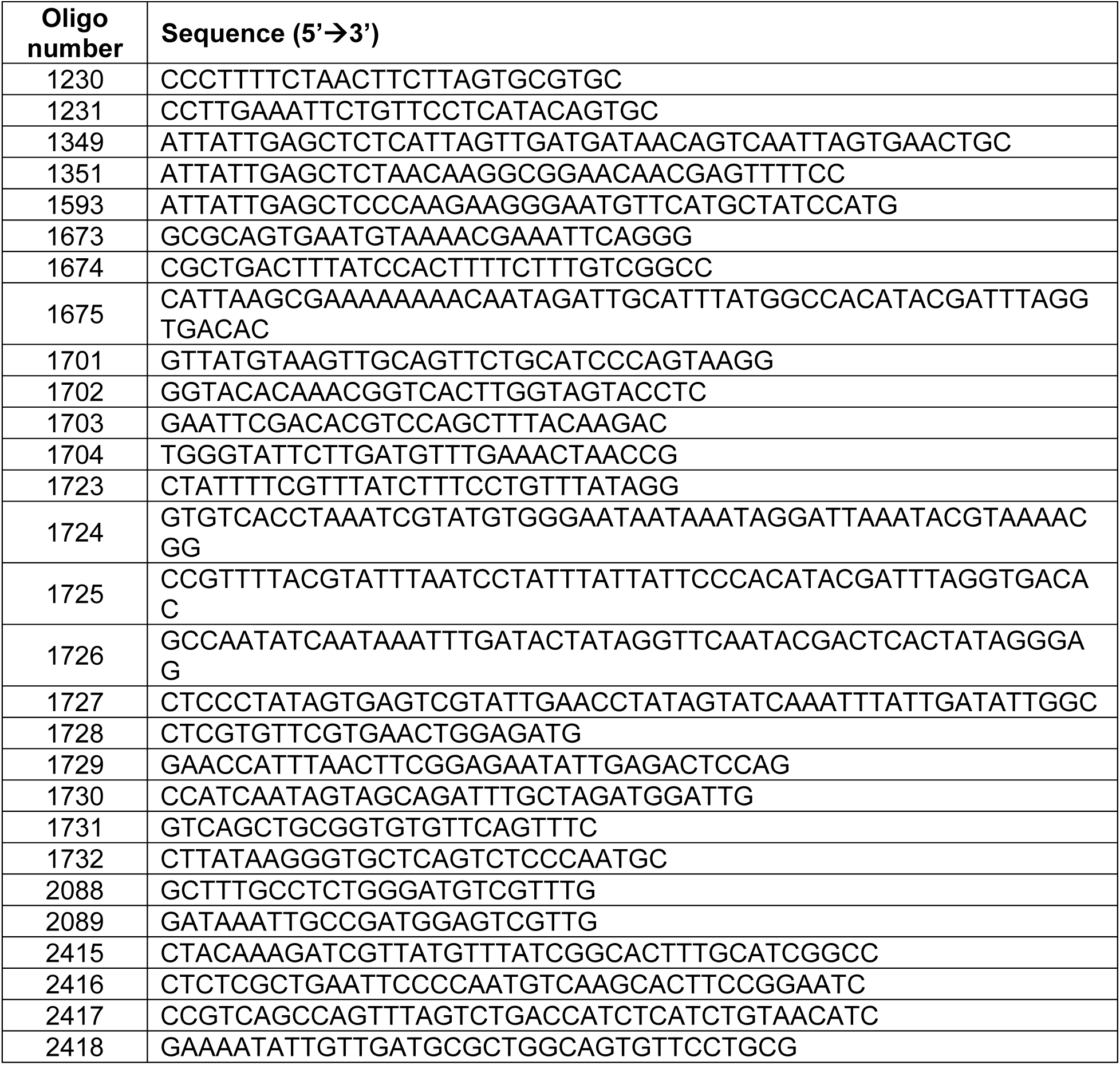

